# Motor cortex flexibly deploys a high-dimensional repertoire of subskills

**DOI:** 10.1101/2025.09.07.674717

**Authors:** Elom A. Amematsro, Eric M. Trautmann, Najja J. Marshall, L.F. Abbott, Michael N. Shadlen, Daniel M. Wolpert, Mark M. Churchland

**Author notes:** Equal contribution (Elom A. Amematsro, Eric M. Trautmann).

## Abstract

Skilled movement often requires flexibly combining multiple subskills, each requiring dedicated control strategies and underlying computations. How the motor system achieves such versatility remains unclear. Using high-density Neuropixels recordings from primary motor cortex (M1) in macaques performing a challenging force-tracking task, we reveal that M1 activity is much higher-dimensional, and far more flexible, than traditionally assumed. Although our task employed only a single external degree of freedom, neural dynamics reflected transitions amongst many dimensions and multiple distinct computations. Different behavioral control strategies were associated with distinct neural locations and dimensions, sometimes used compositionally. Groups of population-level factors became active when a particular form of dynamics was needed, and remained silent otherwise. Neural activity was thus dominated by the engaged subskill, and could be very different even for matched motor output. These findings challenge prevailing views of M1, and reveal an unexpectedly flexible and high-dimensional neural system underlying skilled motor behavior.

## Introduction

Competing in Indonesia, the professional surfer Stephanie Gilmore scored a perfect ten by linking bottom turns, off-the-lip maneuvers and tube-riding into a single fluid ride. Her accomplishment illustrates that motor control often requires composition: in her case, flexibly switching, and even simultaneously blending, multiple surfing subskills. Might repertoires of subskill-specific motor computations ^1^ be instantiated and flexibly deployed by primary motor cortex (M1), the brain area most responsible for skilled voluntary movement? While this ‘flexible-repertoire hypothesis’ is appealing, past and current perspectives suggest the opposite: M1 activity appears constrained by relatively few degrees of freedom ^2^, with a repertoire of available patterns that is fixed and small ^3,4^rather than flexible and large. A fixed computation would accord with M1’s status as a primary cortical area, one synapse from spinal motoneurons. Indeed, a venerable assumption is that task manipulations can disambiguate hypotheses by providing different views of a consistent M1 computation ^5,6^. Alternatively, M1’s apparent inflexibility may reflect experimental limitations: laboratory tasks are rarely designed to engage many subskills, and the scale of neural recordings may have been insufficient to reveal signatures of computational flexibility.

Those potential signatures are suggested by a burgeoning interest in multi-task recurrent networks. In recurrent networks, a dynamical flow-field governs a ‘neural state’ ^7,8^in a state-space where each dimension captures one network factor ^9^. Factors are time-varying activity patterns that form the basis both for network outputs and for ‘internal’ network computations. Consequently, factors also form a basis for single-neuron responses. Multi-task networks leverage multiple forms of internal dynamics, each employing different factors ^10–17^. Flexibility arises from the variety of dynamics and their combinatorial potential. The flexible-repertoire hypothesis proposes M1 may function similarly (Fig. 1A).

**Figure 1.**
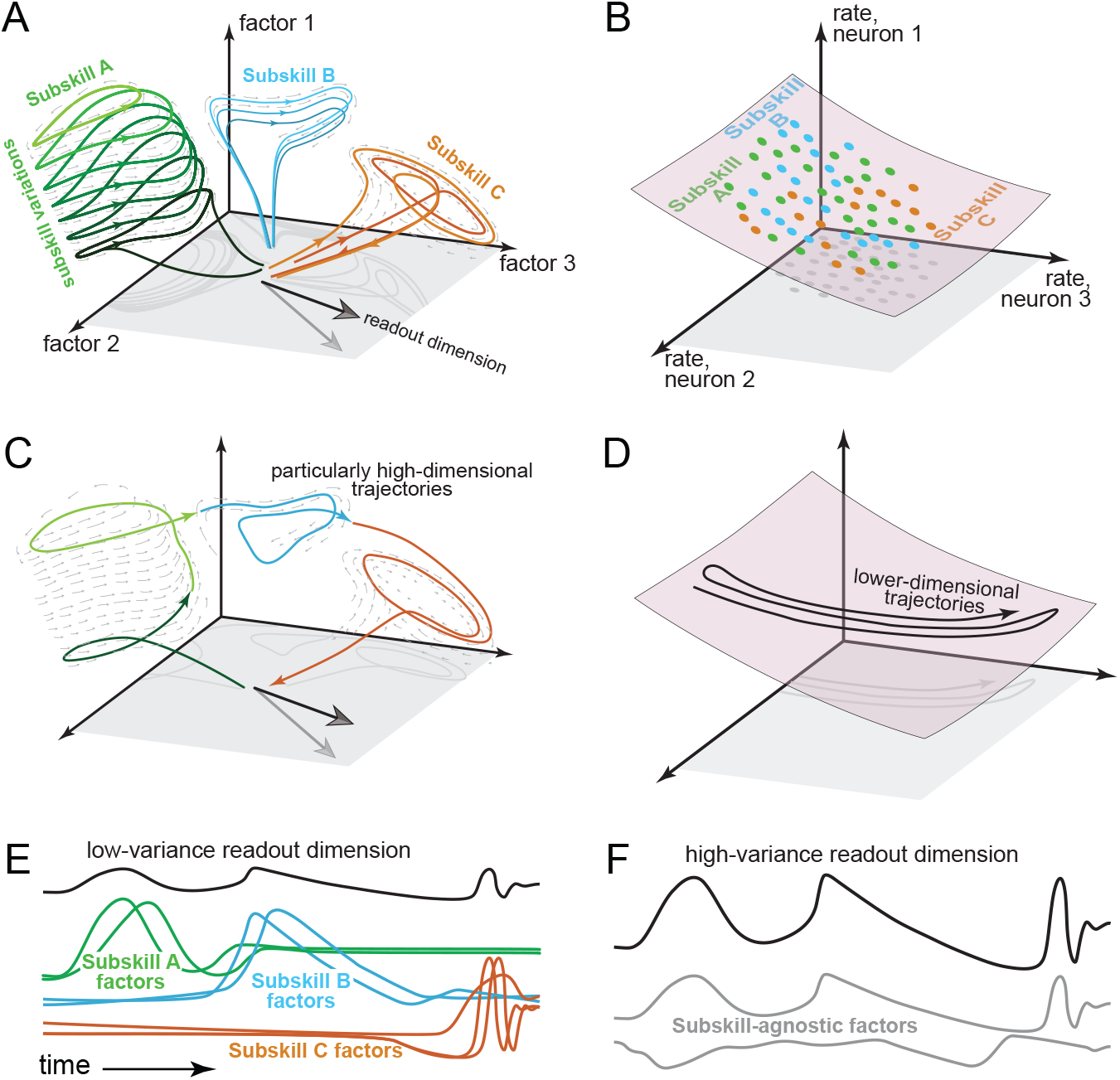
Flexible-repertoire and constrained-manifold predictions. Left column: flexible-repertoire hypothesis; right column: constrained-manifold hypothesis. (**A**) The flexible-repertoire hypothesis predicts high-dimensional activity, resulting from multiple subskill-specific computations that use dedicated locations, dimensions, and dynamics. (**B**) Alternatively, activity occupies a low-dimensional manifold (dots indicate visited states), reused across subskills (colors). (**C**) Flexible-repertoire predictions relevant to the present task. When a task requires multiple subskills, activity should be high-dimensional even if motor output is simple. Some conditions may compositionally reuse subskills, yielding changing locations and dimensions. (**D**) If a task uses one physical dimension and few independent parameters, constrained-manifold activity should become lower-dimensional still. (**E**) Motor output is predicted to rely on different factors at different moments. Consequently, the readout dimension captures little population variance. (**F**) Traditionally, motor readouts derive from a few continuously active high-variance signals.

Like most prevailing conceptions of M1 ^2–4,18^, the flexible-repertoire hypothesis posits dynamics within a state-space ^8,19–21^. Somewhat unusually, the flexible-repertoire hypothesis proposes a very high-dimensional state-space, with different subskills engaging different regions and dimensions (Fig. 1A). Such separation is necessary to isolate different forms of dynamics, thus preventing interference ^22,23^. The potential to combine subskills simultaneously – perhaps on her next wave, Ms. Gilmore blends maneuvers in a surprising new way – requires neural activity that is not restricted to previously observed regions of state-space. These predictions oppose the prevailing view (Fig. 1B) that firing rates are constrained by a low-dimensional manifold ^2,18,24^, preserved and reused across subskills ^3,4^. Contrasting predictions are clearest if a task employs few external degrees of freedom yet engages multiple subskills. Traditionally, eliminating degrees of freedom can only constrain activity further (Fig. 1D). In contrast, under the flexible-repertoire hypothesis, dimensionality is effectively unbounded; it may grow very high if multiple subskills are engaged across conditions or compositionally within-condition (Fig. 1C). Traditionally, readout dimensions target high-variance signals: a few continuously active factors that resemble motor output (Fig. 1F). Under the flexible-repertoire hypothesis, each readout-dimension is impacted by many factors, none consistently resembling the output. Most neural activity is thus output-null ^9,25^and readout-dimensions capture little population variance (Fig. 1E).

Some recent findings potentially align with the flexible-repertoire hypothesis. Learning can alter neural degrees of freedom ^26^, high-dimensional assumptions improve decoding ^27^, and population activity contains numerous ‘small’ (low-variance) signals ^28^. Activity can also traverse different dimensions ^23,29–38^: e.g. when switching from walking to lever-pulling, or switching the arm used to reach. However, these situations involve large changes in muscle-muscle correlations, including which muscles are used. Changes in neural dimensions are thus expected under nearly any hypothesis, including simply because motor-output dimensions change (a common and reasonable interpretation).

An ideal test of the flexible-repertoire hypothesis requires the situation outlined above: a challenging task that engages multiple subskills, yet where simple motor output employs one fixed axis. We leveraged the ‘Pac-Man’ force-tracking task ^28^, which employs one degree of freedom (pressing forward) and is isometric: the arm does not move. Essentially all external variables – including muscle activity and cursor position – mirror force or its derivative. Yet accurate performance demands various combinations of reactive (closed-loop) and proactive (open-loop) strategies. The flexible-repertoire hypothesis predicts (i) activity will be high-dimensional despite physical task simplicity, (ii) the neural manifold will not be preserved: activity will occupy different regions and dimensions across subskills, (iii) some conditions will use multiple subskills compositionally, and (iv) each subskill will be reused across some conditions but not others. A final strong prediction (v) is that neural activity should differ when subskills differ, even if motor output is identical.

An ∼100-neuron population is inadequate to resolve activity that is predicted to be diverse, temporally rich, and high-dimensional. We thus employed primate-optimized Neuropixels Probes ^39^ to record large (∼1000) populations. To aid interpretation via comparison with a downstream neural population, intra-muscular recordings isolated single motor units, each mirroring the spikes of one spinal motoneuron. Motoneuron activity was simple and low-dimensional. M1 activity was high-dimensional and obeyed flexible-repertoire predictions. Thus, the computational role of M1 appears far more flexible than typically envisioned. When one considers the repertoire of abilities acquired in our lifetimes, M1 dimensionality and the variety of dynamics it instantiates are presumably vast.

## Results

### A challenging force-tracking task evokes multiple behavioral strategies

Two rhesus macaques (monkeys C and I) performed an isometric force-tracking task ^28^. The height of a Pac-Man icon mirrored forward force applied to an immovable handle (Fig. 2A, Supp. Movie 1). The task required Pac-Man’s height to be continuously adjusted to intercept a leftward-scrolling dot-path. Dot-paths specified 12 (monkey C) and 11 (monkey I) distinct target force profiles (i.e. conditions, Fig. 2B).

**Figure 2.**
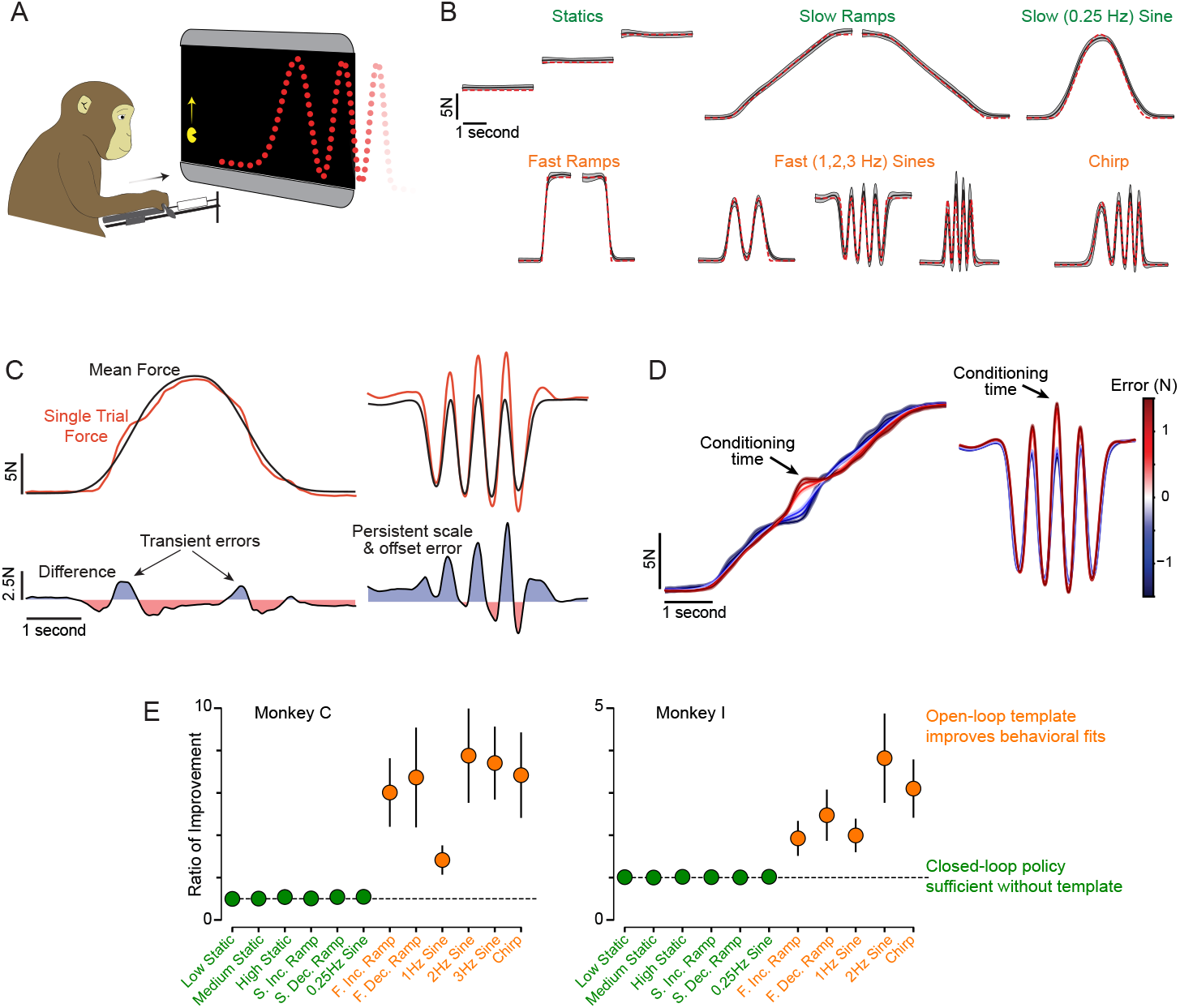
Task and behavior. (**A**) Task schematic. The leftward-scrolling dot-path revealed ∼1 second of future target force. (**B**) Target (dashed red) and trial-averaged forces (black). Gray envelopes: across-trial SD. Data for Monkey C (Monkey I; Supp. Fig. 1). (**C**) Example single-trial forces (red), mean force (black) and error (their difference, bottom) during 0.25 and 2 Hz sines. (**D**) Trial-averaged forces (slow-increasing ramp and 2 Hz sine), conditioned on error magnitude (six quantiles) at each trial’s mid-point. Envelopes: SD. Scale bar: error at conditioning time. (**E**) Behavioral-fit improvement when Autoregressive Exogenous Input models (Methods) used profile-specific templates. At each moment in each trial, change-in-force was fit given recent errors and (if used) the template. Fits improved only for some profiles (orange). Bars: SD across partitions, equivalent to SEM.

In a moving multi-joint arm, muscle activity rarely mirrors endpoint force. In contrast, the Pac-Man task is isometric, involves a single degree of freedom (pushing forward), and is performed by elbow and shoulder extensors working together. Muscle activity is thus expected to correlate strongly with endpoint force, as confirmed via motor-unit recordings in Monkey C (see below). Additionally, nearly all ‘external’ parameters reflect force or its derivative. Dot-height specifies desired force, closely tracked by actual force. Pac-Man’s position and velocity mirror force and its derivative. Joint torques will be proportional to force. Stretch-receptor feedback will reflect muscle tension (which reflects force) or its derivative. Under all current hypotheses, activity should become simpler and lower-dimensional; movement is uni-directional, there are few independent parameters, and biomechanical complexity is minimized. Conversely, under the flexible-repertoire hypothesis, neural dimensionality is not bounded by external degrees of freedom but instead reflects the variety of engaged subskills.

Encouraged by a nonlinear reward schedule, monkeys became remarkably adept at matching a wide variety of target profiles (Fig. 2B, gray envelopes show across-trial SD). Observing behavior in real time (Supp. Movie 1) suggests that performing this challenging task involves multiple strategies. When force changes slowly, a reasonable strategy is to focus on reducing the current discrepancy between Pac-Man’s height and dot height (a ‘closed-loop’ strategy). In agreement, errors were transient – i.e. corrected quickly – during low-frequency profiles (Fig. 2C, left). Higher frequencies require recognizing the approaching profile, with force produced proactively using an internal template (an ‘open-loop’ strategy). A mis-scaled or mis-centered template would cause persistent errors (e.g. Fig. 2C, right). To examine error persistence, we conditioned trial-averaged forces upon error in each trial’s middle (Fig. 2D). For slowly changing profiles, error diminished steadily over ∼300 ms, consistent with closed-loop tracking. For high-frequency profiles, error disappeared swiftly then reappeared later, consistent with an internal template that almost (but not quite) matched the target. When fitting control-theory models to single-trial behavior, closed-loop strategies sufficed at lower frequencies. Higher frequencies benefited from including open-loop templates (Fig. 2E).

Open-loop strategies must be template-specific, and high-level performance may combine closed- and open-loop strategies. Under the flexible-repertoire hypothesis, this would prompt M1 to switch amongst subskills. For this task, we define a ‘subskill’ to be a computation (implemented by network dynamics) that can generate a particular type of force-profile (e.g. 2 Hz sinusoids of various phases) and/or implement closed-loop corrections. More generally, a subskill is a motor ability suitable to generate muscle-activity patterns ^40^ (including corrections) in some situations but not others ^1^. We accept a likely continuum between completely distinct subskills versus subskill variations; our analyses are designed to respect both possibilities. We also anticipate that subskills might be reused compositionally: e.g. the slow (0.25 Hz) sinusoid might recruit, sequentially, subskills used for slow rising and falling ramps. Rising and falling ramps might in turn use the same or different subskills (control policies might or might not be symmetric). How performance is (or isn’t) segmented into subskills is difficult to infer externally but should, under the flexible-repertoire hypothesis, become apparent in the neural data.

### High-dimensional neural activity shows subskill-specific features

Neural activity was recorded primarily using 384-channel NeuroPixels Probes (1.0 NHP ^39^) and pooled across multiple probes and sessions. Testing flexible-repertoire predictions requires a large population of individually well-resolved responses. We thus analyzed only stable isolations with excellent estimates of rate modulation (1257 and 864 neurons for Monkey C and I; additional recordings made for task variants described later). For example, the neuron in Fig. 3C had a firing-rate range 47 times greater than its average SEM (average ratio of 32 for this dataset; envelopes show SEMs). Microstimulation of the recorded region activated the task-relevant muscles: *anterior deltoid* and *triceps brachii*. Motor-unit recordings (134 total) employed these same muscles. Individual motor-unit spikes (corresponding one-to-one with motoneuron spikes) were sorted using customized methods ^28^. Motor-unit activity was relatively simple; each motor unit, if active, correlated with force. The example in Fig. 3A was typical: its response reflects force overall (*r* = 0.87) and within-condition (range 0.58–0.97). Motor-unit responses were not identical ^28^, but all correlated with force when active (mean *r* = 0.79). Consequently, the first principal component (PC) of motor-unit activity correlated strongly with endpoint force (*r* = 0.85).

**Figure 3.**
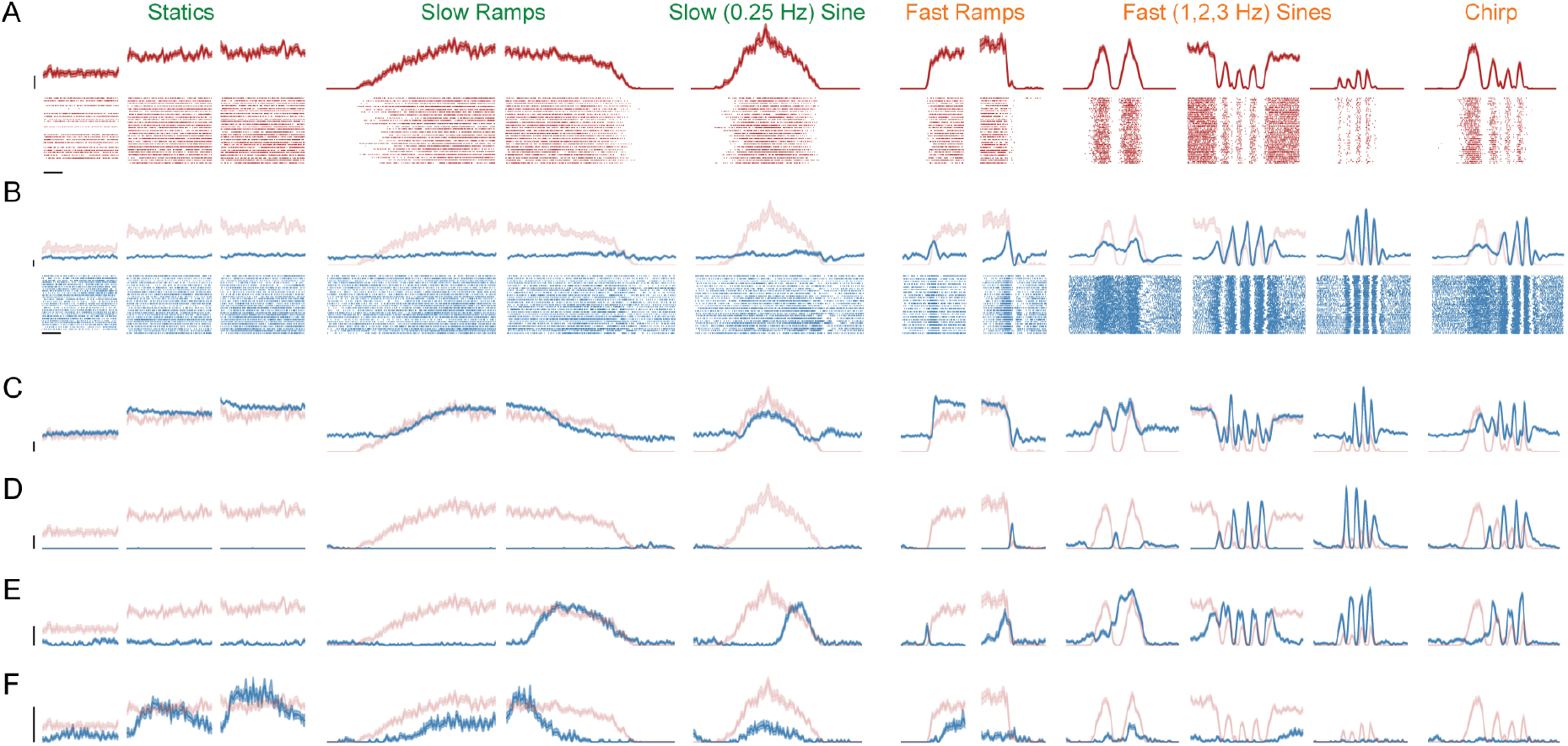
Example motor-unit and M1-neuron responses. (**A**) Trial-averaged firing-rate for an example motor unit. Envelopes indicate SEM. Horizontal and vertical bars denote 500 ms and 5 spikes/sec. Spike rasters are plotted with one line per trial. This mean motor-unit response is repeated below, providing a reference for expectations if a response reflects force. (**B**) Same but for an example M1 neuron. (**C**) Mean rate of an atypical M1 neuron whose response resembles force. (**C-F**) Additional, more typical, example M1 neurons.

The flexible-repertoire hypothesis predicts that, despite external task simplicity, M1 activity should involve many factors and thus be high-dimensional. In agreement, accounting for 90% of population variance required >75 PCs (both monkeys), versus 4 for the motor units. M1 dimensionality in reaching tasks is often estimated as ∼10–12 ^2,4^. Studies ^41–43^have stressed that dimensionality exceeds that predicted by representational models, but dimensionality in the present task is higher still, despite it using only one direction.

High-fidelity firing-rate estimates make it simple to appreciate, through inspection, the reasons for high-dimensionality. A few neurons (e.g. Fig. 3C) had responses that consistently resembled force, yet this was very rare (Supp. Fig. 2). For example, the neuron in Fig. 3B mirrors force only when force changes rapidly. The neuron in Fig. 3D responds out of phase with force, at only some frequencies. The neuron in Fig. 3E tracks force during only some portions of some conditions. The neuron in Fig. 3F responds only during low-frequency forces. These examples illustrate a common source of diversity: putatively subskill-specific features. A neuron could be active for some conditions (or some parts of some conditions) and inactive for others, despite similar force ranges. More generally, neurons typically had activity whose relationship with force changed across conditions. The combination of between- and within-condition response variety yielded extremely diverse responses that defy easy explanation under any traditional hypothesis. For example, single-neuron activity has often been observed to reflect muscle force ^6,44–46^and/or related parameters. Given traditional expectations, that relationship should be particularly straightforward in this task; forces are unidirectional and there are few other independent parameters. Instead, correlations with force (and its derivative) were generally very small (Supp. Fig. 2). Nevertheless, force could be accurately decoded (see below). Understanding this seeming paradox requires stepping through flexible-repertoire predictions.

### Neural state-space locations form a map of subskills

Noise-robust network dynamics require low trajectory-tangling: nearby states should avoid moving in different directions ^22,23^. Under the flexible-repertoire hypothesis, subskills are proposed to leverage different dynamical flow-fields. Low tangling thus requires separation of subskills into different state-space regions (Fig. 1A). When considering where activity is centered for each condition (its ‘centroid’), four predictions follow (Fig. 4A). First, to prevent overlapping / tangled trajectories, centroids should be separated in dimensions beyond those capturing within-condition trajectories. Second, centroids should be organized, with less-similar subskills having more-distant centroids (the more flow-fields differ, the greater the separation needed). Third, more-distant centroids should correspond to trajectories in less-aligned dimensions (separating dynamics into different dimensions also reduces tangling ^23^). Finally, trajectory tangling should remain low when computed across all conditions. A constrained-manifold makes different predictions (Fig. 4B): conditions may have differing means – e.g. due to differing mean forces – but these differences don’t reflect subskill *per se* (subskills reuse the same manifold), and simply occur in the same space as the trajectories.

**Figure 4.**
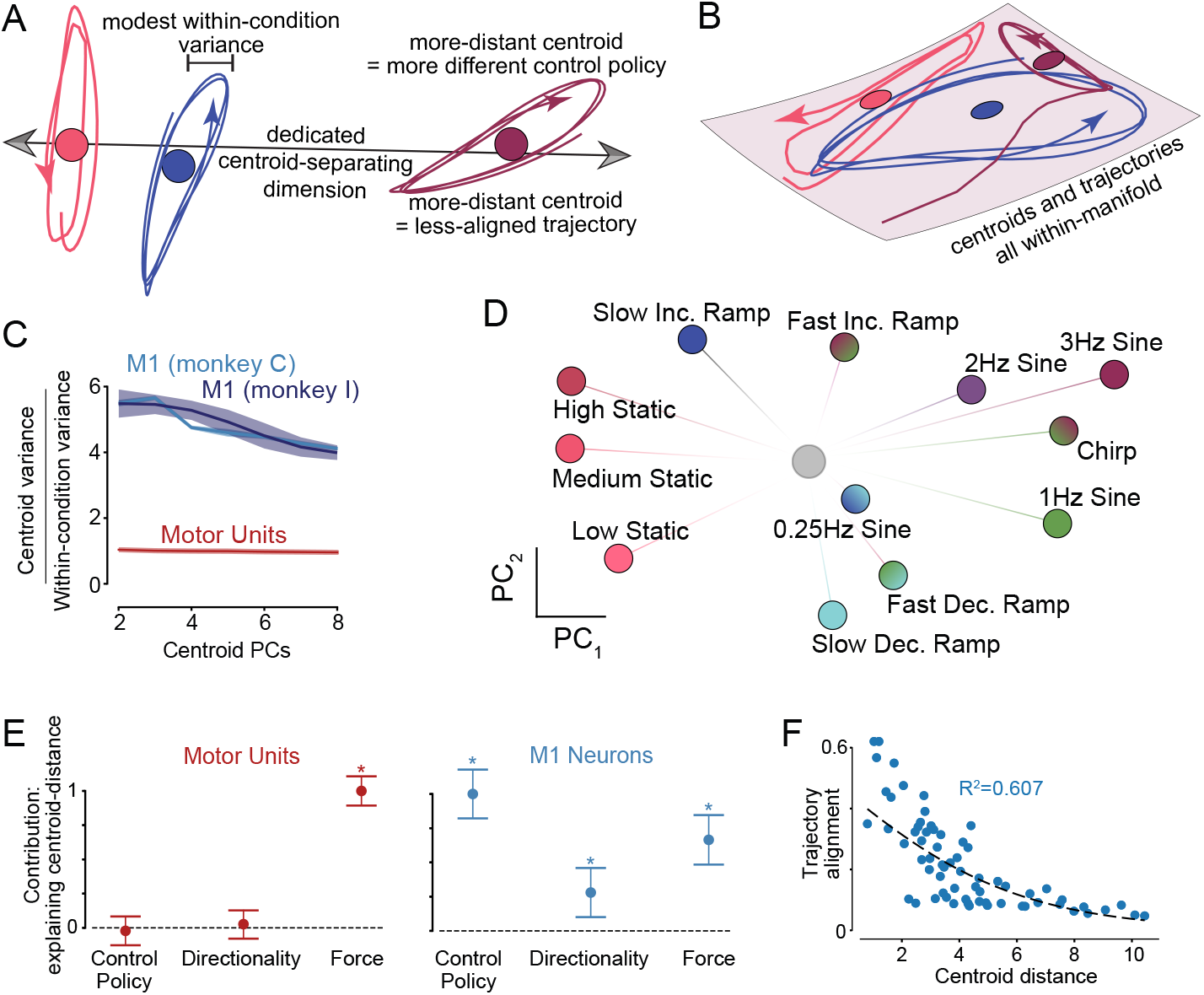
State-space location reflects subskill. (**A**) Flexible-repertoire predictions regarding each condition’s activity centroid (mean activity, colored circles) and its within-condition trajectory (colored traces). The hypothesis predicts dimensions (arrow) that preferentially separate centroids versus capturing trajectories. More-distant centroids should correspond to less-similar subskills and less-aligned trajectories. (**B**) Constrained-manifold predictions. Across conditions, trajectories may have different means, but these simply occur in the same manifold. (**C**) Ratio of centroid-variance to within-trajectory-variance (both un-normalized) captured by centroid PCs. Ratios *>*1 indicate centroid-separating dimensions differ from those that best-capture trajectories (as in A). Ratios near unity accord with panel B. Envelopes: SD of bootstrapped values (equivalent to SEM). (**D**) Organization of M1 centroids in their top PCs. Gray dot indicates baseline activity. Color choices reflect the factors engaged for that condition (assessed below). Lines connecting baseline and centroids added for visualization. Data for Monkey C. (Monkey I; Supp. Fig. 3). (**E**) Contribution of behavioral distances when fitting centroid distances. Significance (*, *p <* 0.005, two-sided) and 95% CIs computed via each coefficient’s t-statistic. Contributions are normalized (maximum one). (**F**) Trajectory alignment versus centroid-distance. One dot per condition-pair. Fit via logistic model (*p <* 10^−5^). Monkey I analysis and additional details in Supp. Fig. 3.

We computed each condition’s state-space trajectory, then estimated its centroid by averaging over time. For now we ignore the possibility of compositionality (considered later) and treat each condition as involving one potential subskill. M1 populations obeyed flexible-repertoire predictions: activity was separated in dimensions that preferentially captured centroid distances versus within-condition trajectories (Fig. 4C, blue). In contrast, the space capturing motor-unit centroids was merely the same space that captured the trajectories themselves (Fig. 4C, red). M1-centroid organization (Fig. 4D) reflected features beyond motor output. The three static centroids lay near each other, despite mean forces being very different. Increasing- and decreasing-ramp centroids lay far apart (top and bottom) despite nearly identical mean force. Conditions involving closed-loop versus open-loop strategies had particularly distant centroids.

To explore this organization, we modeled centroid distances (between all pairs, in full-dimensional space) as a linear function of three behavioral quantities: difference in mean force (force distance), the degree to which force increased versus decreased (directionality distance), and how well a control-policy model generalized from one condition to another (control-policy distance, see Methods). Force-distance should matter under nearly any hypothesis, and thus be predictive for both M1 and motor-unit populations. Directionality and control-policy distances relate to the subskill used. These distances should be irrelevant for the motor-units but should matter (under the flexible-repertoire hypothesis) for M1. Centroid-distances were fit well for both motor-units (*R*^2^ = 0.86) and M1 (*R*^2^ = 0.808, *R*^2^ = 0.691; Monkey C and I; Supp. Fig. 4). Motor-unit fits relied solely on force distance (Fig. 4E, red). All three distances were explanatory for M1, and control-policy distance was most explanatory (Fig. 4E, blue). Also as predicted, more-distant M1 centroids involved trajectories in less-aligned dimensions (Fig. 4F; alignment index described below). Thus, a substantial portion of M1 activity reflects subskill-organization rather than motor output. Also as predicted, trajectory tangling was far lower (16%) for M1 populations versus motor-units.

### Neural state-space dimensions are subskill-specific

We used the subspace-alignment index ^29^ to assess whether activity shares dimensions across condition-groups. Groups were based on force-profile features: e.g. all statics, or all fast sines. All groups included a similar range of forces and (except for statics) both increasing and decreasing forces. The alignment index ranges from zero (subspaces share no dimensions) to one (all shared). For a constrained manifold, alignment should approach unity. The flexible-repertoire hypothesis predicts varied alignment: high when two condition-groups share subskills and low otherwise. For the motor-units, alignment was consistently high (mean 0.85; *>*0.75 every comparison; Supp. Fig. 7). M1 populations had variable alignment (0.14–0.81 monkey C; 0.33–0.81 monkey I) even though all condition-groups involved force in the same direction. Low alignment occurred for condition-groups that likely used different control strategies: e.g. statics versus fast-sines (0.14 and 0.35, monkey C and I). Overall, condition-groups tended to share only half their dimensions: alignment averaged 0.41 and 0.51 (monkey C and I; also see Supp. Fig. 5, Supp. Fig. 6, Supp. Fig. 7).

**Figure 5.**
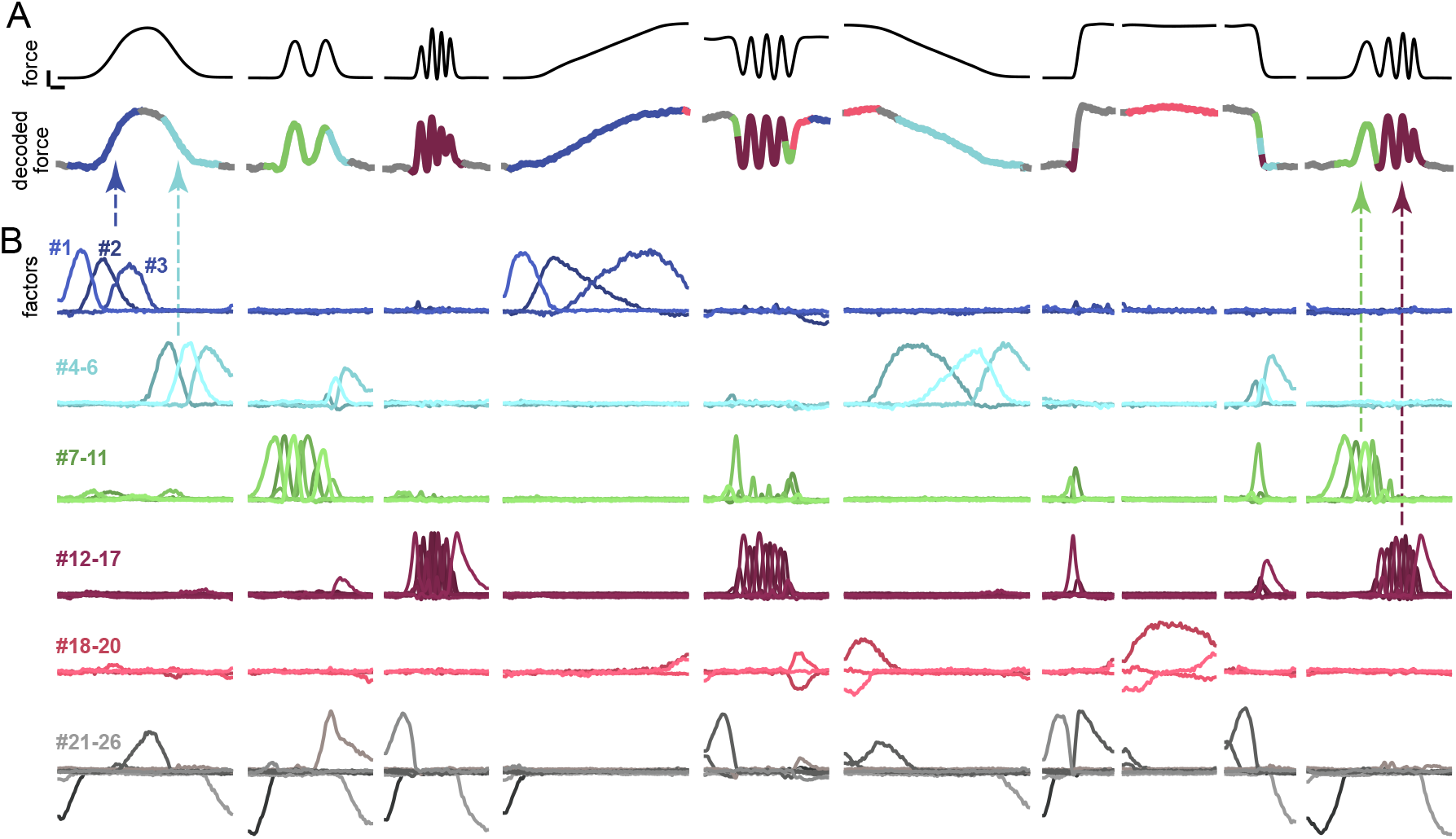
Factors are numerous and make temporally-specific contributions. (**A**) Actual and decoded force (a weighted sum of factors below) across conditions. Trace color reflects which factor group contributed most to decoding at that moment. (**B**) Each factor is a projection of the population response onto an SCA-identified dimension. Factors form rough groups, indicated by color. Arrows illustrate two conditions where decoded force depended upon sequentially active factor groups. Data from Monkey C (Monkey I; Supp. Fig. 11). Not shown are a few factors dominated by sampling error (when many SCA factors are requested, any ‘extra’ factors typically just capture noise).

**Figure 6.**
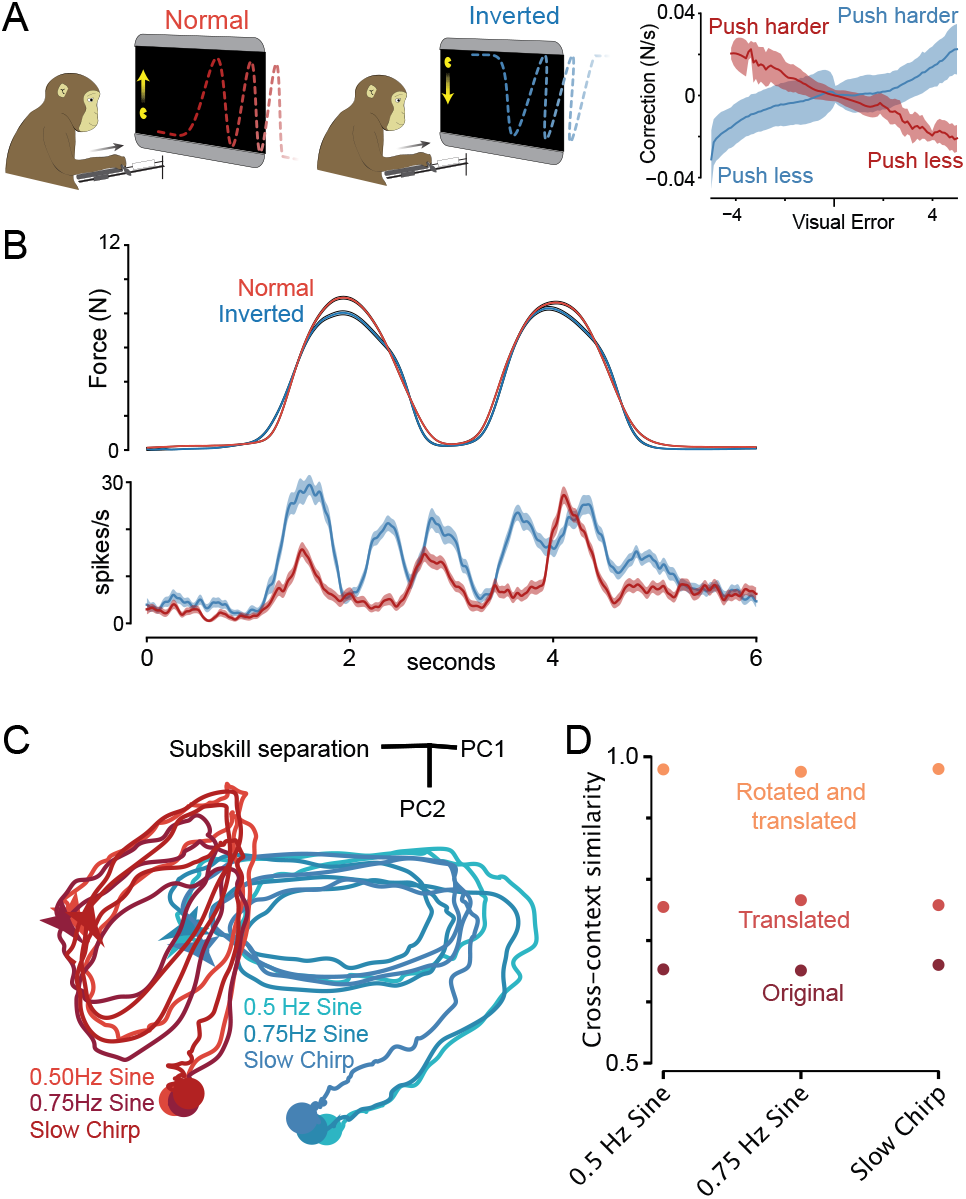
Neural Activity during Normal and Inverted Subskills. (**A**) Left: Task schematic. Right: Change in force’s derivative (*±* SD) 150 ms after visual error (Pac-Man’s height minus target height). Data for Monkey C (Monkey I: Supp. Fig. 20). (**B**) Trial-averaged force (top) and example-neuron activity (bottom) during a 0.5 Hz sine, for normal and inverted subskills. Envelopes show SEM. To be representative, the example neuron was chosen so that its between-subskill activity difference was near the population median. (**C**) State-space projections of neural activity. Two axes were found via PCA. The third captured between-subskill differences. (**D**) Cross-context similarity of the population response (Methods) between normal and inverted subskills. Before computing similarity, data underwent no transformation (maroon), a similarity-maximizing translation (red-orange), or a similarity-maximizing rotation and translation (orange). Analysis used a relatively high (30) dimensional space to ensure trajectory differences were not lost.

Why do these results differ from Gallego et al. ^4^, who found strong alignment (*>*0.8) across sub-tasks, or Golub et al. ^3^, who found preserved covariance? The flexible-repertoire hypothesis predicts high alignment when situations reuse subskills, which seems likely in these prior tasks. In agreement, we also observed examples of reuse: e.g. the slow sine and slow-ramp group were well-aligned (0.81 for both monkeys), unlike fast-sine and slow-ramp groups (0.36 and 0.44). Under the hypothesis of reuse, activity during the rising phase of the slow sine should overlap dimensions for the slow-rising ramp, and come closest to that centroid. During the falling phase, activity should overlap dimensions for the slow-falling ramp, and come closest to that centroid. This was confirmed, along with additional forms of reuse (Supp. Fig. 9, Supp. Fig. 10).

### Population factors reflect selective recruitment of subskill-specific dynamics

Under the flexible-subspace hypothesis, M1 expresses a rich variety of subskill-specific dynamics, each employing a group of factors. There should thus be many total factors spanning many total dimensions. Each factor-group is predicted to be sparsely active, becoming engaged only when that subskill is used (Fig. 1E) and generating (at those moments but not others) a basis for outgoing commands. Large-scale recordings should allow direct tests of these predictions, via examination of estimated factors, if one can bypass limitations of standard factor-estimation methods. For instance, PCA tends to combine sparsely active factors, something traditionally overcome through supervision: guessing which conditions or times might use different factors ^29^. To avoid such guessing and allow potential subskill-divisions to emerge directly from the data, we employed a novel unsupervised method, Sparse Component Analysis (SCA) ^47^. SCA behaves much like PCA (and captures similar overall variance: 94% as much for both monkeys), but avoids PCA’s tendency to concentrate variance in its first components and thus mix sparsely active factors. If factors are not sparse, SCA cannot make them so, nor does it impose assumptions regarding the nature of potential sparsity. Under the constrained-manifold view, SCA should thus identify relatively few factors, most active at most times. Under the flexible-repertoire hypothesis, SCA should identify many (and thus individually low-variance) factors, with different factor-groups active at different moments. Yet sparsity should not be too extreme; factor-groups should be reused when different moments can leverage the same subskill.

SCA identified many factors (Fig. 5B), which formed rough groups. A ‘blue’ factor-group was engaged only when force rose slowly: during the 0.25-Hz sine and slow rising ramp. A ‘cyan’ factor-group became engaged when force decreased slowly. ‘Green’ and ‘purple’ groups were engaged during medium-frequency (1–2 Hz) and high-frequency (2–3 Hz) sinusoids. A ‘red’ group was engaged when high force was maintained. A ‘gray’ group tended to reflect transitions: e.g. the darkest gray factor increased before each transition from rest. Force was accurately decoded from these factors (Fig. 5A, *R*^2^ = 0.98), much as it was from the full population (cross-validated analysis in Supp. Fig. 16). Different factor-groups contributed at different moments (reflected by decoded-trace color, Fig. 5A). No factor was dominant; each contributed 1.1–6.3% of total captured variance. A factor that accounts for 2% of population structure would traditionally have been considered irrelevant (and would have been difficult to resolve in any case). Yet these low-variance factors made large contributions at key moments. For example, artificially silencing the blue factor-group impaired decoding of slowly increasing forces. Dynamics differed across conditions (Supp. Fig. 12), reflecting the active factors; e.g. 1 Hz and 3 Hz sines involved medium- and high-frequency oscillatory population dynamics, reflecting (respectively) green and purple factor-group activity.

SCA cannot ‘invent’ unrepresentative sparse factors ^47^. Doing so would significantly increase reconstruction error, something discouraged by the objective function (Methods) and absent empirically (Supp. Fig. 15). Two further observations illustrate this point. First, motor-unit SCA factors were not sparse (Supp. Fig. 14), even when sparsity was encouraged at the expense of increased reconstruction error (Supp. Fig. 15). Second, M1 SCA factors were in fact very representative of single-neuron response features. M1 responses were extremely diverse, consistent with numerous factors. Some neurons were sparsely active, just like the factors. More generally, neurons displayed condition-dependent relationships with force, as expected if a neuron reflects multiple sparsely active factors (which was indeed common; Supp. Fig. 13). Factor activity also accords with subspace-alignment results (discussed below). Thus, SCA simply reveals, in greater detail and without supervision, effects visible in other ways.

The presence of many sparsely active factors upends some common expectations. For example, in a ‘force-centric’ task, one would traditionally expect force to be strongly represented. And in one sense it was: a force-readout dimension yielded accurate decoding (cross-validated *R*^2^ = 0.99 and 0.96; Monkey C and I). However, across different degrees of regularization, this dimension captured at most 13% and 7% of population variance (Supp. Fig. 16). This seeming paradox is resolved by the observation that (as predicted in Fig. 1E) force decoding relied on many transiently active factors. In contrast, the motor-unit force-readout dimension captured up to 73% of population variance (Supp. Fig. 16) and relied on one continuously active factor (Supp. Fig. 14). Thus, force was encoded in a traditional manner by the motor units, and in a very different way by the M1 population.

### Subksills are reused compositionally across conditions

M1 activity is unexpectedly sparse, yet less sparse than it could be; neither M1 factors nor M1 neurons approached the extreme sparsity found in songbird area HVC ^48^. Factors were often active for sizable spans of time, and were reused in new combinations across conditions. Arrows (Fig. 5) highlight two examples of sequential reuse. The 0.25 Hz sine engages blue then cyan factor-groups, also engaged (respectively) during slow increasing and decreasing ramps. The chirp engages green then purple factor-groups, also engaged (respectively) during 1–2 Hz and 2–3 Hz sines. Fast ramps recruit green and purple groups simultaneously (plus the cyan group for the decreasing ramp), perhaps reflecting the ramp’s broad frequency content. The oscillatory dynamics of green and purple factor-groups, patent during 1-3 Hz sines, are less-obvious during fast ramps yet still visible in the eigenvalue spectra of dynamical fits (Supp. Fig. 12H), reminiscent of what occurs during swift reaches ^19,49^. Some factor combinations (e.g. coactive cyan, green and purple groups) were specific to one condition (e.g. fast-decreasing ramps).

The scale of our recordings allowed us to estimate the timecourse of ∼26 (monkey C) and ∼24 (monkey I) factors. (There may, of course, have been more active factors than we could accurately resolve). Each condition engaged only a subset of these factors: 7.3 on average (range 1-14) for monkey C and 6.3 on average (range 3-9) for monkey I. Accordingly, a given factor was engaged during only some conditions: on average 3.0 of 12 (monkey C; range 1-6) and 4.9 of 11 (monkey I; range 2-7). These findings accord with subspace-alignment results above: conditions typically shared only some dimensions (each factor employs one dimension). Conditions likely to employ multiple subskills engaged more factors: e.g. the chirp engaged 14 and 9 (monkey C and I). Simpler conditions engaged fewer: the high static engaged 2 and 4 respectively. Factor activity replicates (and explains) reuse of dimensions between some conditions but not others (Supp. Fig. 9, Supp. Fig. 10). For example, during the chirp, sequential use of green and purple factors explains the progressive alignment with dimensions used during 1, 2 and 3 Hz sines. Conversely, alignment was low when conditions did not share engaged factors.

### The same motor output can be subserved by different subskills

The results above indicate that M1 activity strongly reflects subskill, even more so than motor output. Why was this large effect not previously apparent? One likely reason is that, even when an experiment uses two sub-tasks (e.g. variants of center-out reaching ^4^), they may reuse subskills, yielding a conserved manifold. At the other extreme, for the few prior tasks that surely engaged different subskills (e.g. walking versus lever-pulling ^30^), the axes of motor output differed dramatically. Neural activity is thus guaranteed to differ under any hypothesis, including engaging different output-dimensions within a low-dimensional manifold. The Pac-Man task sidesteps this interpretational hurdle by keeping force direction constant. An even more stringent test would compare neural activity across subskills with matched time-varying outputs. We thus trained monkeys to perform an ‘inverted’ Pac-Man task, in addition to the normal version (Fig. 6A). The inverted task used blue dot paths, with Pac-Man moving downward (from zero force at the top) with increasing forward force. By also inverting visual target profiles, we created condition-pairs with matched forces (Fig. 6B; Supp. Fig. 19, Supp. Fig. 20).

Visuomotor reversals are challenging during continuous-tracking tasks ^1,50^; they cannot be solved by rapid adaptation nor by purely cognitive strategies. Consequently, extensive practice was required to accurately perform both versions in interleaved trials. Post-training, rapid responses to small errors had opposing signs (Fig. 6A), confirming subskill-specific control policies. An intriguing possibility is that two related but distinct internal subskills may develop within M1. Each subskill’s trajectory would be a copy of the other, and they would produce the same typical motor output, yet they would unfold in different locations and dimensions. In contrast, if changing motor output ^30,35,37^is necessary to cause changes in M1 neural dimensions / locations, then neural activity should be nearly identical across these subskills.

Single-neuron responses differed across subskills, despite matched forces (Fig. 6B, further examples and analyses in Supp. Fig. 19, Supp. Fig. 20, Supp. Fig. 23). Neural trajectories shared similar shapes across subskills (Fig. 6C) and could be brought into almost perfect register via translation and rotation (Fig. 6D; Supp. Fig. 21). Yet trajectories were centered in different locations and occupied different (but not orthogonal) dimensions. Between-subskill trajectory differences were sizable relative to across-force-profile differences (the three similar-colored trajectories). A direct impact of visual features (e.g. dot color) on M1 activity cannot be ruled out, but is unlikely; e.g. we previously found preserved neural activity when different visual cues prompted the same reaches ^51^.

### Different motor outputs can be subserved by the same subskill

Force was closely, but not perfectly, matched across normal and inverted subskills. We performed an additional experiment and set of recordings, in monkey C, to control for these small differences. This experiment also further tests the flexible-repertoire prediction that activity should be influenced more by subskill differences than by motor-output differences. To create a scenario where output differences exceed subskill differences, the gain linking force and Pac-Man’s height was incrementally decreased within a session, eventually requiring 67% more force to produce the same height (Supp. Fig. 22). Adaptation to each gain decrement was essentially immediate, consistent with swift alteration of an existing subskill rather than gradual learning of a new one ^1,52^.

Changes in neural activity were much smaller than between normal and inverted Pac-Man, despite far larger changes in force (Supp. Fig. 23, Supp. Fig. 24). The flexible-repertoire hypothesis predicts that, for both large (across-subskill) and modest (during adaptation) changes in activity, motor output should be consistently decodable via a stable low-variance dimension. This was confirmed: *R*^2^ = 0.96 and 0.97 (monkey C and I) when decoding across normal and inverted subskills, and *R*^2^ = 0.96 across gain changes. Readout dimensions captured modest population variance: at most 8.7, 5.0, and 11.7% respectively.

## Discussion

Our findings confirmed the predictions made by the flexible-repertoire hypothesis. They thus argue that M1 instantiates a repertoire of learned computations – one that may be vast when considering our full behavioral repertoires. This perspective evokes a very different picture regarding the nature and purpose of movement-related cortical activity. Its dominant features don’t reflect motor commands, movement parameters, or target properties as has often been assumed, but instead reflect the currently engaged subskill. Rather than being constrained to a low-dimensional manifold, activity moves flexibly amongst many locations and dimensions. Different experimental manipulations cannot be presumed (as in many classic experiments ^5,6^) to provide different views of a consistent M1 computation or representation; they may instead recruit very different computations. This flexibility is surprising for a primary cortical area that communicates directly with the spinal cord. By analogy with primary visual cortex, one might expect M1 to perform a relatively fixed computation. Instead, our data argue that M1 instantiates many acquired computations, organized within a very high-dimensional activity space. These findings speak to a burgeoning interest in the computational basis of motor and cognitive flexibility ^10–16,53–57^. It appears increasingly likely that understanding intelligent behavior requires understanding how neural networks instantiate large repertoires of computations and navigate amongst them to meet the needs of the moment. Our results reveal that this occurs in motor cortex, and document its empirical signatures. These findings illustrate the kind of computational flexibility that our brains possess, and that our field’s theories and hypotheses should attempt to explain. This is not to deny that many computations – e.g. keeping track of heading direction, or maintaining eye position – employ fixed neural manifolds to robustly perform a consistent computation. Yet if a ‘low-level’ primary area such as M1 is flexible, then flexibility may be more the rule than the exception.

The flexible-repertoire hypothesis integrates ideas regarding network dynamics ^9,49,58–60^, subspaces ^18,61–63^, and optimal feedback control ^64–67^. To instantiate different subskills, including subskill-specific control policies ^14,68^, a network must employ a variety of dynamics. This can be accomplished by using different locations and subspaces ^23,36,37^, arranged in a neural space where most dimensions are output-null ^9^. The proposal that distinct M1 subspaces can support distinct computations was initially motivated by orthogonality of preparatory and execution subspaces ^29^. There have since been multiple examples of situation-specific neural dimensions ^23,30–37^(sometimes accompanied by changes in state-space location ^23,35,37^). Interpretations have varied regarding whether this is more likely to reflect different computations versus motor outputs. Motor output is surely sometimes a key factor ^30,35^, but computation was also deemed potentially relevant: low trajectory tangling requires isolating different dynamics via location and dimensions ^9,22,23^. Our results confirm that different subskills are, on their own, sufficient to cause neural activity to change locations and dimensions. This is true even when motor-output dimensions do not change – indeed, even when motor-output is continuously matched. Furthermore, different neural locations can provide not only variants of the same dynamics (as in ^23,37^), but qualitatively different dynamics and/or control policies.

Synthesizing prior ideas is critical to explain these results – each on its own is insufficient. As one example, the feedback-control model of Kalidindi et al. ^69^ uses limited neural dynamics and incorrectly predicts low-dimensional activity dominated by muscle commands and proprioception. However, their control-centric focus becomes compatible with present results – indeed it helps explain them – if one assumes that rich dynamics instantiate many subskill-specific control policies. As another example, although our results are incompatible with a low-dimensional manifold, that idea remains useful within-subskill. Each subskill’s flow-field presumably constrains how local trajectories unfold ^20,27^. Suppose only one existing subskill is appropriate to the current task. Neural activity would indeed be constrained until learning alters that subskill ^26^. Similarly, low-dimensional network solutions (created via construction ^70^ or training ^59^) often provide good matches with empirical data ^63^. Indeed, encouraging lower-dimensionality can be essential for realistic solutions ^49^. Such networks presumably remain good models of within-subskill dynamics. That said, it is not simply that the flexible-repertoire hypothesis proposes a much higher-dimensional (and less-linear) manifold. By design, the combinatorial expressivity of compositionality – especially the ability to superimpose or blend subskills – will always be capable of generating activity outside a previously observed manifold.

It was recently shown that force-field adaptation shifts activity (in M1 and premotor cortex) during preparation but not movement ^71^. Additionally, new stimulus-movement associations alter preparatory (but not movement) dimensions in mouse premotor cortex ^38^. These findings presumably relate to present results, but in the domain of preparation rather than movement. Thus, the principle of instantiating different learned computations in different locations / dimensions is likely very broad ^9^.

Behavioral investigations indicate normative principles governing how subskills are organized and updated ^1,72–74^. A common metaphor is a landscape containing skills, subskills, and their variants. This landscape is navigated by cognitive-motor processes to produce the variety of actions needed to move within a complex world ^75^. Combined with findings regarding movement variants ^23,37^, our results suggest this conceptual landscape may be reflected in the literal state-space organization of neural activity. This raises intriguing possibilities. Might a subskill’s neural scaffolding be replicated, in a nearby location, when learning a new version of that subskill (a possibility raised by normal-and-inverted Pac-Man results)? Might learning to play squash initially rely on tennis subskills ^1^, with squash-specific subskills ‘split off’ over time? The flexible-repertoire hypothesis makes it possible to pose such questions, and to inquire how our motor repertoires develop and are navigated intelligently by upstream areas.

## Acknowledgments

We thank Yanina Pavlova for expert monkey care. We thank Andrew J. Zimnik, Saurabh Vyas, and Sean M. Perkins for their helpful feedback on figures and analyses. We thank Ciela Chavez-Gilbride, Aleyna Silcott, Mahham Fayyaz, Rozanna Yakub, and Allison Ong for administrative support. Most importantly, we thank subjects C and I for their time and patience. Research was supported by the Grossman Charitable Trust, the Simons Foundation, and NINDS R01NS135240. E.A.A was supported by a National Science Foundation graduate research fellowship, the Kavli Foundation, and Gatsby Foundation award GAT3708. E.M.T was supported by Brain and Behavior Research Foundation NARSAD Young Investigator Grant and a Grossman Center fellowship. N.J.M was supported by NIH F31 NS110201 and a National Science Foundation GRFP fellowship. D.M.W was supported by NINDS R01NS117699. M.N.S was supported by the Howard Hughes Medical Institute and a Grossman-Kavli Scholar award.

## Author Contributions

N.J.M. and M.M.C. designed the basic Pac-Man task, and N.J.M., E.A.A. and E.M.T. trained monkeys to perform that task. E.M.T., M.N.S., D.M.W. and M.M.C. designed the additional task variants (normal-and-inverted Pac-Man and multi-gain Pac-Man), and E.M.T. and E.A.A. trained monkeys to perform those additional tasks. E.A.A., E.M.T. and N.J.M. performed the neural recordings for the basic Pac-Man task. E.M.T. and E.A.A. performed the neural recordings for the additional task variants. N.J.M. and M.M.C. performed motor-unit recordings. E.A.A. and E.M.T. performed analyses, with guidance from M.M.C., L.F.A., M.N.S. and D.M.W. E.A.A. and M.M.C. created the figures. E.A.A. and M.M.C. wrote the initial manuscript. All authors contributed, often extensively, to editing, revisions, and modifications to analyses and figures.

## Methods

### Subjects and Task

All protocols were in accordance with the National Institutes of Health (NIH) guidelines and approved by the Columbia University Institutional Animal Care and Use Committee (protocol number AC-AABT8672). Subjects were two adult male macaques (Monkeys C and I, Macaca Mulatta), 10 and 11 years-old. During each experiment, the subject sat in a custom-built primate chair with the head restrained via surgical implant and the right arm comfortably secured. To perform the task, subjects grasped a handle with their left hand while resting their left forearm on a small platform that supported the handle. During initial training, subjects developed a stereotyped grasp posture that they found comfortable. To ensure consistent placement within and between sessions, once they achieved this posture we applied tape around the hand and Velcro around the forearm. The handle did not move; isometric force was measured by an attached load cell.

By pressing on the handle, subjects controlled a ‘Pac-Man’ icon, displayed on an LCD monitor (Asus PN PG258Q, 240-Hz refresh, 1,920 × 1,080 pixels) using Psychophysics Toolbox 3.0. The horizontal position of the Pac-Man icon was fixed on the left side of the screen, and its vertical position was directly proportional to the forward force registered by the load cell. A 0 N applied force corresponded to a resting position at the screen’s bottom. Larger forces caused Pac-Man’s position to rise. For the task-variant where we altered the gain (Supp. Fig. 22), the relationship between vertical movement and force was reduced (from the normal gain of 1) to 0.8 and then to 0.6. For the ‘inverted’ task (Fig. 6), the resting position was at the screen’s top and forward forces moved the Pac-Man icon downwards. Earlier experiments for monkey C used a single-degree-of-freedom load cell that measured only forward force. All experiments for monkey I and later experiments for monkey C employed a six-degree-of-freedom load cell. This was helpful in confirming, during the normal and inverted Pac-Man task, that forces were similar not only in the forward direction but also in off-axis directions.

Different force profiles were presented in interleaved trials. In each trial, a series of dots scrolled leftward on the screen at a constant speed (1,344 pixels per second). Subjects modulated Pac-Man’s position to intercept the dots, for which they received a juice or water reward (based on preference). Thus, the shape of the scrolling dot path determined the temporal force profile that the monkey needed to apply to the handle to obtain reward. We trained subjects to generate static, step, ramp, and sinusoidal forces over a range of amplitudes and frequencies. We define a ‘condition’ as a particular target force profile (for example, a 2 Hz sinusoid) that was presented on many ‘trials’, each a repetition of the same profile. Each condition included a ‘lead-in’ and ‘lead-out’ period: a static profile of 1 second (0.75 seconds for Monkey I) attached to the beginning and end of the target profile. Trials lasted 2.25–6 seconds, depending on the particular force profile. Juice was given throughout the trial, so long as Pac-Man successfully intercepted the dots, with a large ‘bonus’ reward given at the end of the trial. The reward schedule was designed to be encouraging: greater accuracy resulted in more frequent rewards (every few dots) and a larger bonus at the end of the trial. At high-frequencies, even slight errors in response phase (or latency) translate into a large vertical error, between Pac-Man’s height and dot height. To avoid discouraging failures, we tolerated small errors in the response phase at high frequencies. For example, if the target profile was a 3-Hz sinusoid, it was considered acceptable if the monkey generated a sinusoid of the correct amplitude and frequency but that led the target by 100 ms. To enact this tolerance, the target dots sped up or slowed down to match his phase. The magnitude of this phase correction was scaled with the target frequency and was capped at *±*3 pixels per frame. To discourage inappropriate strategies (for example, moving randomly or holding in the middle with the goal of intercepting some dots), a trial was aborted if too many dots were missed (the criterion number was tailored for each condition). These criteria led to excellent performance on the vast majority of trials. Aborted trials were not analyzed, nor were trials where the generated force had an unusual profile that could not be aligned with other trials (and thus could not be included in trial averaging).

### Surgical Procedures and Intracortical Recordings

After initial training, we performed a sterile surgery to implant head restraints and recording chambers, positioned to allow access to the arm area of primary motor cortex (M1), including both sulcal and surface cortex. We also recorded from the immediately adjacent dorsal premotor cortex (PMd), where response properties are very similar to those in surface M1. All recordings were made in the right hemisphere. Chamber positioning was guided by structural magnetic resonance imaging prior to implantation. We used intracortical microstimulation to confirm that recordings were from the arm region of motor cortex (biphasic pulses, cathodal leading, 250 ms pulse-width delivered at 333 Hz for a total duration of 50 ms). Microstimulation typically evoked contractions of the shoulder and upper arm muscles, including the *deltoid* and *triceps*. Current thresholds ranged between 20 and 200 *µ*A depending on the location and depth of stimulation (this wide range of current thresholds is expected, as we recorded in both M1 and PMd and from a variety of depths).

In Monkey C, the earliest recordings were made with 32-channel Plexon S-Probes, then the pre-production passive version of the primate-specific NeuroPixels probe, which allowed recording from 128 channels. Digitization (30 KHz) employed Blackrock Neural Signal Processors. Thereafter, and for all recordings in Monkey I, we employed the primate-optimized Neuropixels 1.0-NHP probes ^39^. Each such probe provided recordings from 384 channels, selected from 4416 total. Digitization (30 KHz) employed the standard system integrated within each Neuropixels probe, with data acquisition via connection to a PXIe system.

For Monkey C, the Neuropixels 1.0-NHP probe was held using a standard 0.25” dovetail mount rod with a custom adapter to mount it to a hydraulic drive (Narishige Inc.). A 21 gauge blunt guide tube, 25 mm in length, was held using a custom fixture and placed over the desired recording location. The dura was then penetrated with a tungsten electrode (FHC, size E), which was bent at 27 mm to prevent the tip from inserting further than 2 mm past the end of the guide tube. The tungsten electrode was inserted manually via forceps, once or several times, as necessary, which also provided feedback on the depth and difficulty of penetrating the dura. The Neuropixels 1.0-NHP probe was then aligned using the Narishige tower XY stage, lowered into the guide tube, and carefully monitored to ensure that the tip of the probe was aligned with the small dural opening created by the tungsten electrode.

For Monkey I, the Neuropixels 1.0-NHP probe was held using a custom fixture mounted to a linear rail bearing (IKO Inc.). This apparatus was designed to enable close packing of many probes, and to solve the challenge of precisely targeting structures deep in the brain. The linear rail was mounted in a custom 3D printed base, mounted directly to the recording chamber. The geometry of the 3D printed base component determined the insertion trajectories and prevented mechanical interference between probes and the chamber. This base also provided support for either sharp or blunt guide tubes, as required. In general, blunt guide tubes were preferred, but if necessary sharp guide tubes were sometimes used when the dura had become thicker and difficult to penetrate. The linear bearing was connected to a commercial drive system (NAN Inc.) via a ∼50 mm long, 0.508 mm stainless steel wire, which provided rigid connection between the NAN drive electrode mount and the Neuropixels probe mounted on the rail bearing, yet also allowed a small amount of misalignment between the drive axis and the insertion axis. This apparatus simplified the procedure of using many probes in a small space, and avoided relying on commercial drives to provide the mechanical rigidity required to safely insert a delicate probe. Additional details on the custom hardware is provided in the Neuropixels 1.0-NHP user wiki ^39^.

### Spike Sorting and Unit Inclusion

Recordings were made at tip depths ranging from 5.6 mm to 12.1 mm relative to the dural surface (note that recording sites spanned multiple mm above the tip). Digitized voltages were spike-sorted using KiloSort 2.5. Sessions were analyzed only if probe drift was minimal. Both well-isolated single units and high-quality multi-units were included in analysis, after restriction based on several criteria. To be included in analysis, a unit had to be stable throughout the full session, had to have an average rate (across all times and conditions) *>* 1 Hz, and had to pass a threshold in terms of our ability to accurately estimate its modulation. For each unit we computed a signal-to-noise ratio (SNR). Signal was the modulation of the mean rate, assessed as its standard deviation across time and conditions. Noise was estimated as the average standard error of that mean (standard error computed across trials, then averaged across time and conditions). The ratio of these two numbers is high if the mean rate is both well-modulated and accurately estimated. We analyzed an isolation only if it had an SNR *>* 1.25. When considering a neuron’s firing rate, its across-time-and-condition range is considerably larger than its across-time-and-condition standard deviation. This is especially true if a neuron is particularly active in some situations but not others. Consequently, the above criterion (SNR *>* 1.25) yielded analyzed units that had firing-rate ranges considerably larger than the standard error with which that rate could be estimated. For example, this ratio was on average 31.9 (monkey C, 1257 analyzed units) and 12.6 (monkey I, 864 analyzed units) for the main datasets. An additional 477 units (average range-to-SEM ratio: 20.3) were analyzed for sessions where monkey C performed the normal-and-inverted Pac-Man task. (Monkey I performed both the basic task and the inverted task within the same sessions, so all recordings are from the same neurons). A further 217 units (average range-to-SEM ratio: 15.0) were analyzed for sessions where monkey C performed the task while being required to adapt to changes in gain.

### Trial Alignment and Averaging

Single-trial spike rasters, for a given neuron, were converted into a firing rate via convolution with a 25-ms Gaussian kernel. One analysis (Supp. Fig. 2) focused on single-trial responses, but most employed trial averaging to identify a reliable firing rate. To do so, trials for a given condition were aligned temporally on the moment when the target force profile ‘began’ – i.e. when the target force profile, specified by the dots, reached Pac-Man. Alignment brought the actual (generated) force profile closely into register across trials. However, because the actual force profile could sometimes slightly lead or lag the target force profile, some modest across-trial variability remained. We thus realigned each trial, by shifting it slightly in time to minimize the mean squared error (MSE) between the actual force and the target force profile. This ensured that trials were well-aligned in terms of the actual generated forces. Trials that could not be well-aligned despite searching over shifts from [−200, 200] ms were excluded from analysis. Alignment was applied to all conditions except the static forces, for which there was no need to do so. After alignment, the average firing rate (and its standard error) was simply the across-trial average (and standard error) of the filtered spikes.

### Data Preprocessing

PCA and other latent-factor estimation methods can be biased towards capturing the responses of a few high firing-rate neurons. For instance, a neuron whose firing-rate range is 10–100 Hz has more than 20 times the variance of a neuron whose firing-rate range is 10–30 Hz. Because neurons vary in their firing-rate range, any population will inevitably contain a subset of high-rate neurons, whose activity will dominate estimates of the latent factors. When estimating factors, this increases sampling error because factor-estimates are effectively based on fewer neurons. A common approach is thus to standardize responses such that each neuron has unit variance across time and conditions, allowing all neurons to contribute equally. This approach has the disadvantage that sampling error can be exacerbated for a different reason: low-firing rate neurons tend to have lower signal-to-noise ratios, and normalization increases the contribution of such neurons.

One wishes to strike a balance that minimizes these two effects. To do so, we employed a soft-normalization methods that we have consistently used across many studies (e.g. ^19,22^). Each neuron’s firing rate is soft-normalized by dividing by the firing-rate range plus a constant (5). To illustrate, consider that, when applied to the hypothetical neurons described above, soft-normalization shifts their effective firing rate range to be more equal: 0.095–0.95 and 0.286–0.857. The first neuron still contributes more variance, but only by about a factor of two. Soft-normalization thus allows factor estimation to be based more equally on all neurons, while still ensuring that neurons with low rates (and low SNRs) make smaller contributions.

### Motor-unit recordings

A motor unit is defined as a spinal motoneuron and the muscle fibers it uniquely innervates. When the motoneuron spikes, so do all its fibers, with essentially 100% reliability. Muscle-fiber spikes can be recorded intramuscularly. Each such spike corresponds, one-to-one, with the spike of a spinal motoneuron. A given muscle contains (depending on its size) on the order of 100 motor units. An advantage of recording motor-unit spikes (rather than bulk EMG) is that it facilitates direct comparisons between two neural populations: M1 neurons and spinal motoneurons. To aid such comparisons, motor-unit spiking was converted to firing-rate estimates using the same procedures as for M1 neurons (described above). All analyses of M1 population and motor unit populations thus paralleled one another. Motor-unit recordings were greatly aided by the isometric task we used, which provided unusually good recording stability, such that the voltage waveform, corresponding to a given motor-unit’s spike, remained consistent and sortable over the session.

As described previously ^28^, intramuscular EMG activity was recorded acutely using paired hook-wire electrodes (Natus Neurology, PN 019-475400). Electrodes were inserted ∼1 cm into the muscle belly using 30 mm x 27 G needles. Needles were promptly removed and only the wires remained in the muscle during recording. Wires were thin (50 um diameter) and flexible and their presence in the muscle is typically not felt after insertion, allowing the task to be performed normally. Wires were removed at the end of the session. We employed several modifications to facilitate isolation of MU spikes. As originally manufactured, two wires protruded 2 mm and 5 mm from the end of each needle (thus ending 3 mm apart) with each wire insulated up to a 2 mm exposed end. We found that spike sorting benefited from including 4 wires per needle (i.e., combining two pairs in a single needle), with each pair having a differently modified geometry. Modifying each pair differently meant that they tended to be optimized for recording different MUs; one MU might be more prominent on one pair and a different MU more prominent on the other pair. Electrodes were thus modified as follows. The stripped ends of one pair were trimmed to 1 mm, with 1 mm of one wire and 8 mm of the second wire protruding from the needle’s end. The stripped ends of the second pair were trimmed to 0.5 mm, with 3.25 mm of one wire and 5.25 mm of the second wire protruding. Electrodes were hand-fabricated using a microscope (Zeiss), digital calipers, precision tweezers and knives. During experiments, EMG signals were recorded differentially from each pair of wires with the same length of stripped insulation; each insertion thus provided two active recording channels. Four insertions (closely spaced so that MUs were often recorded across many pairs) were employed, yielding eight total pairs.

Raw EMG voltages were amplified and analog filtered (band-pass 10 Hz - 10 kHz) with ISO-DAM 8A modules (World Precision Instruments), then digitized at 30 kHz with a neural signal processor (Blackrock Microsystems, Cerebus). EMG signals were digitally band-pass filtered online (50 Hz - 5 kHz) and saved. Offline, and prior to spike sorting, EMG signals were digitally filtered using a second-order 500 Hz high-pass Butterworth. Any low signal-to-noise channels were omitted from analyses. Motor unit (MU) spike times were extracted using a custom semi-automated algorithm. We adapted recent spike-sorting advances, including methods for resolving superimposed waveforms, documented extensively in ^28^. As with standard spike-sorting algorithms used for neural data, individual MU spikes were identified based on their match to a template: a canonical time-varying voltage across all simultaneously recorded channels. Spike templates were inferred using all the data across a given session. A distinctive feature of intramuscular records (compared to neural recordings) is very high signal-to-noise: peak-to-peak voltages are on the order of mV, rather than uV, and there is negligible thermal noise. At the same time, it is common for more than one MU to spike simultaneously, yielding a superposition of waveforms. This is relatively rare at low forces but can become common as forces increase. The motor-unit spike-sorting algorithm was thus tailored to detect not only voltages that corresponded to single MU spikes, but also those that resulted from superposition of multiple spikes. Detection of superposition was greatly aided by the multi-channel recordings; different units were prominent on different channels. We analyzed artificial spiking data with realistic properties (e.g. recruitment based on the actual force profiles we used, and waveforms based on actual recorded waveforms) to verify that spike-sorting was accurate in circumstances where the ground truth was known. Spike sorting criteria were extremely stringent, so that only very well isolated motor units were analyzed. Each recording session yielded 3-20 simultaneously recorded motor units, which were then pooled across sessions. This also involved pooling across muscles. For present analyses, such pooling is desirable because the task is performed using both the *triceps* and *anterior deltoid*, and stimulation of the recorded region of M1 activated these muscles. Consistent with the fact that these muscles are used together, the pooled motor-unit population still had low dimensionality.

We have previously documented that motor-unit recruitment is flexible ^28^. Recruitment reflects force profile, muscle length, and (when stimulating cortex) the precise stimulation site. The example motor-unit in Fig. 3 shows some flexibility: it becomes less active, for the same force, at higher frequencies. Motor-unit recruitment flexibility is one reason why the dimensionality of the motor-unit population is higher than one (the principal additional contribution being non-linearity – a motor unit typically remains silent below its force threshold). Yet while motor-unit flexibility exceeds traditional expectations, motor units never showed the extreme flexibility of M1 neurons. M1 neurons had a variety of correlations with force (positive, zero, and negative) and the sign and magnitude of these correlations often differed (for the same neuron) across conditions. This was not true of motor-units: if a motor-unit had a sufficiently low threshold (and thus was generally active across conditions) its activity consistently correlated positively with force.

### Estimating Control Strategies via Autoregressive Exogenous Input (ARX) Models

To estimate the control policy deployed in each condition, we modeled the force output on each trial using an autoregressive model with exogenous inputs (ARX): as described in Wasicko et al. ^76^. This model captures how force at each time point depends on its own recent history, the recent history of errors, and on an internal template. Specifically, we modeled the change in force at time *t* as: 

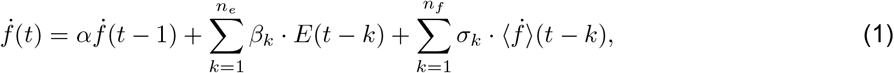

where 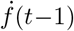 is the most recent derivative of force, *E*(*t*) is the force-tracking error, and 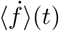 is the derivative of force averaged across all trials with the same force profile. The first of these terms yields dependence on recent history (providing smoothness). The second term provides dependence on error (the difference between force and target force). The third term acts as a template. For example, if the third term dominates, the predicted derivative of force can be entirely determined by the typical (trial-averaged) derivative. The parameters *α, β*, and *σ* were fit using ridge regression. Control policy distance (see below) was based on how well these parameters, when fit to one condition, accounted for data from another condition. Model hyperparameters (ridge penalty, number of parameters (*n*_*e*_ and *n*_*f*_)) were selected via cross validation on a held-out dataset (20% of the trials for each condition).

### Generalized Linear Models

We asked how well single-neuron (and single-motor unit) activity was explained via reference to force (Supp. Fig. 2). In addition to force, we also considered related variables that a feedback-controller might use. Specifically, we asked how well single-trial activity was accounted for by four explanatory variables: instantaneous force, error, and their derivatives. Analysis was performed at the level of single trials. Spikes were counted in 1 ms bins, and behavioral data were sampled at 1 kHz. The spike count was then fit via a Binomial Generalized Linear Model with a logit link function: 

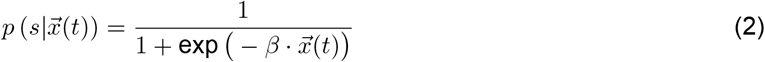

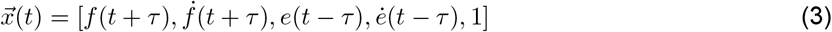

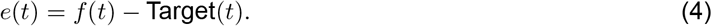

The lag, *τ*, was set to 80 ms for all units. We used this lag to reflect two assumptions. First, any dependence of neural activity on error (the visual difference between Pac-Man’s height and the dot height) will necessarily occur after the error itself. When visual events impact M1 activity (e.g. the response to target onset in a reaching task), they typically do so with a lag of 60–80 ms, hence this choice. Second, there will necessarily be a lag between neural activity and the downstream force it will cause. This lag is due both to axonal conduction and to the delay between electrical activation of muscle fibers and force generation. Estimates of this delay vary, but are typically in the 50-100 ms range. For the sake of simplicity we used a unified value of 80 ms for both lags (with opposite signs). Results were extremely similar across a reasonable range of choices (50–100 ms). Goodness of fit was calculated on a held out dataset using two metrics: bits per spike and *R*^2^. Bits per spike was calculated as, 

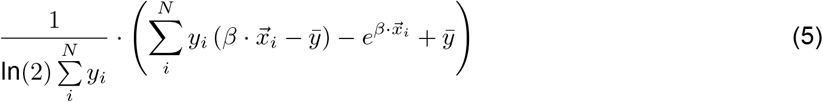

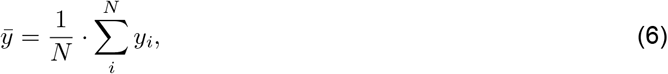

where *N* is the total number of spikes emitted by the neuron and *y*_*i*_ denotes the *i*^th^ spike of the neuron.

The second metric, *R*^2^, was calculated as the *R*^2^ between the predicted PSTH (after filtering and averaging over trials) and the empirical PSTH.

### Subspace Alignment Index

Activity for a given condition (or condition-group) occupies a set of dimensions – i.e. a subspace. To quantify the degree to which two conditions (or condition-groups) *A* and *B* share the same subspace, we employed the subspace alignment index ^29^: 

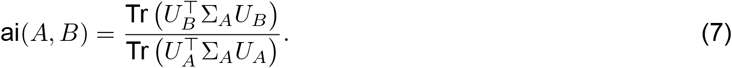

Here, *U*_*A*_ and *U*_*B*_ are matrices whose columns contain the top *k* principal components computed for condition *A* and *B* respectively. The matrix Σ_*A*_ is the covariance of the data during condition *A*. Thus the numerator is the variance from condition *A* captured by PCs from condition *B*, while the denominator is the variance from condition *A* captured in its own PCs (which by construction capture the maximum variance of any *k*-dimensional subspace). This index ranges from 0 (no alignment) to 1 (perfect alignment).

The value of the alignment index is typically robust across a range of values of *k*, but becomes sensitive at the extremes. At one extreme, if *k* equals the total number of recorded neurons, alignment will trivially be one for all comparisons. At the other extreme, if *k* = 1, alignment may be low if the first dimension is not shared across conditions, even if other dimensions are. Given that we are interested in situations where alignment may sometimes become low, a conservative choice is to use a reasonably large value of *k*. We thus set *k* = 15, as this exceeds the number of dimensions (roughly 10–12) typically assumed by the constrained-manifold hypothesis. Empirically, this value was well-suited to the present data, as it revealed both situations where activity was well-aligned and situations where it was not. That said, other reasonable choices produced nearly identical results (see below). For the motor-unit population, we set *k* = 5 to be conservative in the opposite direction (to avoid overestimating alignment).

Another reasonable way of choosing *k* is as a percentage of the size of the recorded population. To explore this choice, we set *k* to be 3% of the total number of recorded neurons for that population (resulting in *k* ≈ 30 for the M1 populations and *k* = 3 for the motor-unit population). Results were nearly identical to those for the original analysis in Supp. Fig. 7. We also performed a control that matched both *k* and population-size for M1 and motor-unit populations (Supp. Fig. 8). The M1 population was downsampled (many times) to match the size of the motor-unit population, and we used the same value of *k* = 3 for both (equivalent to 3% of neurons as described above). Alignment remained far lower for M1.

We performed two styles of alignment-index-based analyses: static and rolling. Static analyses considered activity over a large, static, epoch of time (the whole condition, or multiple similar conditions), which is how the alignment index has traditionally been computed. Static analyses employed a cross-validated approach (described below) that slightly improves the estimate of alignment by removing bias due to sampling error, while also symmetrizing the alignment metric. Rolling analyses considered a sliding window of activity in one of two conditions. For rolling analyses, we were less concerned with interpreting the exact magnitude, and more concerned with estimating its time-course. Furthermore, rolling analyses are intrinsically asymmetric (the sliding window applies to one condition but not the other). Rolling analyses thus use the standard non-cross-validated approach as in ^29^.

### Rolling Subspace Alignment Index

The rolling alignment index quantifies the degree to which activity, within a sliding window during a ‘test’ condition, is captured in a subspace computed from the full duration of activity during a ‘reference’ condition (Supp. Fig. 9 and Supp. Fig. 10). Let *A*_test_(*t*) correspond to data from within a sliding window of the test condition, from *t* − 500 ms to time *t*. Let *B*_ref_ correspond to data from the full duration of the reference condition. The rolling alignment, at time *t*, was then 

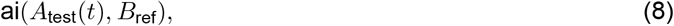

using the definition of the alignment index above. Confidence intervals (equivalent to standard errors of the mean) were computed using a bootstrap, by resampling across neurons.

### Cross-validated Subspace Alignment Index

Trial-by-trial noise can reduce the measured alignment between subspaces. Suppose conditions *A* and *B* were identical; i.e. evoked the same firing rates in all neurons. For a finite number of trials, the alignment index would still be slightly less than unity, because the principal components found for condition *B* will not be exactly identical to those for condition *A*, and the latter by definition capture maximal variance. This bias, though usually quite small, can cause alignment to be slightly underestimated. Given the nature of the hypotheses that were tested, we wanted to be careful not to underestimate alignment. We therefore computed a cross-validated subspace alignment index, adapted from the standard approach ^29^.

We constructed two independent sets of PSTHs for each condition by splitting trials into two non-overlapping groups. This resulted in two PSTHs per condition: one for group 1 trials and the other for group 2 trials. Using these splits, we constructed two separate subspaces for each condition group: *A*^1^ and *A*^2^. The cross-validated alignment index between condition groups *A* and *B* with (*A* ≠ *B*) was then computed as: 

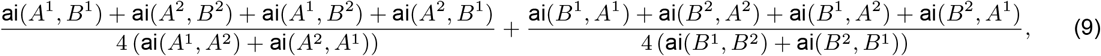

where each term uses the standard alignment index described above. The cross-terms in the numerator average the empirical alignment across trial partitions: all combinations of *A* with *B* and *B* with *A*. The terms in the denominator provide an empirical estimate of the upper limit of alignment given sampling error; in the limit of infinite data, each of these terms would simply be unity.

### Sparse Component Analysis

A prediction of the flexible repertoire hypothesis is that population activity should involve distinct sets of factors (occupying distinct dimensions / subspaces) at different moments. Factors should thus be sparse in some ways (at a given moment, many factors may be inactive) but not others (factors may be reused across conditions if subskills are reused). Aspects of these predictions were confirmed by the alignment-based analyses in Supp. Fig. 9 and Supp. Fig. 10. For example, subspaces associated with slow increasing and slow decreasing ramps were sequentially reused during the 0.25 Hz sine condition but not the during the chirp, which instead reused the 1, 2, and 3 Hz subspaces. While these analyses demonstrate compositional reuse of subspaces, they relied on supervision. Specifically, they required an informed guess regarding which putative subskills were likely relevant when. Furthermore, although the alignment index has the advantage of simplicity (it is high when dimensions are reused and low when they are not), it does not allow one to directly inspect how the underlying factors evolve with time. This matters, because many of the predictions of the flexible-repertoire hypothesis pertain not only to when factors will be active, but to their time-course when they are active. For example, if a group of factors becomes active only for fast sines, that group should instantiate dynamics consistent with generating a fast sine, and should form a basis for decoding force during fast sines.

To estimate factors in an unsupervised manner, we applied Sparse Component Analysis ^47^ (SCA) to both M1 and motor-unit populations. Like PCA, SCA seeks to find latent factors that form a basis for the observed neural responses. That is, each neuron’s response should be well-approximated as a linear sum of the factors (e.g. Supp. Fig. 13). However, unlike PCA, SCA does not attempt to concentrate variance in the initial factors. The SCA cost function encourages maximization of variance captured across all factors, but not variance captured by (for example) the first factor. Additionally, the SCA cost function encourages the identified factors to be sparse if possible; the cost function is reduced if the basis includes factors that are active at some moments and not others. SCA can be viewed as a variant of PCA with an *L*_1_ penalty applied to the projections to encourage sparsity.

SCA functions as a form of hypothesis-guided dimensionality reduction ^77^ that is ideally suited to the competing hypotheses being evaluated. Under the traditional constrained-manifold hypothesis, population activity should be accounted for by relatively few factors, most of which should be active across conditions (i.e. should be non-sparse). Because the SCA cost function is dominated by the goal of capturing variance, it cannot find sparse factors unless they form a natural basis set. Thus, under the constrained-manifold hypothesis, SCA should identify relatively few factors, most of which will be non-sparse. That SCA behaves this way for a low-dimensional non-sparse population response is confirmed by application to the motor-unit population (Supp. Fig. 14).

Under the flexible-repertoire hypothesis, accounting for the population response should require many factors, many of which will be sparsely active. Because the SCA cost-function encourages a sparse factor basis, if such a basis is not found, the flexible-repertoire hypothesis can be rejected. If a high-dimensional sparse basis is found, that would tend to support the flexible-repertoire hypothesis. One would then evaluate the behavior of the factors. There are many ways for factors to be sparse, and the flexible-repertoire hypothesis makes specific predictions (e.g. regarding dynamics during fast sines, as described above). Thus, a proper test of the flexible-repertoire hypothesis requires asking not only whether sparse factors exist, but also whether their activity, across time and condition, accords with predictions.

Two key hyperparameters govern the behavior of SCA: the total number of latent factors and the strength of the *L*_1_ penalty, *λ*. Results were robust across a wide range of total factors. We set the number of total factors to be 3% of the number of recorded units, yielding 3 for the motor-unit population and ≈ 30 for the neural populations. This choice resulted in factors that captured the majority of population variance for both M1 and motor-unit populations. Additionally, the vast majority of individual factors (e.g. those plotted in Fig. 5 and Supp. Fig. 11) captured clear structure. If we increased the number of requested factors further, the additional identified factors tended to be low-variance (*<* 1%) and lack clear structure (i.e. they appeared to be dominated by sampling error). Thus, given the limitations imposed by population-size and trial-count, the number of factors whose time-course can be accurately estimated appears to be ≈ 30. This does not rule out the possibility that more factors could be accurately estimated with larger population sizes or higher trial-counts. At the other extreme, it was easy to avoid the regime where too few factors were requested, because this resulted in too little population variance being captured.

To test the robustness of our findings to the sparsity hyperparameter, we systematically varied *λ*. Doing so also acts as a further test of competing hypotheses. When *λ* is zero, the SCA cost function is more similar to that for PCA, and does not seek sparsity. At very large values of *λ*, the sparsity term dominates and reconstruction error will increase (the identified factors will be a poor basis). One can thus ask what occurs as *λ* is steadily increased. Different hypotheses make different predictions. Under the flexible-repertoire hypothesis, there exists a natural basis set where factors are somewhat sparse. Consequently, as *λ* increases, factor sparsity is predicted to increase before there is a meaningful rise in reconstruction error; the estimated factors will simply align better with the true sparse factors. This was indeed what occurred for both M1 populations (Supp. Fig. 15B,C). Under the constrained-manifold hypothesis, there does not exist a natural basis set of sparse factors. Thus, as *λ* increases, reconstruction error will rise before there is much change in sparsity. Put differently, increasing *λ* will increase sparsity only slightly, at the expense of a sizable increase in reconstruction error. This was indeed what occurred for the motor-unit population (Supp. Fig. 15A).

The analysis described above (and shown in Supp. Fig. 15) used two metrics of sparsity: excess kurtosis and latent sparsity. Excess kurtosis is defined as: 

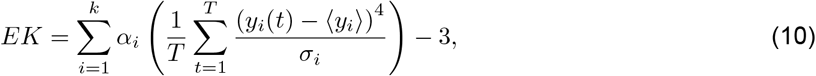

where *y*_*i*_ is the *i*^th^ factor, *σ*_*i*_ is its standard deviation, and *α*_*i*_ is the ratio of the variance explained by the *i*^th^ factor relative to that explained by all *k* factors. Excess kurtosis measures how much higher the average kurtosis of the latent factors is than expected from a Gaussian distribution. A high excess kurtosis indicates the data is more heavy-tailed (thus sparser) than expected under a Gaussian null model. Latent sparsity is defined as: 

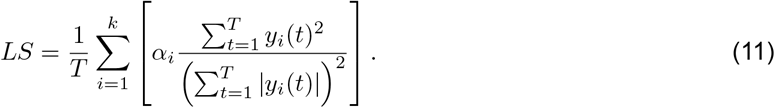

Latent sparsity also measures how sparse the latents are but is bounded between 0 (not sparse) and 1 (maximally sparse).

### Control Policy Distance

To quantify the similarity between control policies inferred from different conditions, we define a control policy distance metric based on generalization error. Intuitively, if two conditions share a similar control strategy, then a policy learned from one should perform well when applied to data from the other.

Let *π*_*i*_ = *π*(· ; *α*_*i*_, *β*_*i*_, *σ*_*i*_) denote the control policy inferred from condition *i*, parameterized by the fitted autoregressive (*α*), feedback (*β*), and feedforward weights (*σ*) as described in Equation (1). Let 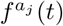 be the force generated for condition *a* on trial *j* and let 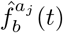 be the predicted force trajectory for condition *a* on trial *j* using the control policy from condition *b*. The latter is obtained by integrating the predicted force derivative generated by *π*_*a*_ over the all timesteps. We define the control policy distance between conditions *a* and *b* as: 

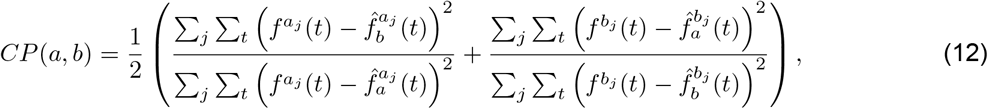

This metric thus reflects the relative decrease in prediction accuracy when applying policies across conditions, normalized by each policy’s performance on its own data. A value of 1 reflects perfect generalization, while larger values reflect poor transfer across conditions.

### Force Distance

Force distance was computed as the difference between the average force value, 

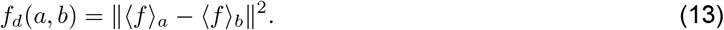

### Directionality Distance

The directional distance between two conditions was computed as the difference between the average of the sign of their derivatives, 

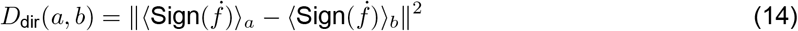

### Neural Distance Prediction

We computed, for each condition, its activity centroid: the location, in full-dimensional space, of neural activity averaged over time. Any two conditions thus had a distance between them, which was small for some conditions and large for others. To assess the factors that determined centroid separation, we fit (squared) centroid distance as a linear function of the three behavioral distances above: control policy distance, force distance, and directionality distance (each also expressed as a squared distance). To aid interpretation of regression weights, we normalized each feature to have unit variance over the dataset.

### Exponential Fitting Procedure

To quantify the relationship between neural centroid distance and sub-space alignment, we modeled their relationship using a logistic function of the form: 

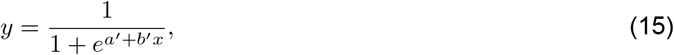

where *y* is the subspace alignment index, *x* is the (unsquared) centroid distance between two conditions, and *a*^*′*^, and *b*^*′*^ are fitted parameters.

To estimate this model, we applied a logit transform to the alignment index: 

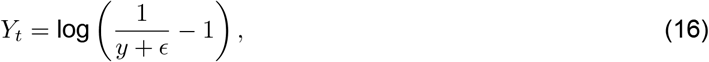

where *ϵ* is a small constant added to ensure the argument of the log is always positive. We then fit a linear model to the transformed data: 

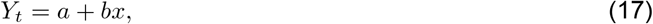

where *a* and *b* correspond to the intercept and slope, respectively.

After fitting in the transformed space, we inverted the transform to obtain the final exponential predictions in the original space: 

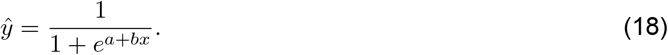

The significance of the relationship was assessed using the *p*-value of the slope from the linear model. The overall goodness-of-fit was computed using the untransfomed alignment index and the nonlinear model predictions.

### Linear Dynamics System Fitting

To capture the low-dimensional dynamics of population activity during each condition (Supp. Fig. 12), we fit a linear dynamical system (LDS) model to neural data using Dynamic Mode Decomposition ^78^.

We first reduced neural activity to its top 10 principal components. This was done separately for each condition. We then fit a 10-dimensional LDS of the form: 

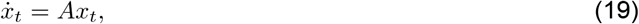

where *x*_*t*_ is the projection of the neural state vector onto the top principal components at time *t* and *A* is a learned matrix describing the linear dynamics.

DMD estimates the best linear approximation of the dynamics, *A*, by relating consecutive datapoints: 

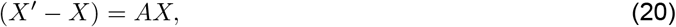

where *X*, and *X*^*′*^ contain the state vector projections at times *t* and *t* + 1, respectively. We computed *A* using the standard DMD solution: 

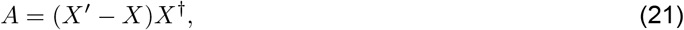

where *X*^*†*^ denotes the Moore-Penrose pseudoinverse of *X*.

Dimensionality was fixed at 10 to capture the dominant low-rank dynamics while avoiding overfitting. While 10 dimensions would clearly be insufficient (under the flexible-repertoire hypothesis) to capture global dynamics, the present analysis simply seeks to find a local linear approximation that fits each condition on its own. This respects the hypothesis that different conditions involve activity in different regions of state-space, with different local linear approximations to the dynamics.

### Analysis of Decoding Accuracy versus Encoding Strength

We used ridge regression to fit linear decoders to reconstruct force based on neural activity, using the model, 

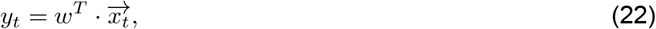

where the solution is given by, 

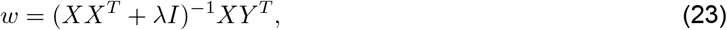

where *λ, X*, and *Y* are the ridge penalty, the matrix of neural (or motor unit) responses, and the vector containing the values of force across time and conditions.

We quantified two aspects of this fit: decoding accuracy and encoding strength. Decoding accuracy was the *R*^2^ value of the fit to *y*_*t*_, computed for held-out data (trials not used during training). Parameters were fit using firing-rate and force estimates from one partition, and generalization performance was tested for another partition. Encoding strength quantified the proportion of population variance, within 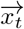, that is captured by the decoding axis *w*, computed as 

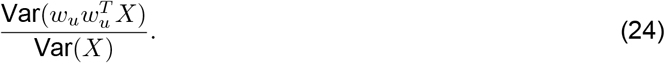

where *w*_*u*_ = *w*/∥*w*∥. For a population of pure force-tuned neurons, both decoding accuracy and encoding strength would be near unity. However, it is also possible for decoding accuracy to be high while encoding strength is low. For example, this would occur in a population where force is only one of many signals reflected in single-neuron responses.

Estimates of decoding accuracy and encoding strength depend on *λ*. As *λ* increases, stronger regularization encourages greater encoding strength. Decoding accuracy will also tend to initially improve, because regularization improves generalization. However, if the true encoding strength is low (i.e. if force is a low-variance signal within the population) increasing values of *λ* will soon cause decoding accuracy to decline. This occurs because the estimated decoding dimension (which is encouraged by regularization to be high-variance) can no longer align with the true (low-variance) dimension that encodes force. At very high values of *λ*, decoding will necessarily be poor simply because the weights *w* are forced towards zero.

Some analyses (Supp. Fig. 16A) consider the full relationship, as *λ* increases, between decoding accuracy and encoding strength. We also summarized this relationship by identifying the ‘knee’ where accuracy began to fall as encoding strength increased (Supp. Fig. 16B,C). To do so we chose the largest value of *λ* where decoding accuracy remained above 80% of its maximal value. Doing so estimates the strongest that encoding strength can plausibly be, given that force decoding must be accurate.

The above analysis was applied to trial-averaged responses. This choice reduces the impact of sampling error, and thus allows decoding accuracy to potentially approach unity (which indeed it sometimes did) even if encoding strength is low. Analysis was repeated using the derivative of force as the target variable (Supp. Fig. 16D-F). We also repeated analysis for M1, after restricting the population to the most force-tuned neurons (Supp. Fig. 17).

### Neural Dissimilarity

We computed the neural dissimilarity of the response of the *i*^th^ neuron, across contexts *a* and *b*, using, 

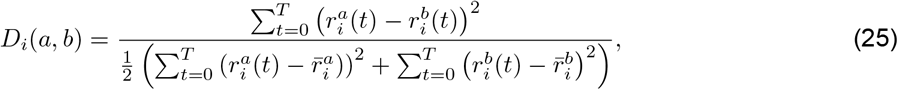

where 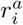 and 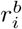 are the mean firing rate of neuron *i* in the two contexts, and *t* indexes across all times within a condition. A dissimilarity of zero indicates an identical mean response across contexts, while a value larger than 1 indicates that across context differences exceed the within-context variance of activity. Dissimilarity was computed per neuron and per force-profile, to respect the fact that a neuron’s response might differ little across contexts for some profiles, and a lot for others (this was indeed true for monkey I, who performed a larger variety of force profiles). Cumulative distributions considered dissimilarities for each neuron and force profile.

### Cross-context Similarity

Cross-context similarity is a population-level metric that quantifies the similarity of neural population activity between contexts *a* and *b*. It is closely related to one minus the Neural Dissimilarity metric, but operates on projections onto the PCs rather than on individual-neuron responses. We first performed PCA jointly on neural activity pooled across both contexts. We then projected activity from each context onto the top *k* PCs, yielding context-specific trajectories **x**^*a*^(*t*) and **x**^*b*^(*t*) in a shared *k*-dimensional subspace. We set *k* = 30 to capture sufficient overall variance while still rejecting noise due to sampling error. Similar results were obtained across a range of reasonable choices. Similarity was then computed as: 

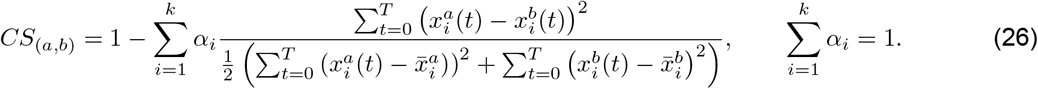

The ratio computes, for each PC, the normalized squared difference in activity. That difference is then weighted by *α*_*i*_, which is proportional to the variance captured by the *i*^*th*^ dimension, normalized to sum to unity for the number of PCs considered.

### Removing Translation Between Contexts

To determine how much of the observed cross-context difference in population structure could be explained by simple mean shifts (translations), we computed similarity in a manner that ignores differences in the overall mean across contexts: 

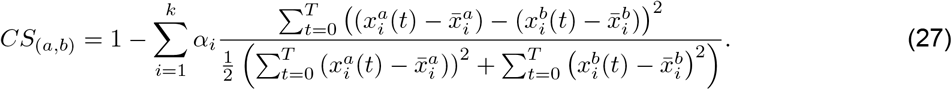

Comparing the original cross-context similarity to the translation-removed cross-context similarity thus quantifies the extent to which cross-context differences reflect a uniform shift versus changes in the shape of the trajectory.

### Removing Rotation Between Contexts

After removing translation (see above), we asked to what degree similarity could be further increased by a rotation of the data within the PCs. Towards this end we solved the following Procrustes problem: 

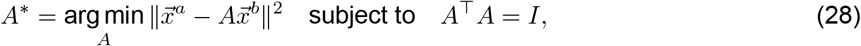

where 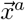 and 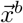 are the demeaned population trajectories, projected onto the top PCs (*k* = 30 as above) for contexts *a* and *b*, respectively. Solving this problem yields the optimal orthogonal transformation, *A*^*^, that best aligns the trajectories of context *b* with those of context *a*. We then applied the optimal rotation *A*^*^ to the trajectories from context *b*, and recomputed the cross-context similarity as previously described.

## Supplemental Figures

**Supplementary Figure 1.**
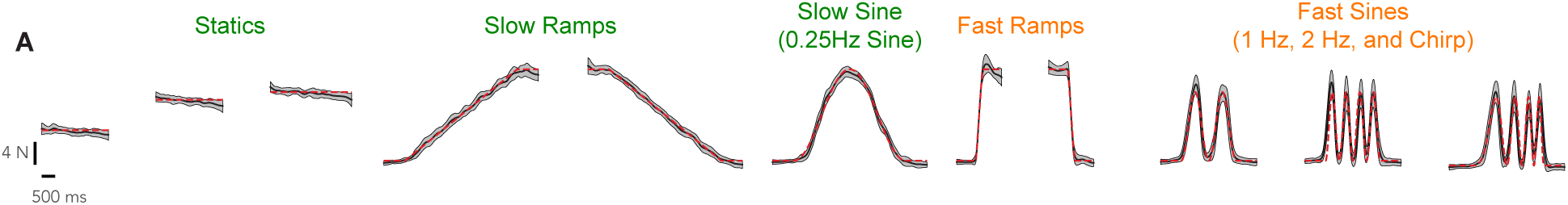
Monkey I Behavior. (**A**) Same as for Fig. 2B but for Monkey I. Conditions are the same with two exceptions. First, the 2 Hz sine started and ended at the bottom, rather than the top. Second, Monkey I could not consistently perform the 3 Hz sine condition with high accuracy, and it was thus not included in his condition-set.

**Supplementary Figure 2.**
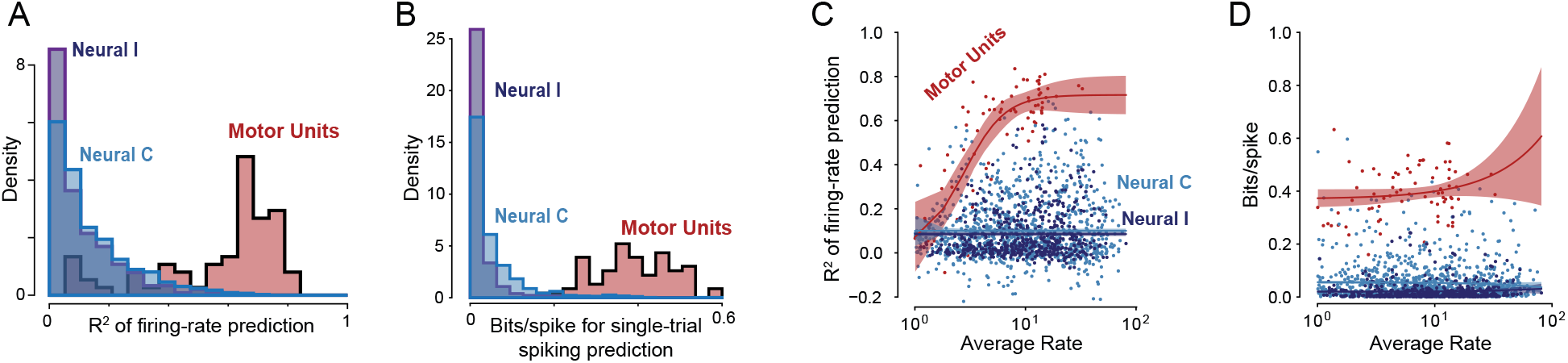
Activity of single motor units, but not M1 neurons, strongly reflects force and related behavioral variables. We asked whether single-neuron spiking was a function of features a feedback controller might reasonably be expected to use: force, error, and their derivatives. To do so, we trained generalized linear models (GLMs) to predict spiking activity using these features. Model performance was evaluated on held-out trials. To quantify performance, we used both a cross-validated *R*^2^ metric (computed between predicted and actual trial-averaged rates) and a single-trial bits/spike metric. (**A**) Distribution of *R*^2^ values, shown separately for neural (M1) and motor-unit populations. Motor unit activity was well predicted by the GLMs, consistent with the role of motor units in force production. M1 neuron activity was predicted poorly. This agrees with what can be seen by inspection: it was rare for the rate of an M1 neuron to consistently encode force or its derivative. (**B**) Distribution of normalized bits/spike values across the same three populations in (A). Again, motor-unit activity was accounted for quite well, but M1-neuron activity was not. (**C**) A potential concern regarding the previous analysis (in panels A and B) is that differences in fits between M1 and motor-units might be secondary to differences in mean rate. To address this, for each neuron (blue) or motor unit (red), we plotted the *R*^2^ of model fit is versus firing rate. Lines are fit using a Hill model; shaded regions denote 95% confidence intervals for model fit, computed via bootstrap. For a given rate range, motor-unit fits were typically much better. (**D**) Same as (C) but fit quality is assessed in terms of normalized bits/spike. Again, for a given rate range, motor-unit fits were typically much better. These analyses demonstrate that single-neuron firing rates in M1 rarely correlated well with force (or related variables). This does not imply that force cannot be decoded accurately from the M1 population response – it could, as will be documented below.

**Supplementary Figure 3.**
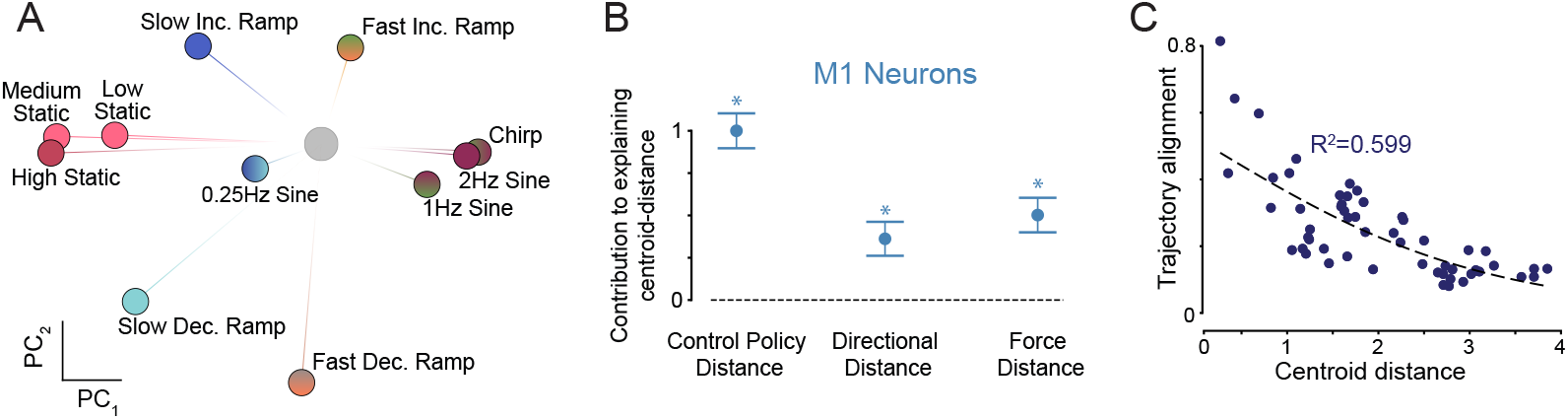
Same as Fig. 4 but for Monkey I. Panel C of Fig. 4 already included results for monkey I; thus, that analysis is not repeated here. (**A**) Same analysis as in Fig. 4B. As for monkey C, centroids were well-separated. Aspects of centroid organization are also similar. For example, conditions requiring closed-versus open-loop control are well-separated. (**B**) Same analysis as in Fig. 4E. (**C**) Same analysis as in Fig. 4F. This analysis tests the flexible-repertoire prediction that, when two conditions use the same or similar subskills, centroid distance should be small and subspace alignment should be high. When they use very different subskills, centroid distance should be large and subspace alignment should be low. Consequently, centroid distance and subspace alignment should be negatively correlated when compared across condition pairs. To quantify the relationship between these two variables, we fit a logistic model by applying a logit transform and fitting a linear model to the transformed data (Methods). The significance of the slope was assessed in the transformed domain (linear-regression *p*-value), while the model fit (*R*^2^) considered the final nonlinear predictions. Analysis was across all pairs of single conditions. Each dot corresponds to one condition pair and plots the alignment index between those conditions versus their centroid distance. Centroid-distance was computed in full-dimensional neural space. Solid line shows the fit. As for monkey C, there was a negative relationship between centroid distance and subspace alignment (*p <* 10^−5^).

**Supplementary Figure 4.**
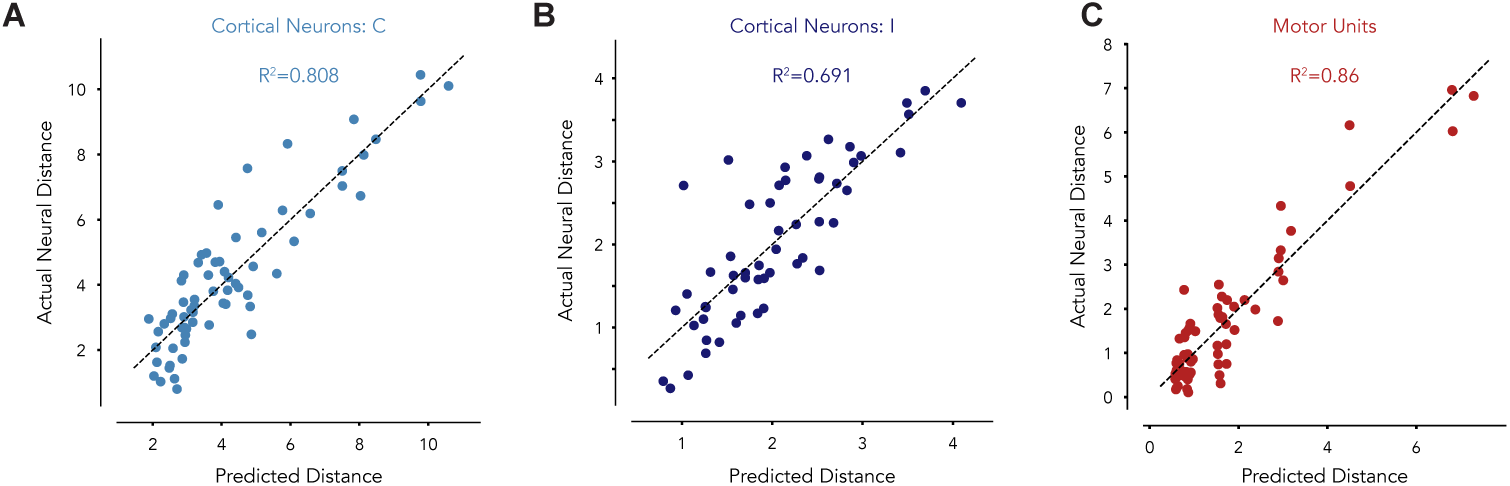
Neural centroid distances were predicted well using a linear model. This analysis provides the basis for the analyses in Fig. 4E and Supp. Fig. 3B. Centroid distances were predicted based on three behavioral distances. These plots document those fits: they show actual versus predicted centroid distance for each pair of conditions. Fig. 4E and Supp. Fig. 3B document the contribution, to these fits, of the different behavioral distances. (**A**) Analysis for M1 centroids for Monkey C. Each dot corresponds to one pair of conditions, and plots the actual distance between their centroids versus the predicted distance. Actual distances were computed in the full-dimensional space after each neuron’s rate was soft-normalized (see Methods). Distances thus have arbitrary units (rather than spikes/s). Predictions used a linear model that employed Force Distance, Directionality Distance, and Control Policy Distance as features (see Methods for details on how these were computed). (**B**) Same analysis for M1 centroids for Monkey I. (**C**) Same analysis for motor-unit centroids for Monkey C.

**Supplementary Figure 5.**
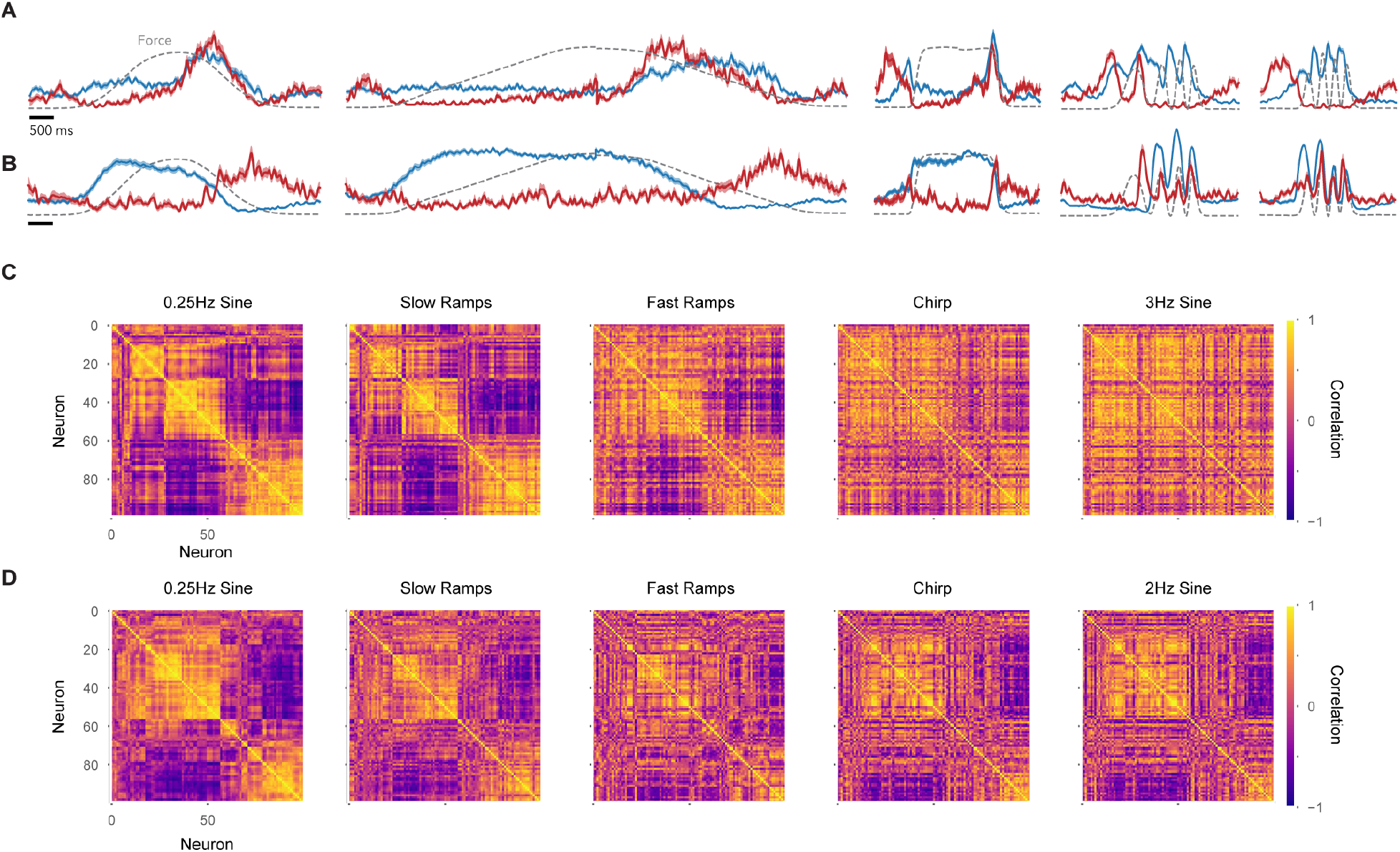
Neuron-neuron correlations, when compared between conditions, are sometimes very similar and sometimes very different. A prediction of the flexible-repertoire hypothesis is that, when conditions differentially recruit subskills, neural activity will occupy non-identical sets of dimensions. A change in neural dimensions is equivalent to a change in neuron-neuron correlations. For example, suppose two neurons are both positively influenced by factor A, captured in dimension 1. Suppose that factor B, captured in dimension 2, has a negative influence on one neuron and a positive influence on the other. When a subskill primarily recruits factor A, the neurons will be positively correlated. When a subskill primarily recruits factor B, the neurons will be negatively correlated. Thus, before considering subspaces, we first examine neuron-neuron correlations. To aid visualization when viewing matrices of correlations, we restricted this analysis to the top 100 highest SNR neurons. (**A**) Average firing rate (envelopes show SE) for two example Monkey C cortical neurons, across various conditions. Dashed-black traces plot mean force. These neurons were positively correlated during low-frequency conditions (e.g. *r* = 0.83 at 0.25 Hz), but negatively correlated at high frequencies (e.g. −0.53 at 3 Hz). (**B**) Two additional example neurons from Monkey C showing the opposite pattern: *r* = −0.74 at 0.25 Hz and *r* = 0.72 at 3 Hz. These examples reflect a broader trend in the cortical population: neuron–neuron correlations shift relatively little between some pairs of conditions but rather a lot between others. In both example pairs, their correlation remained similar during the 0.25 Hz sine and the slow ramps, but reversed sign for the chirp. (**C**) Neuron-neuron correlation matrices for Monkey C. Neuron order was determined by a hierarchical clustering algorithm, which was used to highlight correlation structure during the 0.25 Hz sine condition. By sorting neurons to highlight structure, one can see when and whether this structure is maintained across other conditions. Correlations are shown for 5 conditions. For ‘Slow Ramps’ and ‘Fast Ramps’, we concatenated data across both directions (as illustrated in panel A). The correlation structure evident during the 0.25 Hz Sine is maintained during the Slow Ramps. This is consistent with a variety of other observations, made below. When comparing the 0.25 Hz sine with the fast ramps or the chirp, some broad structure is preserved but much is different (many rows/columns have changed their sign). Correlation structure is very different for the 3 Hz Sine. (**D**) Same analysis but for Monkey I. As above, aspects of neuron-neuron correlations are preserved across conditions (aspects of the block structure remain) while others change (many individual rows and columns change completely). As noted above, Monkey I did not perform the 3 Hz sine.

**Supplementary Figure 6.**
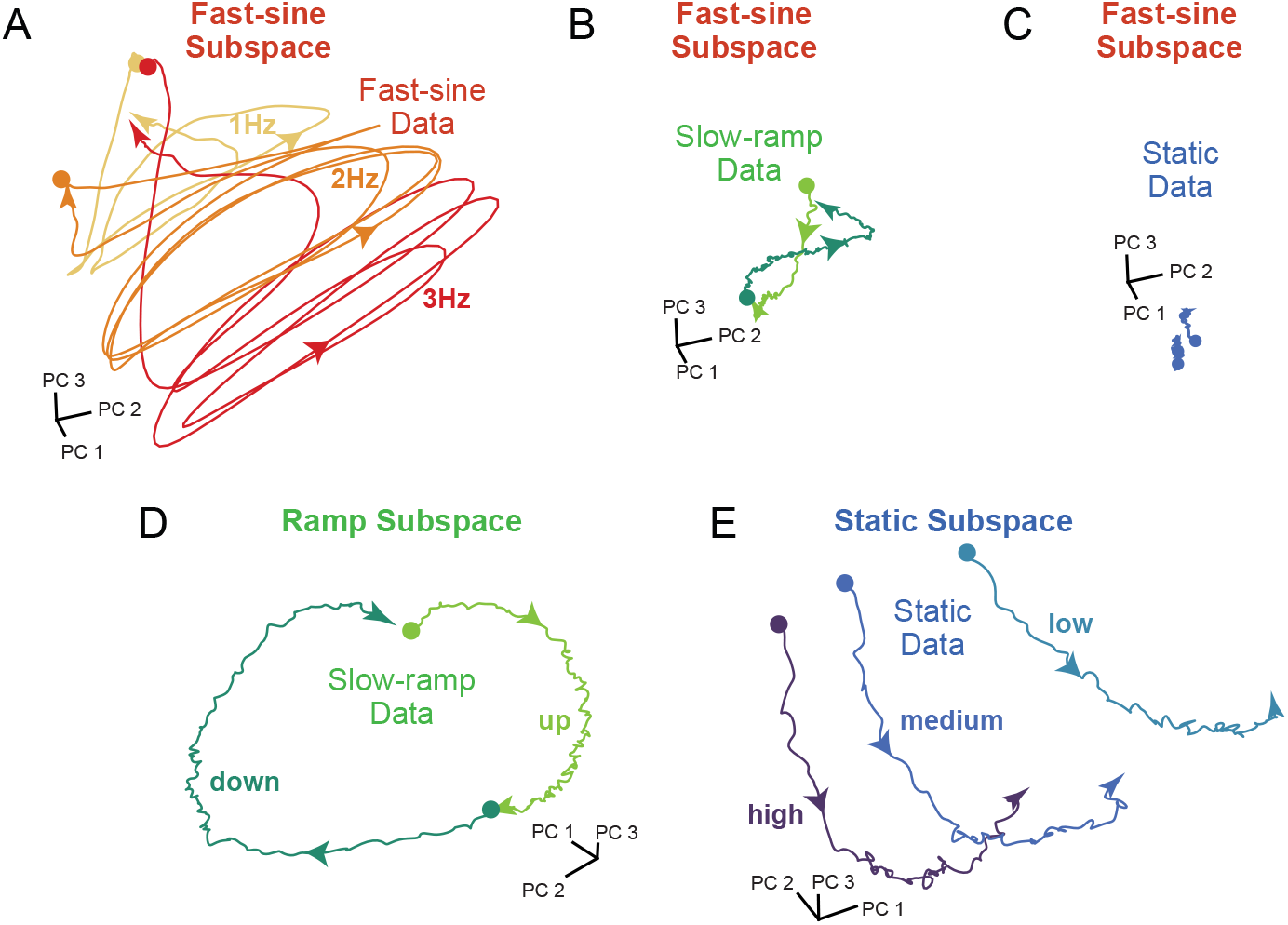
Neural trajectories occupy different subspaces across conditions. The subspace-alignment metric, used in the main text, quantifies how well activity during one condition (or set of conditions) is captured by neural dimensions found for a different condition (or set of conditions). Here, instead of computing variance captured, we directly examine activity by plotting neural trajectories. Each panel plots the projection of neural population activity, for one set of conditions, into a subspace defined by the top three PCs computed from either the same conditions or from different conditions. All data are for Monkey C. (**A**) Neural trajectories for fast (1, 2, and 3 Hz) sine conditions, projected into a ‘fast sine subspace’ based on data from those same conditions. (**B**) Neural trajectories for the two slow ramps, projected into the fast sine subspace. Even though many neurons responded during the slow ramps, trajectories in this subspace are small (i.e. low variance) with unclear structure. (**C**) Neural trajectories for the three static conditions, projected into the fast sine subspace. Trajectories are barely visible; they are quite low-variance in this subspace. (**D**) Neural trajectories for the two slow ramps, projected into a ‘ramp subspace’ based on data from all ramp conditions (fast and slow). Trajectories are larger, with clearer structure, in this subspace. (**E**) Neural trajectories for the three static conditions, projected into a ‘static subspace’ based on data from those same conditions. Again, structure is present that was lost when trajectories were projected into the fast sine subspace.

**Supplementary Figure 7.**
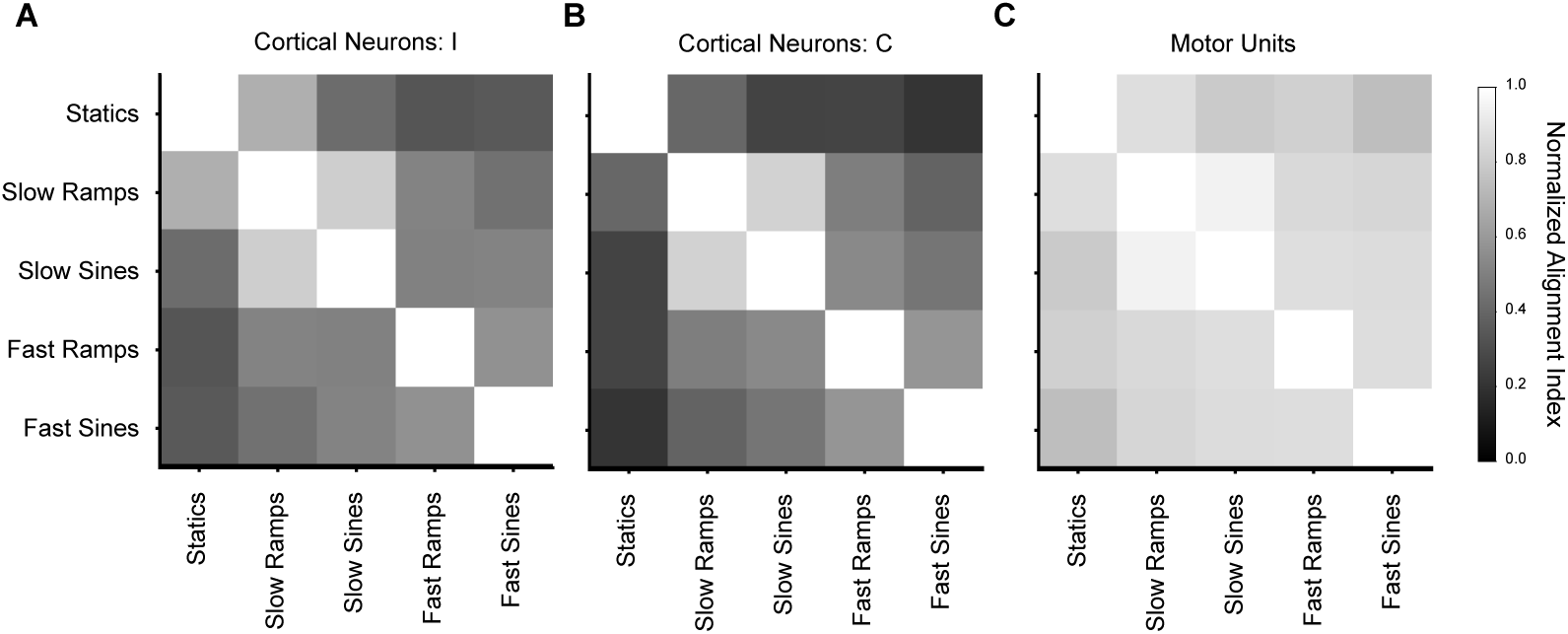
Subspace alignment across condition-groups. Each matrix plots the alignment index for multiple comparisons. The alignment index ranges from zero to one, and asks how well activity for one group of conditions is captured by the top PCs from a different group of conditions, relative to how well activity would have been captured in its ‘own’ PCs. We used the following groups. Statics: low, medium and high statics; Slow Ramps: slow increasing and decreasing ramps; Slow Sine: 0.25 Hz sine; Fast Ramps: fast increasing and decreasing ramps; Fast Sines: 1, 2, and 3 Hz sines (3 Hz for monkey C only) and the chirp. Each condition-group involved a similar range of forces and (except for the statics) included both increasing and decreasing forces. Alignment was computed in a cross-validated fashion (using trial-partitions) and normalized to be unity when comparing a condition with itself (Methods). For M1, analysis was performed using a subspace dimensionality of 15 – i.e. slightly larger than the total dimensionality typically proposed under the constrained-manifold hypothesis. This choice is conservative; if the constrained-manifold hypothesis is correct, we wish to ensure dimensionality is not underestimated. For the motor-unit population, we wished to be conservative in the opposite direction, and thus chose a subspace dimensionality of five. To test sensitivity to these choices, analysis was rerun (for both M1 and motor-unit populations) using dimensionality equal to 3% of the size of the recorded population (*≈* 30 for M1, 3 for motor-units). Results were extremely similar. (**A**) Analysis for the M1 population recorded from Monkey I. (**B**) Analysis for the M1 population recorded from Monkey C. (**C**) Analysis for the motor-unit population.

**Supplementary Figure 8.**
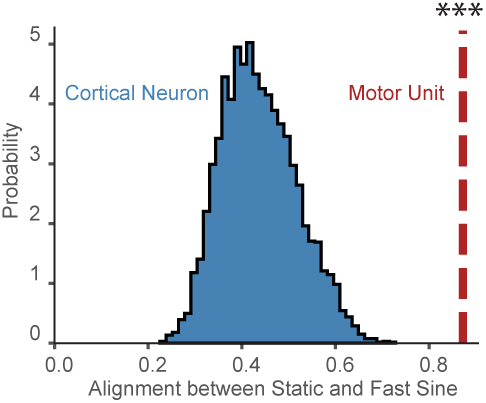
Alignment Index Differences Are Not Due to Population Sizes. Results above demonstrate that alignment is consistently high for the motor-unit population, but becomes low, for some comparisons, for the M1 populations. Here we explore that result again after matching both the population size and the value of *k* (the number of dimensions used to compute alignment). We repeatedly sub-sampled the M1 population of Monkey C to match the size of his motor-unit population. We set *k* = 3 (i.e. 3% of the population size) for both M1 and motor units. (**A**) Distribution of alignment-index values when resampling the M1 population (blue), compared to the value for the motor-unit population (red line). The distribution of indices for M1 is significantly lower than, and non-overlapping with, that for the motor-unit population (*p <* 0.001). This analysis computed the alignment between the ‘static’ conditions (low, medium, and high) and the ‘fast sines’ (1, 2, and 3 Hz Sine conditions and the Chirp). Repeating analysis for all comparisons confirmed the original results; for comparisons where alignment was low in Supp. Fig. 7, alignment remained low after matching population size.

**Supplementary Figure 9.**
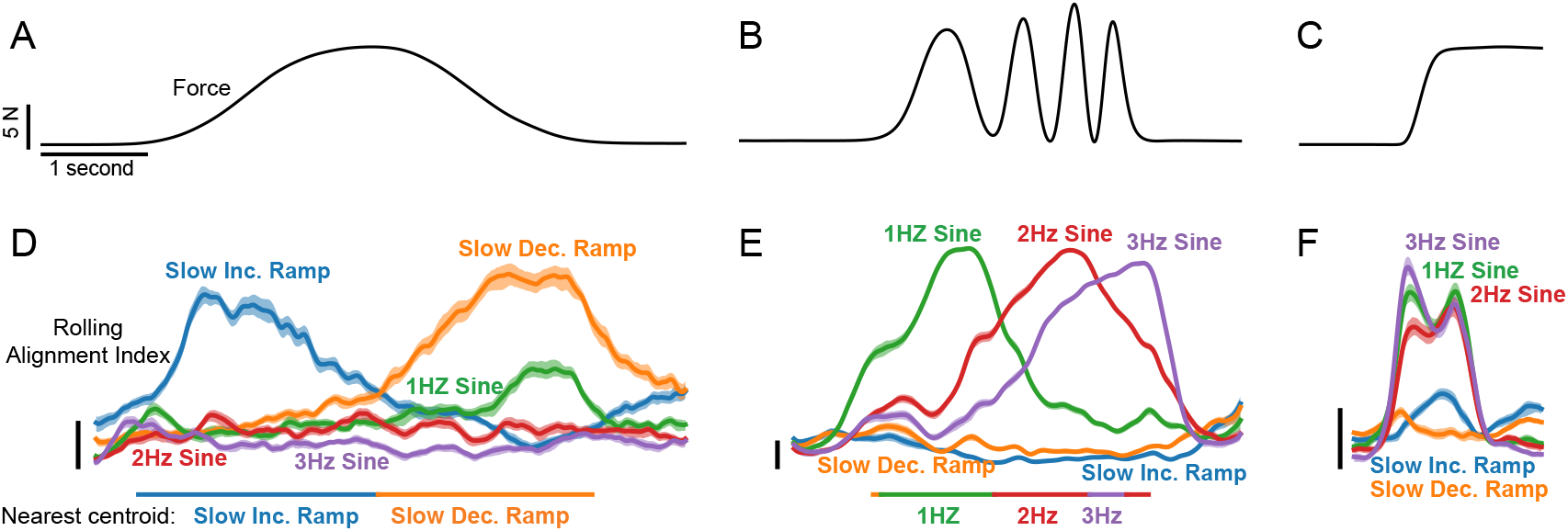
Neural dimensions are reused in specific ways and at specific times (Monkey C). The analysis of subspace alignment above (main text and Supp. Fig. 7) reveals that the M1 population response sometimes reuses dimensions across groups of conditions. For example, alignment was higher when comparing the slow 0.25 Hz sine with slow ramps, and lower when comparing the slow 0.25 Hz sine with fast sines. The flexible-repertoire hypothesis predicts that this should happen for a specific reason: some conditions may reuse subskills compositionally. If so, there should be two forms of specificity: which subspaces align and when they align. Here we test this prediction by computing the alignment index, between various single conditions, as a function of time. We consider three ‘test’ conditions that might potentially involve compositionality: the 0.25 Hz sine, chirp, and fast-increasing ramp. For each, we ask how well the neural trajectory for that condition (in a 500 ms sliding window) is captured in subspaces for five ‘reference’ conditions: the three faster sines and two slow ramps. Subspaces for the reference conditions were computed across the full duration of the reference condition. We also computed the centroid for each reference condition (as in the main text), and determined if and when the trajectory for the test condition neared one of these centroids. (**A**,**B**,**C**) Average force, as a function of time, during the three test conditions. (**D**) Alignment when the test condition was a 0.25 Hz sine. Alignment indices are shown as a function of time. Each trace corresponds to one reference condition (as labeled). Envelopes denote 95% confidence intervals computed using a studentized bootstrap. Vertical scale bar denotes an alignment index of 0.1. The colored bar at the bottom indicates, across time, the reference centroid to which activity was closest. During the rising phase of the 0.25 Hz sine, activity occurs within the subspace used during the slow rising ramp, and comes nearest that centroid. During the falling phase, activity occurs within the subspace used during the slow falling ramp, and comes nearest that centroid. This effect is predicted if the 0.25 Hz sine condition reuses, in order, the subskills used during the slow rising and falling ramps. (**E**) Alignment when the test condition was the chirp. Activity aligns, in order, with subspaces for 1, 2, and 3 Hz sines. (**F**) Alignment when the test condition was a fast increasing ramp. Activity occurs, simultaneously, within subspaces for the 1, 2, and 3 Hz sines. This may potentially relate to the broad frequency content present in a rapid ramp. Activity did not near any single centroid, and thus no bar is shown at bottom.

**Supplementary Figure 10.**
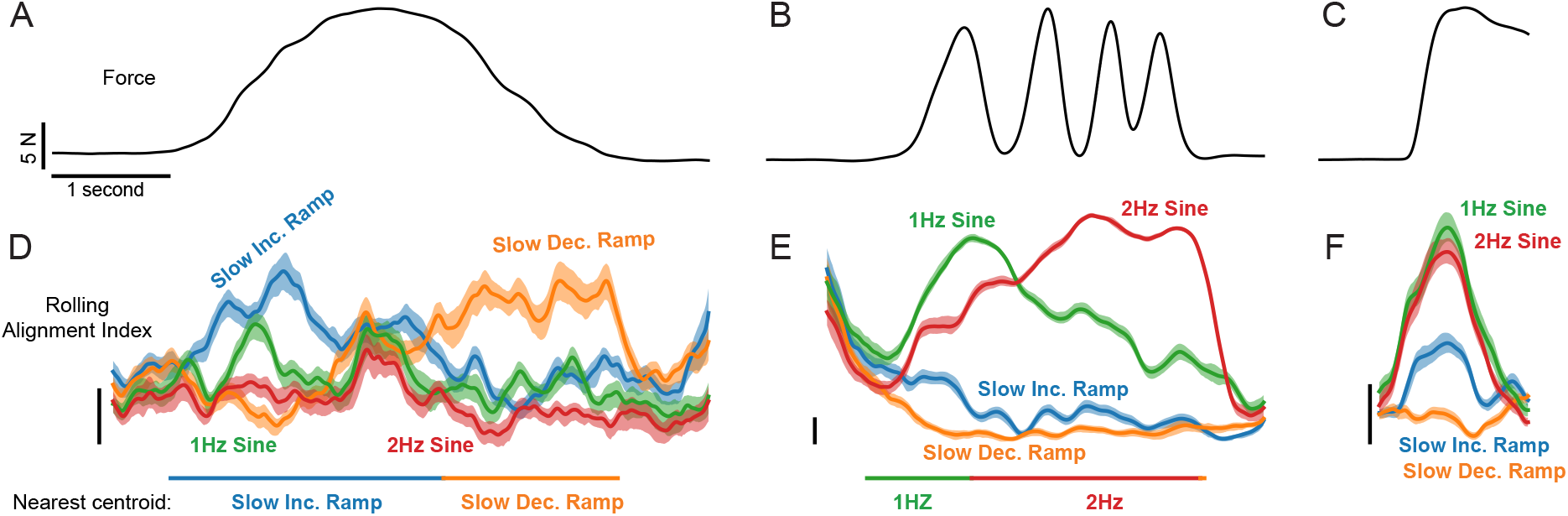
Neural dimensions are reused in specific ways and at specific times (Monkey I). Same analysis as in Supplementary Figure 9, except for Monkey I. The 3 Hz sine is not used as a reference condition because Monkey I did not perform this condition.

**Supplementary Figure 11.**
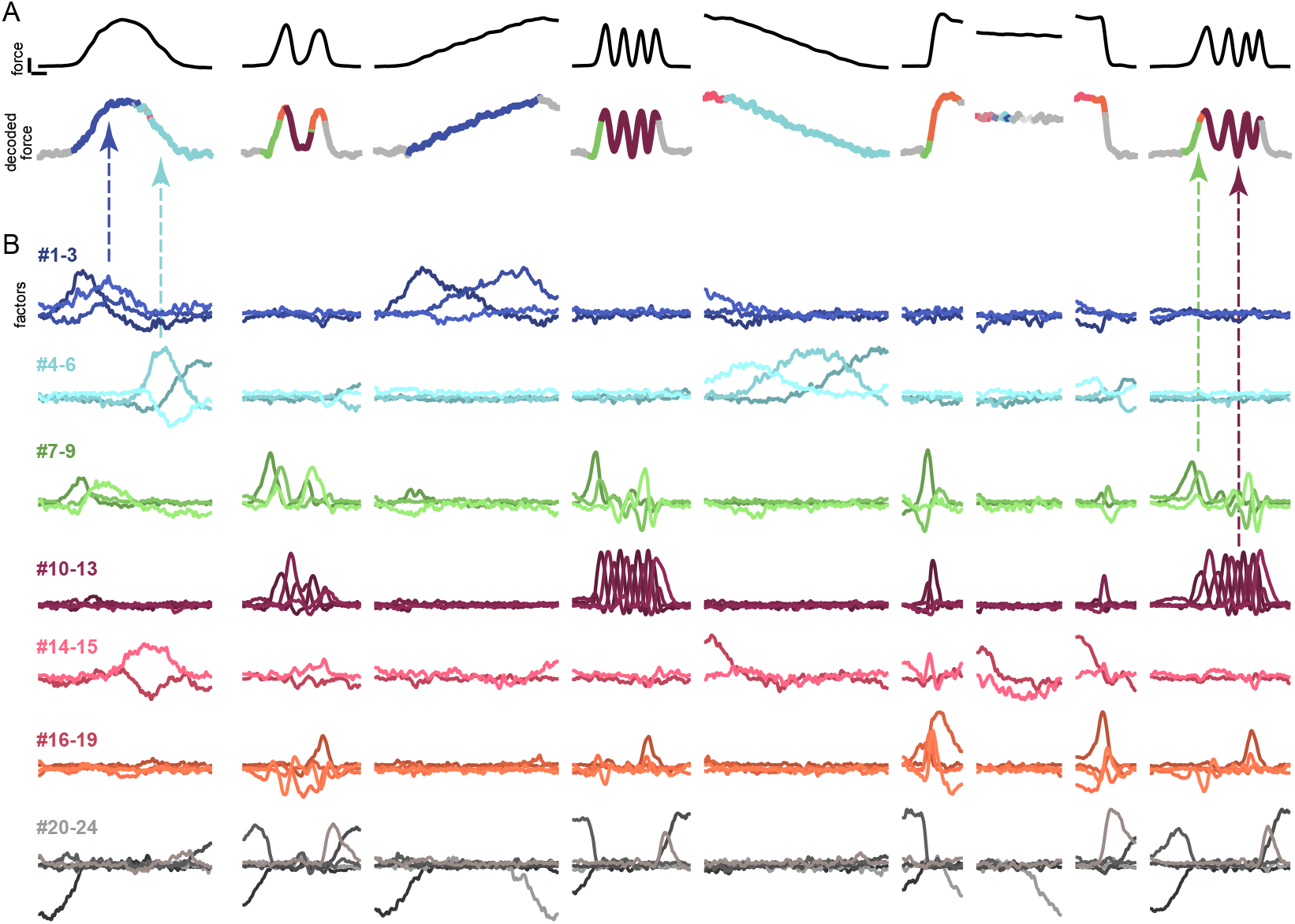
Same as Fig. 5 but for Monkey I. Factors are grouped similarly, but not identically, to those for monkey C. It is anticipated that monkeys may differ somewhat in how they ‘divide’ the overall task into subskills. That said, most divisions, and instances of compositionality, were similar. For example, as for monkey C, the 0.25 Hz sine reuses, in order, the blue factor group (also used during the slow rising ramp) then the cyan factor group (also used during the slow falling ramp). This recapitulates, at the level of the factors themselves, the effect documented via subspace alignment in Supp. Fig. 10D. Similarly, the chirp uses the green factor group (also used during the 1 Hz sine) before the purple factor group (also used during the 2 Hz sine), in agreement with Supp. Fig. 10E.

**Supplementary Figure 12.**
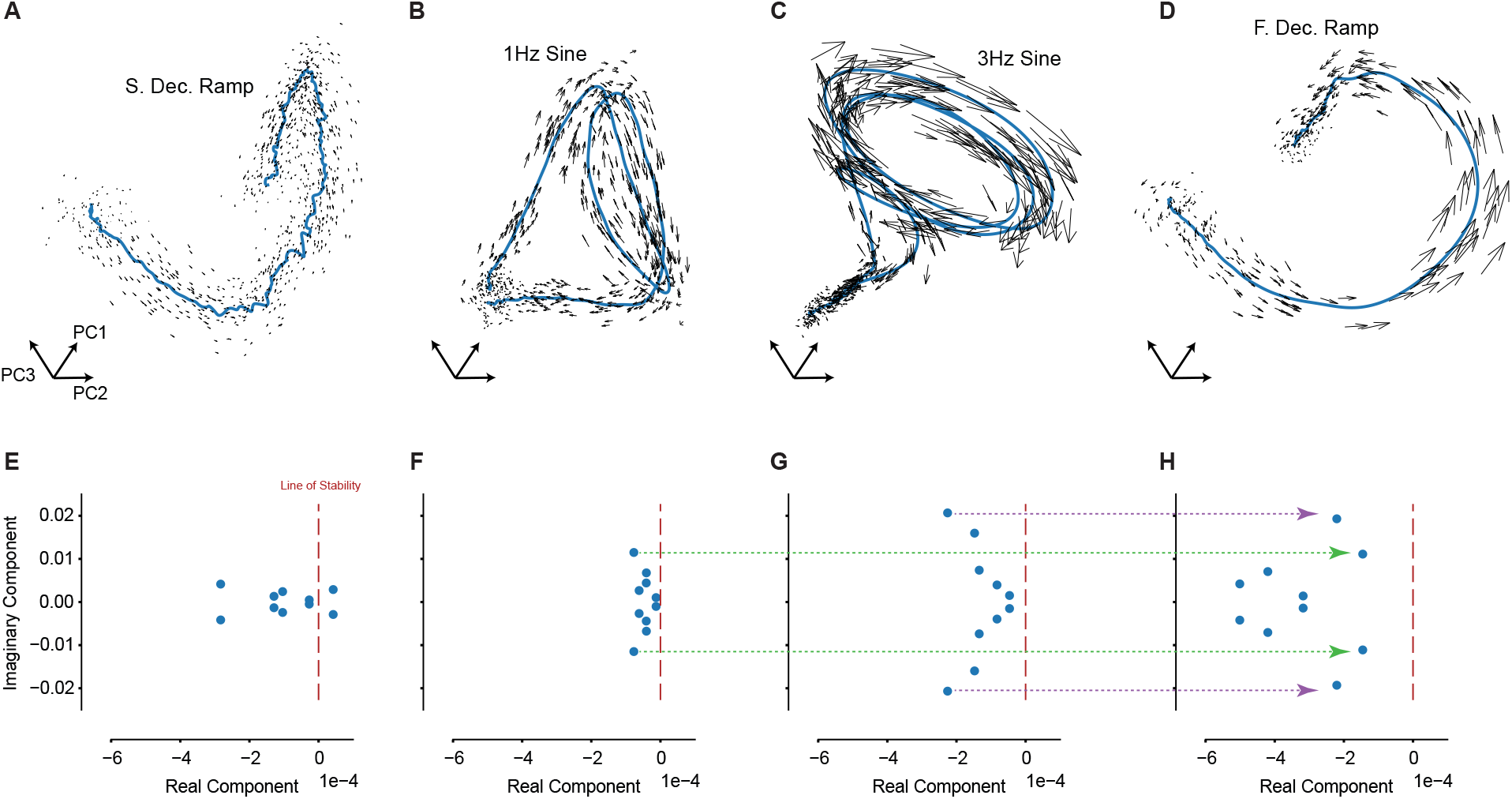
Neural Dynamics Vary Across Conditions. The flexible-repertoire hypothesis posits that different subskills leverage different state-space locations and different dimensions (i.e. different factors). The purpose of doing so is to allow dynamics to differ across subskills while maintaining low trajectory tangling. A key prediction is thus that conditions should display a variety of neural dynamics, each appropriate to the condition in which they are deployed. We fit Linear Dynamical System (LDS) models to estimate the local flow-field for each condition (Methods). Each condition was fit separately, with fits based on activity in the dimensions used by that condition. If eigenvalues of the LDS models differ, then local dynamics differ. This approach is conservative; it is possible for dynamics to differ even when eigenvalues do not. Data are for monkey C, to allow comparison with factor-properties in Fig. 5. (**A**) Neural activity (blue) and the inferred flow-field (black arrows) for the slow decreasing ramp condition. The model was fit to the neural trajectory in the top 10 PCs. The plot shows projections, for both the trajectory and the flow-field, into the top 3 PCs. The flow-field is shown only locally near the trajectory. (**B-D**) Same as (A) but for other conditions. Note that each panel involves activity in a different global location (Fig. 4, Supp. Fig. 3) and in different dimensions (Supp. Fig. 6, Supp. Fig. 7). (**E**) Eigenvalue spectrum for the LDS model fit to the data in (A). The dashed red line denotes the maximum stable eigenvalue. (**F-H**) Same as (E) but for other conditions. Eigenvalue spectra reflect properties visible in the active factors (Fig. 5). For example, the slowly evolving cyan factor-group in Fig. 5 is active during the slow decreasing ramp, in agreement with the eigenvalues in panel E, which lack any fast oscillatory structure (the imaginary component is small). The oscillatory green factor-group is active during the 1 Hz sine, in agreement with the eigenvalues in panel F, which have a larger imaginary component. The higher-frequency oscillatory purple factor-group is active during the 3 Hz sine, in agreement with the higher-magnitude imaginary component in panel G. During the fast decreasing ramp, both green and purple factor groups are briefly active in Fig. 5. In agreement, the eigenvalue spectrum in panel H includes complex eigenvalue pairs that roughly match the higher frequencies from panels F and G (highlighted by arrows).

**Supplementary Figure 13.**
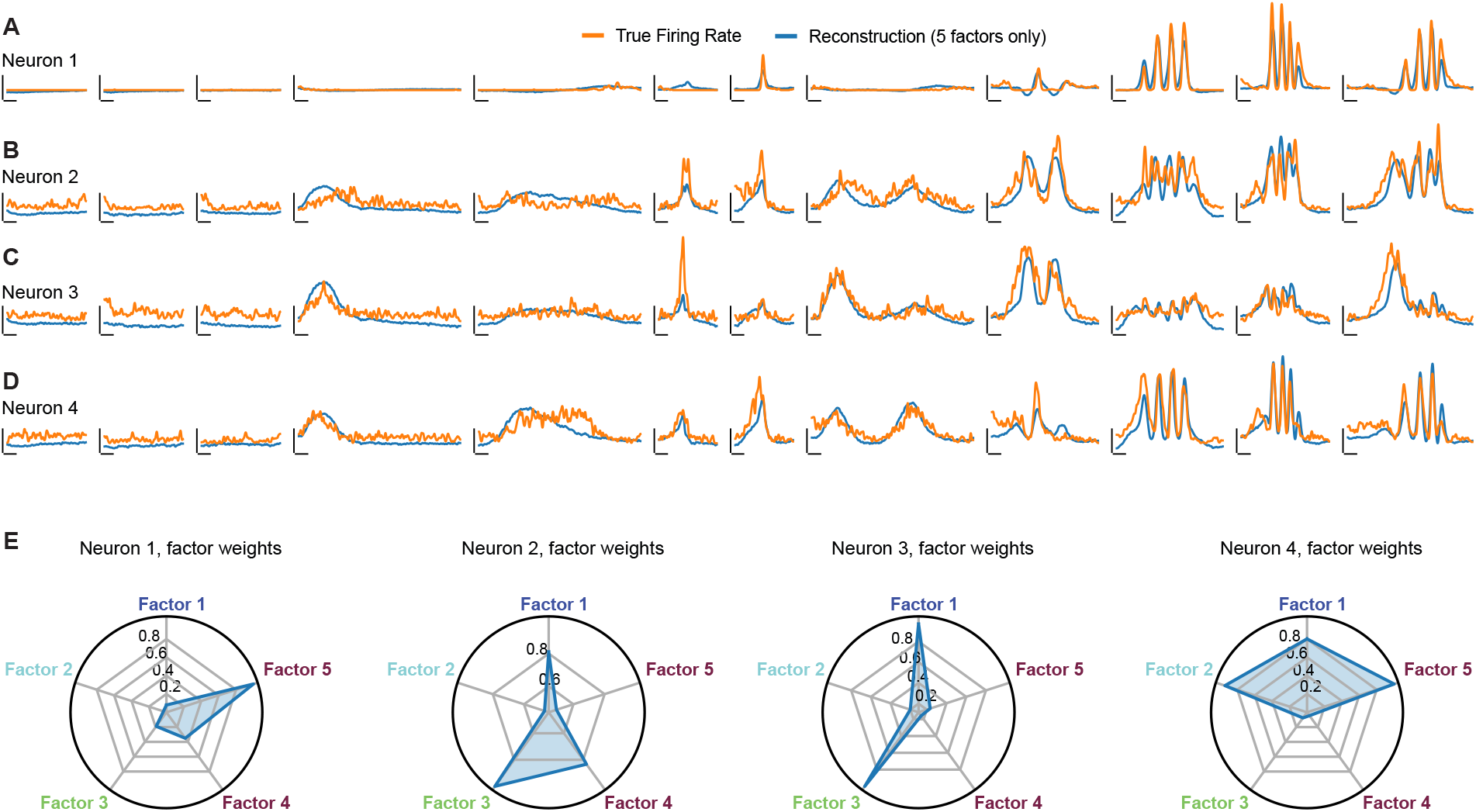
Examples of how single-neuron firing rates reflect factors. Factors are population-level signals, yet also form a basis set for single-neuron responses. Here we illustrate this for four example neurons, by reconstructing each neuron’s rate from five example factors. For ease of illustration, we intentionally chose example neurons that strongly reflected the same set of five factors. These factors were from the blue, cyan, green, and purple groups in Fig. 5. (**A**) Mean firing rate of one neuron (orange) and its reconstruction (blue) via a weighted sum of the five factors. (**B-D**) Same but for different example neurons. (**E**) Normalized contribution of each factor to the reconstructions above. Some neurons (e.g. neuron 1) had rates that primarily reflected one factor, and were thus quite sparsely active across conditions. Other neurons had rates that reflected multiple factors.

**Supplementary Figure 14.**
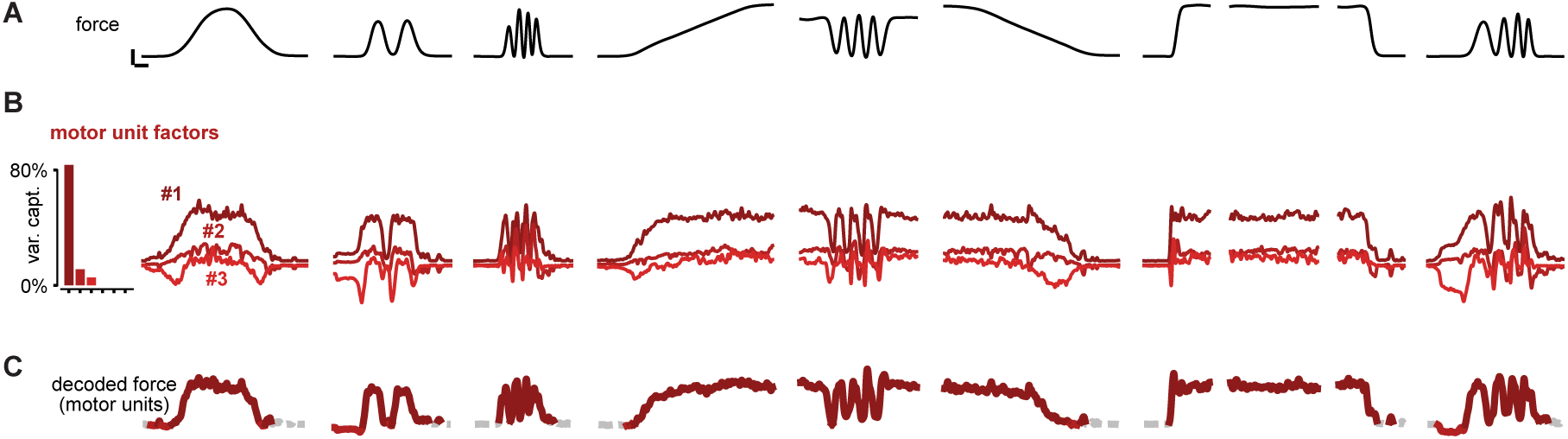
SCA factors for the motor-unit population. Analysis parallels that for the M1 populations in Fig. 5 and Supp. Fig. 11. (**A**) Trial-averaged forces during different conditions. (**B**) Projection of motor-unit activity onto 3 SCA dimensions, yielding the factors. Unlike for M1, one factor dominated. Of the variance explained by SCA, *>* 80% is captured by the first factor (inset shows, for each factor, the percent of total captured variance that it accounted for). (**C**) Decoded force, based on a linear combination of the factors. The color of decoded-force trace indicates, at each moment, which factor made the largest contribution to decoding. Decoding relied almost entirely on the dominant factor. The trace is dashed gray at moments where no factor made a contribution (this occurred when the motor unit population was nearly silent).

**Supplementary Figure 15.**
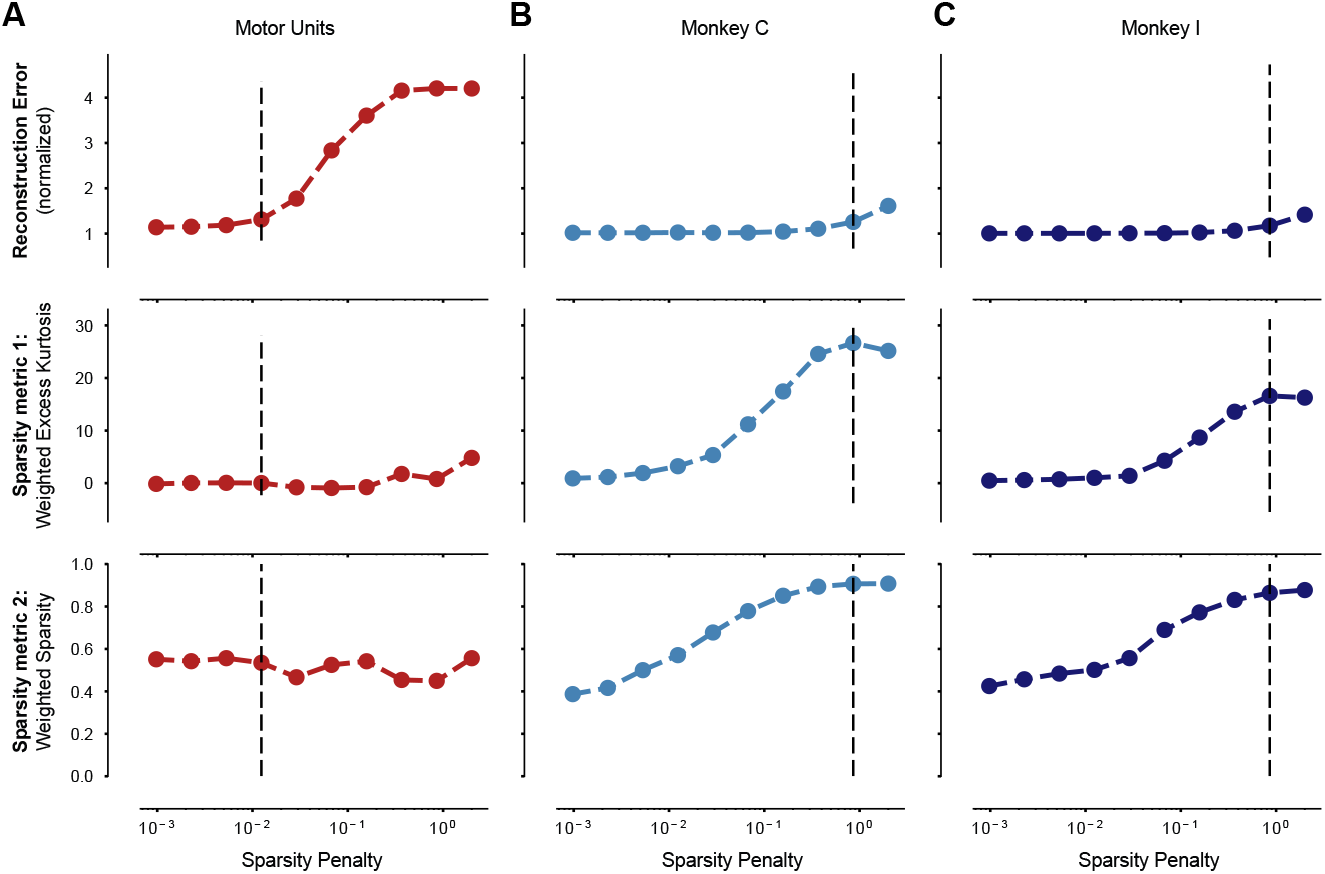
For M1, but not the motor-units, SCA identified sparse factors and did so with little loss in overall variance captured. The SCA objective function includes a ‘sparsity penalty’ that encourages sparsity and a term that encourages variance captured (across all factors). This introduces a potential tradeoff between identifying factors that are sparse and identifying factors that capture variance. In situations where factors truly are sparse, this tradeoff should be minimal: there should be values of the sparsity penalty that identify sparse factors with little loss of variance captured (and thus minimal increase in reconstruction error when reconstructing individual-neuron responses from the factors). In situations where the factors are truly not sparse, this should not be true: increasing the sparsity penalty should produce a considerable increase in reconstruction error, and yet the estimated factors should still be relatively non-sparse. To explore, we swept the sparsity penalty and measured: reconstruction error (top row), excess kurtosis (a measure of sparseness, middle row) and weighted sparsity (a different measure of sparseness, bottom row). Reconstruction error was normalized by the PCA reconstruction error (PCA provides the minimum reconstruction error of all linear dimensionality reduction methods). Excess kurtosis asks how different the distribution of each factor’s values are from a Gaussian. Weighted sparsity was measured using the activity participation ratio. (**A**) Analysis for the motor units. Increasing the sparsity penalty caused a roughly four-fold increase in reconstruction error, but negligible increase in sparsity. Put differently, it was not possible to identify factors that were both sparse and provided a good basis for reconstructing motor-unit activity. The dashed black line corresponds to the sparsity regularization used to generate Supp. Fig. 14. This value was chosen so that the SCA factors explained *>*75% as much variance as the same number of principal components. (**B**) Same but for the M1 population recorded from Monkey C. As the sparsity penalty increased, factor sparsity increased considerably while reconstruction error remained low. Reconstruction error eventually increased, but only at high penalty values. Thus, it was relatively easy to identify factors that were both sparse and provided a good basis set for single-neuron responses. The dashed black line corresponds to the sparsity regularization used to generate Fig. 5. This value was chosen so that SCA factors explained *>*95% as much variance as the same number of principal components. Note that this choice was intentionally conservative: for the M1 population, sparse factors were found despite minimal increase in reconstruction error. This contrasts with the motor-unit population, where sparse factors were not found despite a larger increase in reconstruction error. (**C**) Same but for the M1 population recorded from Monkey I. The dashed black line corresponds to the sparsity regularization used to generate Supp. Fig. 11, and used the same criterion as in (B).

**Supplementary Figure 16.**
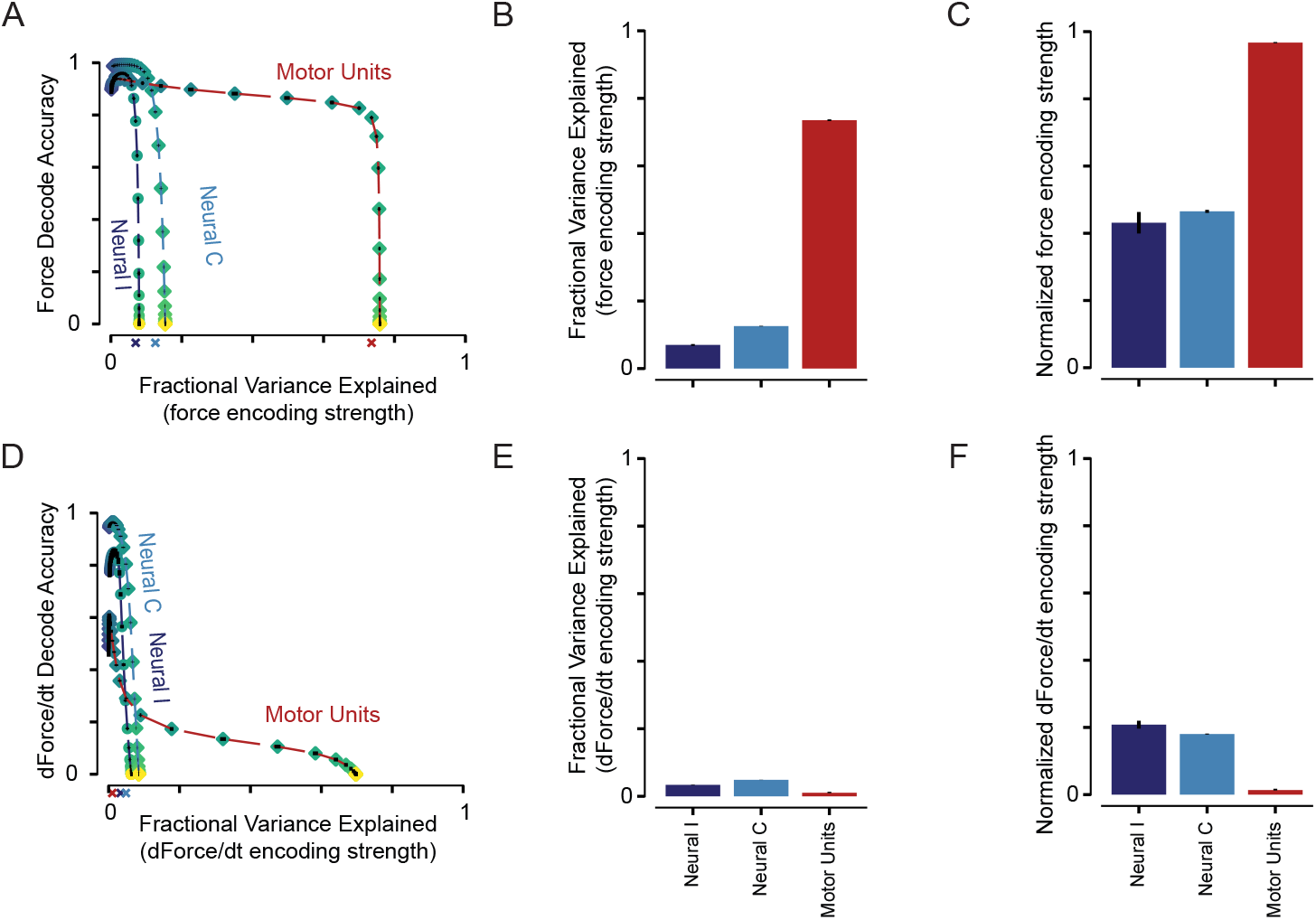
Force is accurately decoded from both M1 and motor-unit populations, but the decoding dimension captures a large fraction of the population’s response structure only for motor units. While the spiking activity of individual M1 neurons did not reliably reflect force, prior work argues that force (or, equivalently, commands for muscle activity) are encoded at the population level. The flexible-repertoire hypothesis similarly assumes that there exists a readout dimension where the projection of population activity provides descending commands. Thus, under both traditional hypotheses and the flexible-repertoire hypothesis, it should be possible to find a dimension where the projection of population activity yields accurate force decoding across all conditions. The Pac-Man task is very ‘force-centric’: many parameters, including Pac-Man’s height, mirror force. Accurate force generation is also the fundamental goal of the task. Thus, under most traditional hypotheses, the force-decoding dimension should capture a large fraction of response structure. Put differently, there should exist a readout dimension where both decode accuracy (the force variance explained) and encoding strength (the neural response variance explained) are high. Under the flexible-repertoire hypothesis, the readout dimension is impacted by many factors (none consistently resembling force) and will thus explain only a small percentage of population response structure; encoding strength will never be high. We thus obtain the following prediction: encouraging the readout dimension to capture more response variance should impair decoding accuracy for M1 but not the motor units. To test this, we used ridge regression to predict force from population activity. Decode weights were learned based on trial-averaged responses from one partition of trials. Accuracy was then assessed when force was decoded based on trial-averaged responses from a second partition. Varying the ridge penalty (*λ*) allowed us to find a continuum of decoding dimensions that trade off decoding accuracy versus encoding strength. Analysis was repeated when decoding the derivative of force. (**A**) Force decode accuracy (*R*^2^) as a function of encoding strength. Encoding strength was defined as the fraction of population variance explained by the decode dimension. Each point corresponds to a unique value of *λ*. For small values of *λ*, decode accuracy was high for both M1 populations and the motor units. As *λ* grew, so did encoding strength. For the M1 populations, decode accuracy declined almost immediately; it was not possible to find a dimension that both provided an accurate decode and captured considerable variance. In contrast, for the motor units, decode accuracy remained high across a range of *λ*, including values where encoding strength was quite high. Decode accuracy eventually declined for very high values of *λ*, simply because very strong regularization encouraged all weights to shrink towards zero. Colored x’s indicate encoding strength when decoding accuracy was still at 80% of its maximum, which forms the basis for analysis in the next two panels. (**B**) To estimate the most neural variance that a force-decode dimension can capture, we measured force encoding strength at 80% of maximum decoding accuracy. This value is low for the M1 populations (blue) but high for motor units (red). (**C**) As in (B), but the variance explained by the decode dimension is normalized by the variance explained by the first PC of population activity. For the motor units, the value near one reflects the fact that the first principal component closely mirrors force. Values are lower (*≈* 0.4) for the M1 population, but are not tiny. The force-decoding dimension captures almost half as much variance as the first PC, which by definition captures the most variance of any dimension. Thus, force is a sizable signal in relative terms; the fact that it captures so little overall response variance is due to the high-dimensionality of the data, as expected given the proposal in Fig. 1A. (**D**) Same analysis as (A) except decoders were trained to predict the derivative of force (equivalent to the velocity of the Pac-Man cursor). Decoding accuracy was never high for the motor units. It was for the M1 populations, but accuracy declined swiftly at higher values of *λ*. (**E**) Same as (B) except when decoding the derivative of force. (**F**) Same as (C) except when decoding the derivative of force.

**Supplementary Figure 17.**
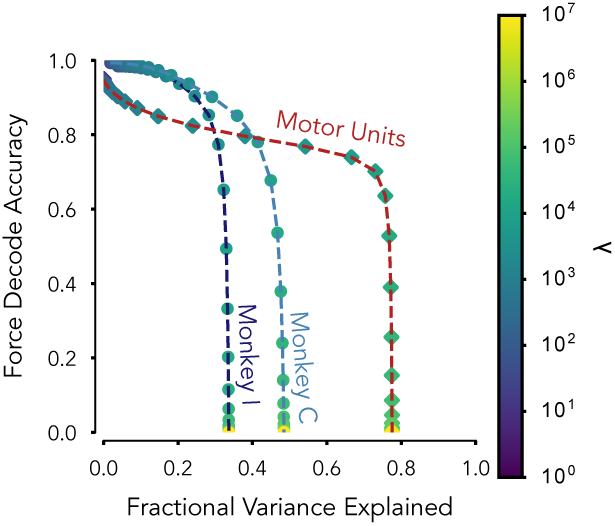
After restricting analysis to M1 neurons with the strongest force tuning, force still explains less variance than for the motor unit population. A reasonable hypothesis is that there might exist a sizable subset of M1 neurons that encode force in a ‘pure’ fashion, as do motor units. Put differently, there might be a subset of neurons whose tuning is consistently aligned with the readout dimension that provides descending commands. Those neurons would consistently encode force, regardless of the currently engaged subskill. In contrast, the hypothesis in Fig. 1E supposes that descending commands are driven by different factors at different moments. Individual-neuron tuning reflects mixtures of those factors. Thus, pure force-tuning will be rare, and become rarer still when activity is examined across multiple putative subskills. To explore, we down-selected M1 populations to include only the 134 M1 neurons whose firing rates showed the highest absolute correlation with force. We chose 134 to match the size of the motor unit population, and because it was *∼*10% of the the full M1 population. If there is a sizable subset of pure force-encoding neurons in M1, down-selection should restrict analysis to that subset, which should then behave much as the motor units. After down-selection, we repeated the decoding analysis described in Supp. Fig. 16 above. Plotting conventions are as in that figure. As expected, down-selected M1 populations had decoding dimensions that captured a larger fraction of population variance, relative to the full population. This can be seen in the plot by noting that decoding accuracy (vertical axis) remains high for larger values of variance explained (horizontal axis), relative to the original analysis in Supp. Fig. 16A. Yet even in the down-selected M1 population, the maximum proportion of population variance that could be explained (before decoding accuracy suffered) was only about half that for the motorunit population. These results agree with our observation that the vast majority of neurons have activity that does not mirror force. Even when responses have a component that reflects force – such that force can be decoded – the force component is still mixed with other signals. Thus, down-selection enriches the strength of force encoding, but does not identify a subpopulation with pure motor-unit-like force encoding.

**Supplementary Figure 18.**
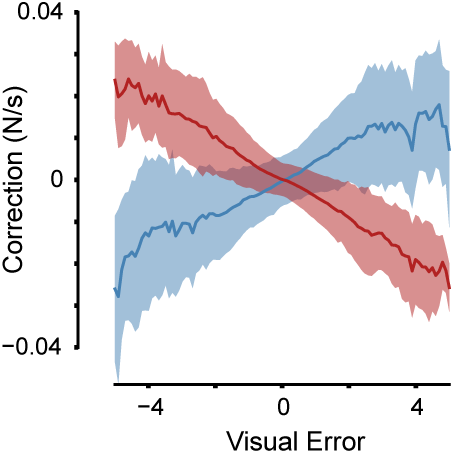
Feedback corrections during normal and inverted Pac-Man for monkey I. Analysis parallels that for monkey C in Fig. 6A. Traces plot the average change in the derivative of force relative to baseline, 150 ms in the future, as a function of current visual error. Baseline was simply the mean derivative for the condition to which that trial belonged. Analysis collapses across all trials. Envelopes show standard deviations. Visual error is defined as the difference between Pac-Man’s height and the height of the left-most portion of the dot-path. Red and blue traces correspond to normal and inverted subskills.

**Supplementary Figure 19.**
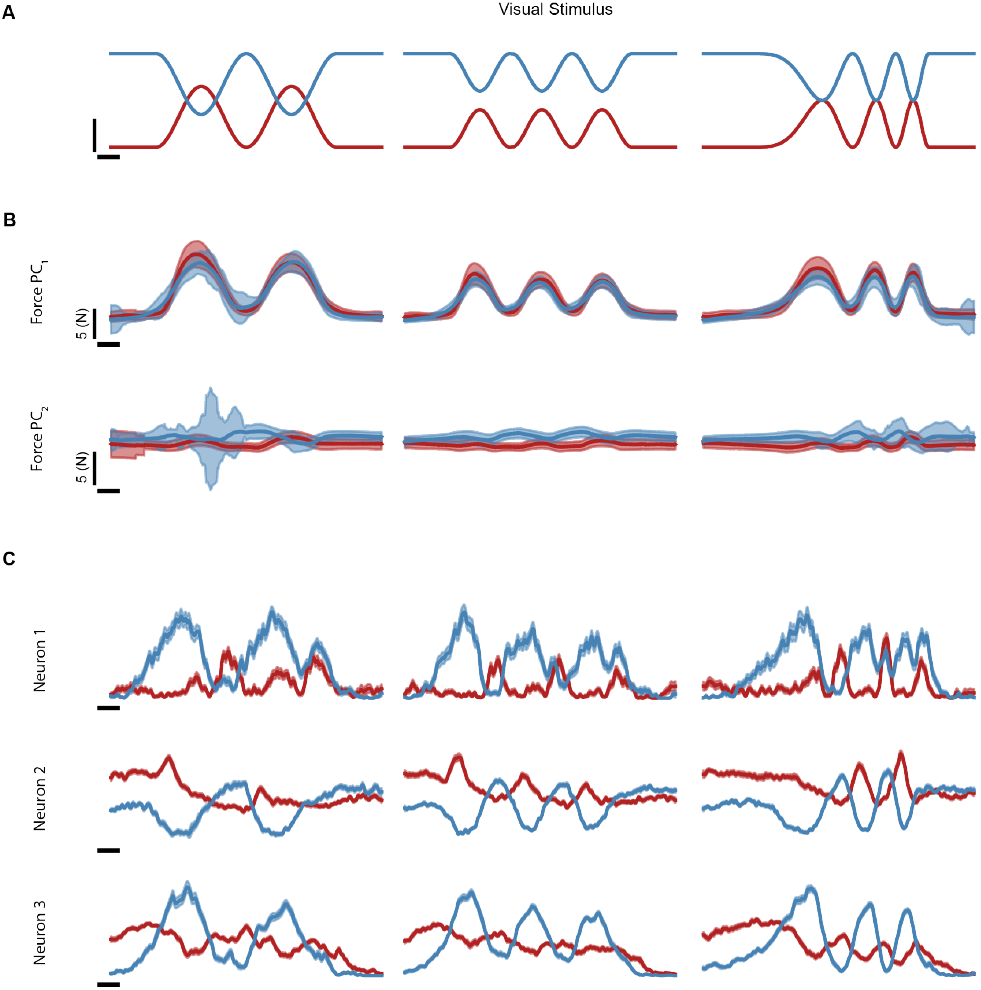
Further analysis of behavior and additional example neural responses as monkey C performed normal (red) and inverted (blue) versions of the Pac-Man task. Analysis complements that in Fig. 6. Monkey C was trained on this two-subskill version of the task following the main experiments. Because this task is challenging and takes considerable time to learn, monkey C was trained on only a few conditions. (**A**) Traces show the visual dot paths, for both subskills, during three conditions. Within each condition, the different dot paths instructed the same force profile for both subskills. (**B**) As intended, and as shown in Fig. 6, similar forces were produced during both subskills. Here we examine that match by considering all three degrees of freedom measured by the 3-axis load cell. Doing so is important because it is conceivable that force could be well-matched only in the forward direction, and not in ‘off axis’ dimensions. We projected three-dimensional forces onto their first two PCs and computed the mean (solid line) and SD (envelopes) for each condition. The first PC is dominated by forward force, which is (as the task requires) well-matched across subskills. The second PC captures off-axis forces. These were small and differed only slightly between subskills. Thus, the two subskills involved very similar time-varying profiles of generated force, not only in the forward direction but overall. (**C**) Additional example responses from three neurons, complementing the example in Fig. 6B. Traces show the mean firing rate with SEM (envelopes). A ‘pure’ force-encoding neuron would show identical responses across subskills. This was observed only occasionally. These examples illustrate that neurons sometimes showed partially inverted responses, yet it was not the case that neural responses simply reflected the visual dot path. A variety of response changes, including amplitude and phase shifts, were observed. Examples for monkey I are shown in Supp. Fig. 20.

**Supplementary Figure 20.**
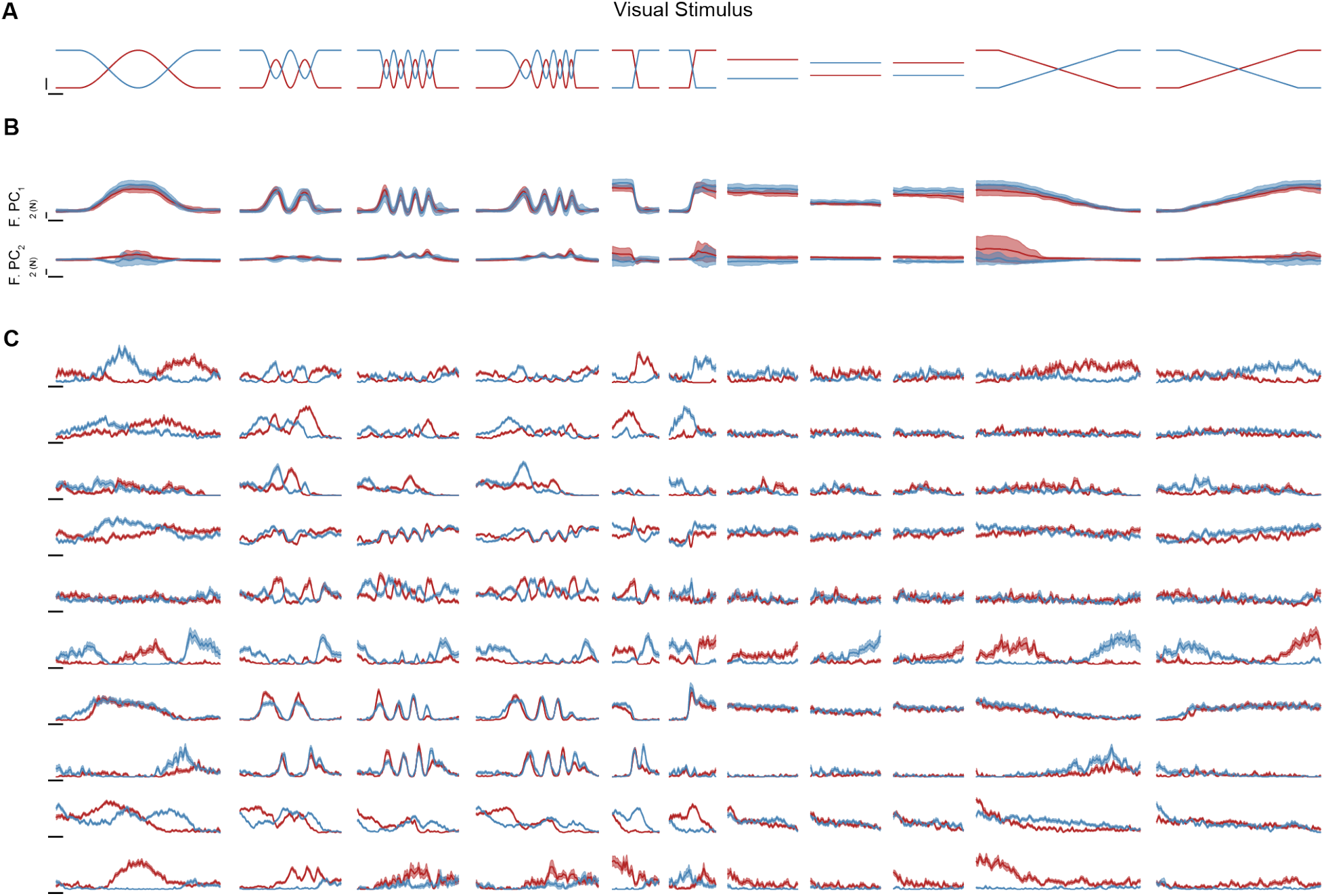
Behavior and example neural responses as monkey I performed normal (red) and inverted (blue) versions of the Pac-Man task. Analysis is similar to that for monkey C in Supp. Fig. 19 above. Monkey I was trained on the two-subskill version of the task before the main experiments began, and became adept at doing so across a broad variety of conditions before recordings began. Hence the larger number of conditions shown here. Correspondingly, we also show a larger number of single-neuron examples. (**A**) Traces show the visual dot paths, for both subskills, for all conditions. Within each condition, the different dot paths instructed the same force profile. (**B**) As in Supp. Fig. 19, we examined the match in force by considering all three degrees of freedom. We projected three-dimensional forces onto their first two PCs and computed the mean (solid line) and standard deviation (envelopes) for each condition. The first PC principally captures forward force, which is (as the task requires) well-matched across subskills. The second PC captures off-axis forces. These were small and, for most conditions, differed only slightly between subskills. However, this was somewhat variable across conditions. Some conditions (especially the slow and fast falling ramps) involved off-axis forces for one subskill but not the other. A reasonable concern is that differences in neural responses across subskills might (trivially) reflect these differences in off-axis forces, rather than reflecting subskill per se. Fortunately, there were multiple conditions (e.g. the faster sinusoids) that had a nearly perfect match for both forward and off-axis forces. As can be seen via inspection of the next panel, neural responses often differed across subskills, and this was true across all conditions. Thus, the main source of such differences cannot be the occasional differences in off-axis forces. (**C**) Example neural responses. Traces show mean firing rate. Envelopes show SEMs. A wide variety of response profiles was observed. Occasionally, a neuron had consistently similar firing rates for both subskills (see third and fourth rows from bottom). Occasionally a neuron’s response mostly inverted across conditions (see top row). Yet neither property was typical; responses typically differed in idiosyncratic ways that varied across conditions.

**Supplementary Figure 21.**
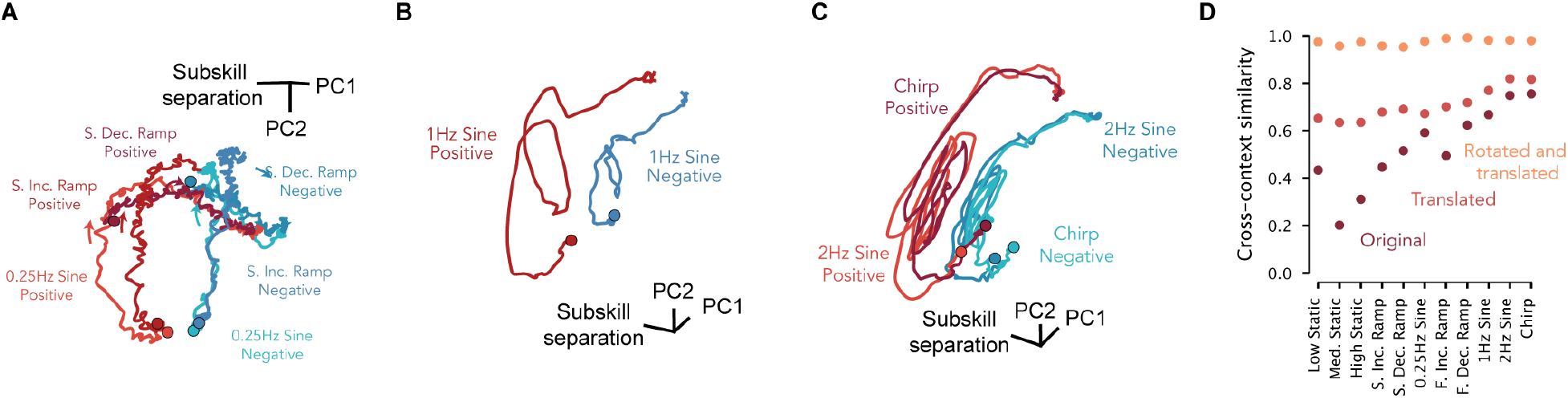
Neural trajectories for normal and inverted subskills, for monkey I. Analysis parallels that for monkey C in Fig. 6C,D. Monkey I performed more conditions, and the degree to which neural activity reflected subskill was condition-dependent. At lower frequencies (e.g. the statics and the slow ramps), across-subskill differences in neural activity were quite large (indeed, larger than for monkey C). Across-subskill differences were smaller (but still sizable) for high-frequency forces. A speculation is that this may reflect the fact that slowly-changing forces rely heavily on closed-loop control, which is necessarily very different between normal and inverted subskills (the same error demands opposing responses). This may be less relevant when high frequencies demand open-loop (e.g. template-based) strategies. (**A**) State-space trajectories for both subskills (red and blue) during three low-frequency force profiles. Two axes were found via PCA (applied to these conditions only). The third axis was selected to capture the mean difference in neural activity between subskills (also for these conditions only). The two subskills involved similarly shaped neural trajectories. However, despite their similar shape, the trajectories were not the same. Trajectories occurred in different regions of state space, as evidence by the horizontal shift between red and blue traces. It was also the case (though it is hard to appreciate in three dimensions) that trajectories unfolded in somewhat different dimensions. (**B**) Same but for activity during the 1 Hz force profiles. (**B**) Same but for activity during 2 Hz and Chirp force profiles. (**D**) Same analysis as in Fig. 6D, but for Monkey I. Across-subskill differences in neural trajectories involved both a rotation and translation. As noted above, differences in neural trajectories between subskills were larger at lower frequencies. Once the translation and rotation were removed, trajectories were very similar.

**Supplementary Figure 22.**
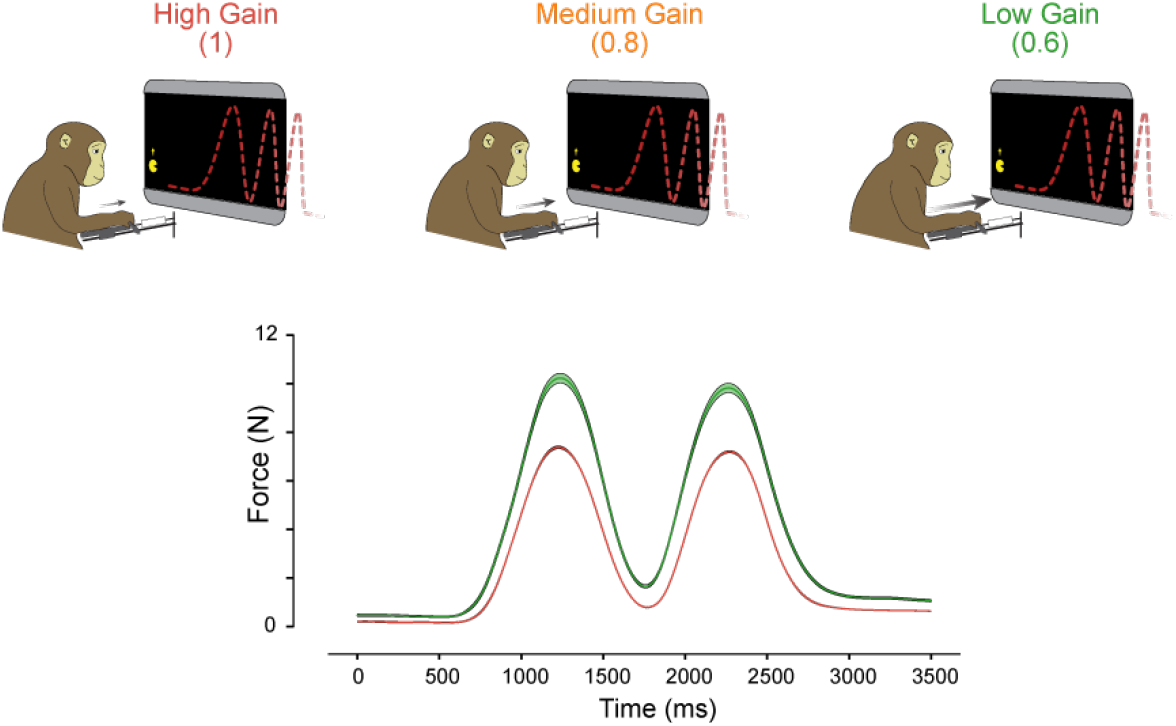
Illustration of the task and behavior used to probe neural activity when force output must scale to compensate for changes in gain. Data are for monkey C. This experiment is intended (in part) as a control for the finding that M1 activity strongly reflects subskill when comparing normal and inverted versions of the task. The forces generated in normal and inverted Pac-Man were very similar, but small differences often remained. Might seeming subskill-specific aspects of activity instead reflect imperfectly matched forces? A natural control is to intentionally create larger differences in force, within subskill. Under the flexible-repertoire hypothesis, neural activity is predicted to change more as a result of changes in subskill than as a result of within-subskill changes in motor output. Because this prediction – activity is dominated by subskill – is fundamental to the flexible-repertoire hypothesis, testing it also provides an additional test of that hypotheses. To do so, we intentionally created sizable differences in force by altering the gain between the load-cell and the height of the Pac-Man cursor (top row). Gain was stepped down at discrete moments during the session and reached a low value of 0.6, such that 67% more force was needed to drive Pac-Man to the same height. Our expectation was that altering gain would produce rapid adaptation of the existing subskill, rather than learning of a new one. Consistent with this expectation, adaptation to the gain change was nearly instantaneous, and was largely complete even within the first trial. Forces during a gain of 0.6 were thus a scaled version of those during a gain of 1 (bottom panel, envelopes show SE). Under the flexible-repertoire hypothesis, only a very modest proportion of the population response reflects motor output, but a sizable proportion reflects subskill. Neural activity should thus differ less between gain changes than between normal versus inverted subskills, even though motor output changes more. This was indeed the case, as is documented below (Supp. Fig. 23, Supp. Fig. 24).

**Supplementary Figure 23.**
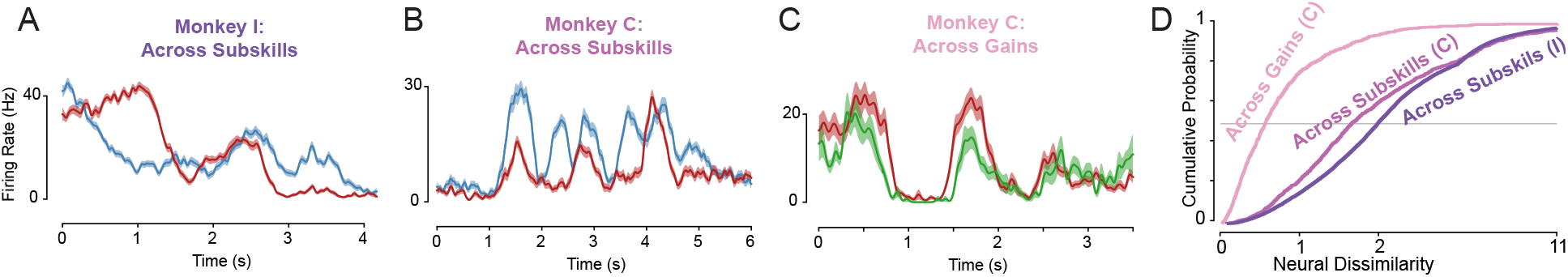
Single-neuron responses differed much more across subskills (normal versus inverted) than across gains. Each example neuron was chosen to be representative of response differences for that experiment. Specifically, the neuron was chosen so that the normalized magnitude of the response difference was near the median. (**A**) Average firing rate (*±* SE) of a neuron recorded from monkey I during the Chirp Condition, for normal (red) and inverted (blue) subskills. (**B**) Same for a neuron recorded from monkey C during the 0.5 Hz sine (this panel repeated from the main text). (**C**) Same for a neuron recorded from monkey C when gain was high (red) and low (green). (**D**) Cumulative Density Function (CDF) of neural dissimilarity scores for monkey C (magenta) and I (purple) during the normal and inverted subskills, and for monkey C during the multi-gain experiment (gain of 1 versus 0.6). Dissimilarity was measured once for each neuron and force profile. Dissimilarity was higher across subskills, even though changes in motor output were smaller.

**Supplementary Figure 24.**
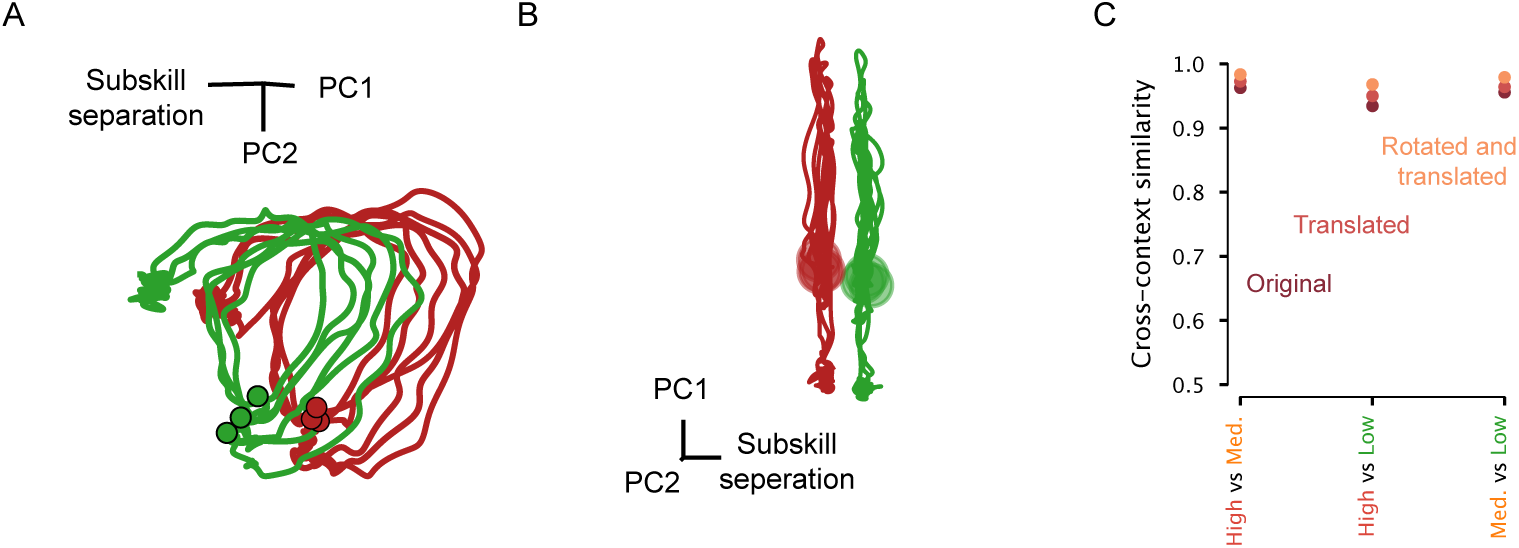
The population response changed only modestly across changes in gain. This analysis parallels the analysis across subskills in Fig. 6C,D and Supp. Fig. 21. (**A**) The population response during a gain of 1 (red) and 0.6 (green) for all three conditions (different traces). (**B**) Same but from a different viewing angle, chosen specifically to maximize separation between the two gains. Separation was present but small. (**C**) Cross-context similarity of the population response, across gains, under different similarity maximizing transformations: no transformation (maroon), translation (red-orange), rotation and translation (orange). Similarity was already high before the transformations, and became only slightly higher after them.

**Supplementary Movie 1. Video of the Pac-Man task being performed, and of motor-unit activity**. The top panel is a black-and-white video of the screen, taken from inside the rig, as the Pac-Man task was performed by monkey C. The video was shot from above, hence the perspective. Video is from a session were multiple motor units were recorded. Motor-unit spikes appear in the bottom panel, which reproduces the experimenter’s view of the recording system (Blackrock Microsystems), located outside the rig. The audio picks up on two sets of sounds: the spikes of a small handful of motor units (played on a loudspeaker outside the rig) and sound of the solenoid that controls juice delivery (also located outside the rig). Motor-unit spikes become more plentiful (and more motor-units begin to spike) in direct proportion to force, which is also reflected in the height of the Pac-Man icon. Yellow labels give a (rough) guess as to the behavioral strategy potentially used at different movements. Many of the features picked up on in the quantitative analyses of behavior, including small adjustments when forces changes slowly and swift commitment when force changes swiftly, can be seen by inspection.

## References

1. Krakauer, J. W., Hadjiosif, A. M., Xu, J., Wong, A. L. & Haith, A. M. Motor learning. Comprehensive Physiology 9, 613–663 (2019).

2. Sadtler, P. T. et al. Neural constraints on learning. Nature 512, 423–426 (2014).

3. Golub, M. D. et al. Learning by neural reassociation. Nature Neuroscience 21, 607–616 (2018).

4. Gallego, J. A. et al. Cortical population activity within a preserved neural manifold underlies multiple motor behaviors. Nature Communications 9, 4233 (2018).

5. Kakei, S., Hoffman, D. S. & Strick, P. L. Muscle and movement representations in the primary motor cortex. Science 285, 2136–2139 (1999).

6. Sergio, L. E., Hamel-Pâquet, C. & Kalaska, J. F. Motor cortex neural correlates of output kinematics and kinetics during isometric-force and arm-reaching tasks. Journal of Neurophysiology 94, 2353– 2378 (2005).

7. Sussillo, D. Neural circuits as computational dynamical systems. Current Opinion in Neurobiology 25, 156–163 (2014).

8. Vyas, S., Golub, M. D., Sussillo, D. & Shenoy, K. V. Computation through neural population dynamics. Annual Review of Neuroscience 43, 249–275 (2020).

9. Churchland, M. M. & Shenoy, K. V. Preparatory activity and the expansive null-space. Nature Reviews Neuroscience 25, 213–236 (2024).

10. Yang, G. R., Joglekar, M. R., Song, H. F., Newsome, W. T. & Wang, X.-J. Task representations in neural networks trained to perform many cognitive tasks. Nature Neuroscience 22, 297–306 (2019).

11. Marton, C. D., Lajoie, G. & Rajan, K. Efficient and robust multi-task learning in the brain with modular task primitives. arXiv (2021). 2105.14108.

12. Goudar, V., Peysakhovich, B., Freedman, D. J., Buffalo, E. A. & Wang, X.-J. Schema formation in a neural population subspace underlies learning-to-learn in flexible sensorimotor problem-solving. Nature Neuroscience 26, 879–890 (2023).

13. Logiaco, L., Abbott, L. & Escola, S. Thalamic control of cortical dynamics in a model of flexible motor sequencing. Cell reports 35, 109090–109090 (2021).

14. Versteeg, C. & Miller, L. Dynamical feedback control: motor cortex as an optimal feedback controller based on neural dynamics. Preprint at 10.20944/preprints202201 (2022).

15. Chiappa, A. S., An, B., Simos, M., Li, C. & Mathis, A. Arnold: a generalist muscle transformer policy. arXiv (2025). 2508.18066.

16. DePasquale, B., Sussillo, D., Abbott, L. & Churchland, M. M. The centrality of population-level factors to network computation is demonstrated by a versatile approach for training spiking networks. Neuron (2023).

17. Riveland, R. & Pouget, A. Natural language instructions induce compositional generalization in net-works of neurons. Nature Neuroscience 27, 988–999 (2024).

18. Gallego, J. A., Perich, M. G., Miller, L. E. & Solla, S. A. Neural manifolds for the control of movement. Neuron 94, 978–984 (2017).

19. Churchland, M. M. et al. Neural population dynamics during reaching. Nature 487, 51 (2012).

20. Oby, E. R. et al. Dynamical constraints on neural population activity. Nature Neuroscience 28, 383– 393 (2025).

21. Genkin, M., Shenoy, K. V., Chandrasekaran, C. & Engel, T. A. The dynamics and geometry of choice in the premotor cortex. Nature 645, 168–176 (2025).

22. Russo, A. A. et al. Motor cortex embeds muscle-like commands in an untangled population response. Neuron 97, 953–966.e8 (2018).

23. Saxena, S., Russo, A. A., Cunningham, J. P. & Churchland, M. M. Motor cortex activity across movement speeds is predicted by network-level strategies for generating muscle activity. bioRxiv 2021.02.01.429168 (2021).

24. Perich, M. G., Narain, D. & Gallego, J. A. A neural manifold view of the brain. Nature Neuroscience 28, 1582–1597 (2025).

25. Kaufman, M. T., Churchland, M. M., Ryu, S. I. & Shenoy, K. V. Cortical activity in the null space: permitting preparation without movement. Nature Neuroscience 17, 440–448 (2014).

26. Oby, E. R. et al. New neural activity patterns emerge with long-term learning. Proceedings of the National Academy of Sciences 116, 15210–15215 (2019).

27. Perkins, S. M., Amematsro, E. A., Cunningham, J. P., Wang, Q. & Churchland, M. M. An emerging view of neural geometry in motor cortex supports high-performance decoding (2024).

28. Marshall, N. J. et al. Flexible neural control of motor units. Nature Neuroscience 1–13 (2022).

29. Elsayed, G. F., Lara, A. H., Kaufman, M. T., Churchland, M. M. & Cunningham, J. P. Reorganization between preparatory and movement population responses in motor cortex. Nature communications 7, 13239 (2016).

30. Miri, A. et al. Behaviorally selective engagement of short-latency effector pathways by motor cortex. Neuron 95, 683–696.e11 (2017).

31. Rouse, A. G. & Schieber, M. H. Condition-dependent neural dimensions progressively shift during reach to grasp. Cell Reports 25, 3158–3168.e3 (2018).

32. Harpaz, N. K., Ungarish, D., Hatsopoulos, N. G. & Flash, T. Movement decomposition in the primary motor cortex. Cerebral Cortex 29, 1619–1633 (2019).

33. Ames, K. C. & Churchland, M. M. Motor cortex signals for each arm are mixed across hemispheres and neurons yet partitioned within the population response. eLife 8, e46159 (2019).

34. Willett, F. R. et al. Hand knob area of premotor cortex represents the whole body in a compositional way. Cell 181, 396–409.e26 (2020).

35. Warriner, C. L., Fageiry, S., Saxena, S., Costa, R. M. & Miri, A. Motor cortical influence relies on task-specific activity covariation. Cell Reports 40, 111427 (2022).

36. Schroeder, K. E., Perkins, S. M., Wang, Q. & Churchland, M. M. Cortical control of virtual self-motion using task-specific subspaces. The Journal of Neuroscience 42, 220–239 (2022).

37. Sabatini, D. A. & Kaufman, M. T. Reach-dependent reorientation of rotational dynamics in motor cortex. Nature Communications 15, 7007 (2024).

38. Kim, J.-H., Daie, K. & Li, N. A combinatorial neural code for long-term motor memory. Nature 637, 663–672 (2025).

39. Trautmann, E. M. et al. Large-scale high-density brain-wide neural recording in nonhuman primates. bioRxiv 2023.02.01.526664 (2023).

40. Ghavampour, A. et al. A paradigm to study the learning of muscle activity patterns outside of the natural repertoire. Journal of Neurophysiology 134, 347–360 (2025).

41. Churchland, M. M. & Shenoy, K. V. Temporal complexity and heterogeneity of single-neuron activity in premotor and motor cortex. Journal of Neurophysiology 97, 4235–4257 (2007).

42. Seely, J. S. et al. Tensor analysis reveals distinct population structure that parallels the different computational roles of areas m1 and v1. PLOS Computational Biology 12, e1005164 (2016).

43. Pellegrino, A., Stein, H. & Cayco-Gajic, N. A. Dimensionality reduction beyond neural subspaces with slice tensor component analysis. Nature Neuroscience 27, 1199–1210 (2024).

44. Evarts, E. V. Relation of pyramidal tract activity to force exerted during voluntary movement. Journal of Neurophysiology 31, 14–27 (1968).

45. Scott, S. H., Gribble, P. L., Graham, K. M. & Cabel, D. W. Dissociation between hand motion and population vectors from neural activity in motor cortex. Nature 413, 161–165 (2001).

46. Ajemian, R. et al. Assessing the function of motor cortex: Single-neuron models of how neural response is modulated by limb biomechanics. Neuron 58, 414–428 (2008).

47. Zimnik, A. J. et al. Identifying interpretable latent factors with sparse component analysis. bioRxiv 2024.02.05.578988 (2024).

48. Katlowitz, K. A., Picardo, M. A. & Long, M. A. Stable sequential activity underlying the maintenance of a precisely executed skilled behavior. Neuron 98, 1133–1140.e3 (2018).

49. Sussillo, D., Churchland, M. M., Kaufman, M. T. & Shenoy, K. V. A neural network that finds a naturalistic solution for the production of muscle activity. Nature Neuroscience 18, 1025–1033 (2015).

50. Yang, C. S., Cowan, N. J. & Haith, A. M. De novo learning versus adaptation of continuous control in a manual tracking task. eLife 10, e62578 (2021).

51. Lara, A. H., Elsayed, G. F., Zimnik, A. J., Cunningham, J. P. & Churchland, M. M. Conservation of preparatory neural events in monkey motor cortex regardless of how movement is initiated. eLife 7, e31826 (2018).

52. DeWolf, T. et al. Neuro-musculoskeletal modeling reveals muscle-level neural dynamics of adaptive learning in sensorimotor cortex. bioRxiv 2024.09.11.612513 (2025).

53. Driscoll, L. N., Shenoy, K. & Sussillo, D. Flexible multitask computation in recurrent networks utilizes shared dynamical motifs. Nature Neuroscience 27, 1349–1363 (2024).

54. Gribble, P. L. & Scott, S. H. Overlap of internal models in motor cortex for mechanical loads during reaching. Nature 417, 938–941 (2002).

55. Briggman, K. & Jr., W. K. Multifunctional pattern-generating circuits. Annual Review of Neuroscience 31, 271–294 (2008).

56. Stroud, J. P., Porter, M. A., Hennequin, G. & Vogels, T. P. Motor primitives in space and time via targeted gain modulation in cortical networks. Nature neuroscience 21, 1774–1783 (2018).

57. Tian, L. Y. et al. Neural representation of action symbols in primate frontal cortex. bioRxiv 2025.03.03.641276 (2025).

58. Shenoy, K. V., Sahani, M. & Churchland, M. M. Cortical control of arm movements: A dynamical systems perspective. Neuroscience 36, 337–359 (2012).

59. Sussillo, D. & Barak, O. Opening the black box: Low-dimensional dynamics in high-dimensional recurrent neural networks. Neural Computation 25, 626–649 (2013).

60. Remington, E. D., Egger, S. W., Narain, D., Wang, J. & Jazayeri, M. A dynamical systems perspective on flexible motor timing. Trends in Cognitive Sciences 22, 938–952 (2018).

61. Churchland, M. M., Yu, B. M., Sahani, M. & Shenoy, K. V. Techniques for extracting single-trial activity patterns from large-scale neural recordings. Current Opinion in Neurobiology 17, 609–618 (2007).

62. Cunningham, J. P. & Yu, B. M. Dimensionality reduction for large-scale neural recordings. Nature Neuroscience 17, 1500–1509 (2014).

63. Langdon, C., Genkin, M. & Engel, T. A. A unifying perspective on neural manifolds and circuits for cognition. Nature Reviews Neuroscience 24, 363–377 (2023).

64. Todorov, E. & Jordan, M. I. Optimal feedback control as a theory of motor coordination. Nature Neuroscience 5, 1226–1235 (2002).

65. Scott, S. H. Optimal feedback control and the neural basis of volitional motor control. Nature Reviews Neuroscience 5, 532–545 (2004).

66. Pruszynski, J. A. & Scott, S. H. Optimal feedback control and the long-latency stretch response. Experimental Brain Research 218, 341–359 (2012).

67. Pruszynski, J. A. et al. Primary motor cortex underlies multi-joint integration for fast feedback control. Nature 478, 387–390 (2011).

68. Pruszynski, J. A., Kurtzer, I. & Scott, S. H. Rapid motor responses are appropriately tuned to the metrics of a visuospatial task. Journal of Neurophysiology 100, 224–238 (2008).

69. Kalidindi, H. T. et al. Rotational dynamics in motor cortex are consistent with a feedback controller. eLife 10, e67256 (2021).

70. Mastrogiuseppe, F. & Ostojic, S. Linking connectivity, dynamics and computations in low-rank recurrent neural networks. arXiv 99, 609–623.e29 (2017). 1711.09672.

71. Sun, X. et al. Skill-specific changes in cortical preparatory activity during motor learning. bioRxiv 2020.01.30.919894 (2020).

72. Sheahan, H. R., Franklin, D. W. & Wolpert, D. M. Motor planning, not execution, separates motor memories. Neuron 92, 773–779 (2016).

73. Heald, J. B., Lengyel, M. & Wolpert, D. M. Contextual inference underlies the learning of sensorimotor repertoires. Nature 600, 489–493 (2021).

74. Cesanek, E., Zhang, Z., Ingram, J. N., Wolpert, D. M. & Flanagan, J. R. Motor memories of object dynamics are categorically organized. eLife 10, e71627 (2021).

75. Pezzulo, G. & Cisek, P. Navigating the affordance landscape: Feedback control as a process model of behavior and cognition. Trends in Cognitive Sciences 20, 414–424 (2016).

76. Wasicko, R. J., McRuer, D. T. & Magdaleno, R. E. HUMAN PILOT DYNAMIC RESPONSE IN SINGLELOOP SYSTEMS WITH COMPENSATORY AND PURSUIT DISPLAYS (1966).

77. Lara, A. H., Cunningham, J. P. & Churchland, M. M. Different population dynamics in the supplementary motor area and motor cortex during reaching. Nature Communications 9, 2754 (2018).

78. Schmid, P. J. Dynamic mode decomposition of numerical and experimental data. Journal of Fluid Mechanics 656, 5–28 (2010).

